# Predicting coexistence in experimental ecological communities

**DOI:** 10.1101/598326

**Authors:** Daniel S. Maynard, Zachary R. Miller, Stefano Allesina

## Abstract

The study of experimental communities is fundamental to the development of ecology. Yet, for most ecological systems, the number of experiments required to build, model, or analyze the community vastly exceeds what is feasible using current methods. Here, we address this challenge by presenting a statistical approach that uses the results of a limited number of experiments to predict the outcomes (coexistence and species abundances) of all possible assemblages that can be formed from a given pool of species. Using three well-studied experimental systems—encompassing plants, protists, and algae with grazers—we show that this method predicts with high accuracy the results of unobserved experiments, while making no assumptions about the dynamics of the systems. These results suggest a fundamentally different study design for building and quantifying experimental systems, requiring a small number of experiments relative to traditional approaches. By providing a scalable method for navigating large systems, this work provides an efficient way to study highly diverse experimental communities.

## Introduction

Ecologists have long used small experimental systems to explore and test fundamental ecological phenomena: Gause’s experiments^1^ illustrated and popularized the principle of competitive exclusion; chemostat-based systems highlighted the need for eco-evolutionary modeling^2^; stage-structured populations grown in laboratory conditions offered a clear example of chaotic dynamics^3^; and yeast communities have provided key insight into ecological tipping points^4^.

Despite these advances, experimenting with more diverse ecological communities has proven exceedingly difficult for several reasons. First, a pool of *n* species can give rise to as many as 2*^n^ −* 1 distinct species assemblages (i.e., combinations of presence/absence of species). Even for moderate *n*, these are too many assemblages to handle experimentally. Second, rarely will all possible assemblages lead to coexistence; often, large communities “collapse” to smaller subsets of species due to extinctions (e.g., a two-species community collapsing to a monoculture). Finally, two replicates of the same assemblage might lead to different outcomes—for example, due to multistability or stochasticity— complicating efforts to characterize the dynamics of the system.

These challenges have been raised in numerous studies that experimented with speciose communities^5–8^. A common approach is to seed plots with single species (monocultures), pairs of species, and several larger assemblages, often chosen at random^9^. Due to the number of species considered, however, the space of possible assemblages is so large that only a small fraction of assemblages can be tested. For example, the “Biodiversity II” experiment included 18 different plant species, from which 168 of the 2^18^*−* 1 = 262, 143 possible species assemblages were planted, with their dynamics tracked for decades^10^. Ideally, one would use a principled method to choose which assemblages to test, so as to maximize the probability of coexistence across assemblages (and therefore not “waste” experiments), while ensuring that the selected subset of experiments allows for robust inference across the full set of unobserved assemblages.

The goal of this work is to address these challenges by developing a method in which a limited number of observations are used to predict coexistence for all 2*^n^ −* 1 assemblages that can be formed from a pool of *n* species. For any combination of species in the pool, our method attempts to determine whether the species can coexist, and if so at what abundances. As a starting point, we require that a limited number of experiments is conducted, using a variety of assemblages, and that species abundances are measured at a single (final) time point in each experimental community. These modest requirements make this method ideally suited for studying speciose communities. While this approach can be applied to any ecological system, it will likely be most useful in experimental settings where stochasticity and exogenous factors can be minimized.

The approach is summarized in Box 1 and Fig. 1, and explained in detail in the Methods. By recording the abundances of species grown in different combinations (a snapshot of abundances that we refer to as a dynamical “endpoint”), we parameterize a matrix *B* that encodes the abundances of species in all possible combinations—including those not observed experimentally. Armed with this estimate, we can predict the outcomes of unobserved experiments: for each possible species assemblage, our method returns the probability that the species will coexist, and, if so, it predicts the abundance of each species in the assemblage. We test this approach using data from three published studies, showing that it can predict out-of-fit experimental outcomes with high accuracy. We then explore the flexibility of this method by simulating data from a variety of nonlinear, non-equilibrium, and non-pairwise dynamical models of species interactions. Lastly, we show that the number of experiments needed to fit this model scales linearly with the number of species, requiring on the order of *n* experiments to predict the outcomes of all 2*^n^ −* 1 possible assemblages.

**Figure 1.**
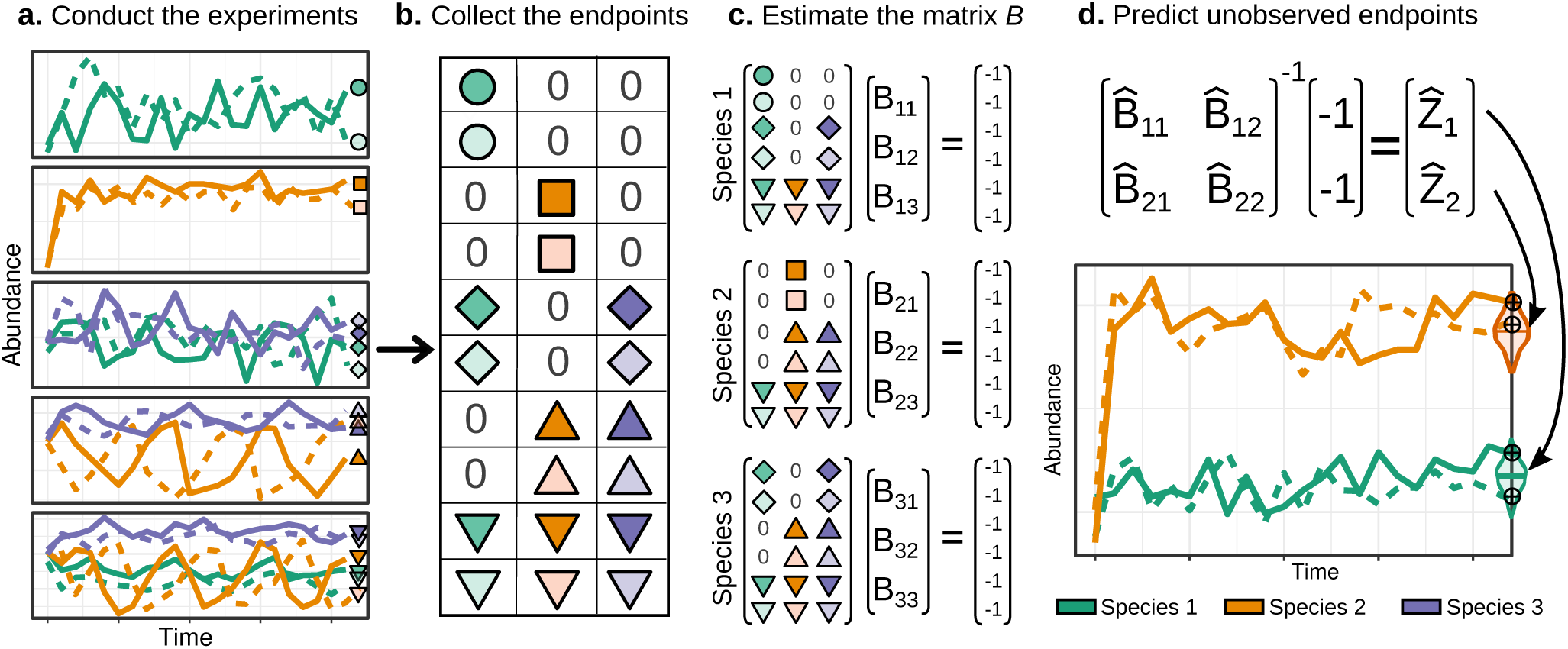
Predicting unobserved experimental outcomes. (a) A set of experiments is conducted, with each assemblage varying in initial composition and initial abundances. Colors denote different species and line types denote each of two replicates. The experiments are run for a sufficiently long time, at which point a single snapshot is taken of each species’ abundance in each replicate. (b) These abundances are collected into a matrix, with columns corresponding to species and rows corresponding to endpoints. Here, shapes represent species’ endpoint abundances for each of two replicates across five different assemblages. (c) The matrix *B*, which defines the full set of endpoints, is estimated by fitting a separate hyperplane through each species’ set of endpoints. (d) To predict the endpoint of an unobserved assemblage—here, the combination of species 1 and 2—we take the corresponding sub-matrix of *B*, invert it, and compute the negative row sums. To analyze the three empirical datasets, we implement a Bayesian approach to estimating *B*, resulting in a posterior distribution for the unobserved endpoint abundances (shown by the violin plots).

### BOX 1 - Approach

#### Goal and applications

Given a pool of *n* species, our method aims to predict the outcomes of all 2*^n^ −*1 possible assemblages. For each assemblage, we predict whether the species can coexist and, if so, at what abundances.

This method can aid experimentalists in a variety of settings. For example, when studying the relationship between diversity and productivity, one could conduct a pilot study using as few as 2*n* plots (e.g., all monocultures and all leave-one-out communities), and then use the method to predict productivity in all other possible assemblages. Similarly, the method could be used to identify candidate assemblages with maximal productivity (for example, for the development of biofuels). Alternatively, the method can be used as a road map to build large experimental communities in which all species coexist—a challenging feat when selecting assemblages at random ^7,22, 23^.

#### Input

The input for the method is a list of empirical “endpoints”, each recording the abundance of species in one sub-community at the end of the experiment. Given a sufficiently large and diverse set of endpoints (Methods), one can use these data to predict coexistence for all unobserved assemblages. In particular, one needs to start with a variety of assemblages that differ in species composition or initial densities. Each assemblage is allowed to follow its dynamical trajectory until a pre-determined time or condition is reached (e.g., optical density [chlorophyll content] stabilizes, when dealing with bacteria [algae]), at which point the abundances of all extant species in the community are recorded (Fig. 1). These endpoint measurements are then organized into a matrix, with columns corresponding to species and rows to communities (Fig. 1).

#### Model

To start with a simple, extensible model, we take the endpoint abundance of each species *i* in each endpoint assemblage *k* to be a linear function of the endpoint abundances of the other species present in the final assemblage:

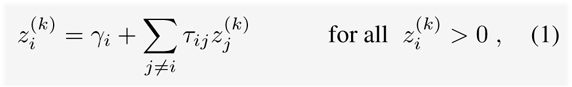

where 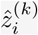 is the centroid (or “average”) endpoint abundance of species *i* in assemblage *k*; *γ_i_* models the abundance of species *i* when growing alone; and *τ_ij_* is the percapita effect that species *j* has, if present, on the *endpoint abundance* of species *i*. Ecologically, we expect *γ_i_ >* 0 for producers and *γ_i_ ≤* 0 for consumers and predators. Eqn. 1 can be rewritten more concisely as (Methods):

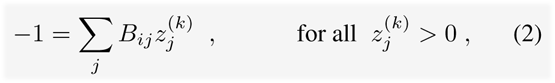

where *B_ii_* = *−*1*/γ_i_* and *B_ij_* = *τ_ij_/γ_i_* for *i ≠ j*. These coefficients are then collected into the *n × n* matrix *B*.

The empirically measured endpoint abundances for assemblage *k* are denoted *x*^(*k*)^, which are assumed to be a noisy estimate of the “true” endpoint abundance *z*^(*k*)^. See Methods for details.

#### Parameter Estimation

In theory, estimating the entries of *B* amounts to performing *n* linear regressions (Fig. 1c), effectively fitting a collection of *n* hyperplanes through the measured endpoints ^29^. However, this straight-forward approach ignores the error structure inherent in the data (Supplementary Information). We thus develop and implement a Bayesian regression approach that accounts for measurement error in each 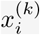 (Methods). This yields posterior distributions for the matrix *B* and for the estimated endpoint abundances 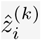.

#### Prediction

Under the model given by Eqn. 1, the matrix *B* encodes all possible endpoints of the system. Thus, with an estimate of *B* in hand, we can determine whether any particular set of species can coexist by taking a matrix inverse (Fig. 1d and Methods). For example, to predict the outcome of the unobserved combination of Species 1 and 2 in Fig. 1d, we take the corresponding 2 *×* 2 sub-matrix of *B*, invert it, and compute the negative row sums. The resulting values are the predicted endpoint abundances for this two-species community. If any abundances are negative, this indicates that these species cannot coexist. If all abundances are positive, then these species may coexist (however, the endpoint might be dynamically unstable, and therefore unreachable experimentally) (Methods). The probability of coexistence for unobserved or out-of-fit assemblages is calculated as the proportion of posterior *B* matrices that result in the coexistence of a given assemblage.

#### Assumptions

This method is best suited to settings where exogenous factors (e.g., dispersal or immigration) and stochasticity can be minimized. It is therefore ideal for controlled laboratory or microcosm studies, but may require higher replication in field-based systems to overcome experimental noise.

Although our fitting procedure assumes that endpoints are related in a linear fashion (Eqn. 1), it does not assume that the community *dynamics* are linear or additive (Extended Data 1-6, Supplementary Information). The method does, however, require that dynamics play out long enough to ensure that species have time to go extinct, thereby preventing “false positive” coexistence (Extended Data 8, Supplementary Information).

Finally, when endpoint abundances are not believed to be linearly related, Eqn. 1 can be modified to include additional terms (e.g., higher-order interactions), and the same fitting approach may be applied without requiring substantially more experiments.

## Results

### Empirical systems

To demonstrate this method, we first analyze published data of three experimental systems. Although these studies were not specifically designed to test our method, each study reports the final abundances of all species in each experimental assemblage, and each contains a sufficient number of unique assemblages to benchmark the method by making out-of-fit predictions.

First, we consider data from Kuebbing *et al.*^11^, who conducted growth experiments using two phylogenetically paired sets of old-field plants. Both sets contain four species, drawn from the families Asteraceae, Fabaceae, Lamiaceae, and Poaceae. In one set, all species are native to the study site; in the other, all species are non-native. For both the native and non-native pools, experimental communities were initialized with 14 out of 15 (= 2^4^ *−* 1) possible assemblages, with each combination replicated 10 times. Dry-weight aboveground biomass for each species in each assemblage was recorded after 112 days of growth^11^.

Second, we analyze data from Rakowski and Cardinale^12^, who grew consumer-resource communities consisting of five species of green algae and two herbivorous species from the family Daphniidae (*Ceriodaphnia dubia* and *Daphnia pulex*). Here we focus on the four algal species that survived in a sufficient number of endpoints to test our method: *Chlorella sorokiniana*, *Scenedesmus acuminatus*, *Monoraphidium minutum*, and *Monoraphidium arcuatum*. The algae were grown in all four-species combinations with high replication, and one of the two herbivores was added to two-thirds of the replicates. The communities were incubated for 28 days, during which time each assemblage collapsed to a subset of the initial pool, generating a variety of distinct endpoints.

Third, we turn to time-series data published by Pennekamp *et al.*^13^, who studied six species of ciliates growing on bacterized medium, capturing 53 of the 63 (= 2^6^ *−* 1) possible assemblages. We omit one species that declined in abundance across the time series (*Tetrahymena thermophila*), leading to the five-species subsystem used here: *Colpidium striatum*, *Dexiostoma campylum*, *Loxocephalus* sp., *Paramecium caudatum*, and *Spirostomum teres*. Each experimental assemblage was replicated twice, and each species’ density in each assemblage was tracked for 57 days^14^. Experiments were repeated at six different temperatures ranging from 15 to 25 *^◦^*C.

These three studies encompass 10 distinct experimental systems: native and non-native plant communities, two different herbivore-algae sub-communities, and six temperature conditions for protists. For each experiment, species abundances for all assemblages were reported, providing a sufficient diversity of endpoints to fit our model and test the accuracy on out-of-fit assemblages using a “jackknife” leave-one-out approach (Methods, Supplementary Information). Specifically, for every system we omitted each of the *k* assemblages in turn, using the remaining *k −* 1 assemblages to fit the model and predict the abundances of all species in the omitted community. We then compared our predictions with what was observed experimentally.

In addition, because the plant experiments included all but one of the possible combinations of species, we were able to implement a more rigorous *k*-fold cross-validation approach. We used a subset of endpoints to parametrize the model and predict the endpoint abundances in the omitted communities; we repeated this procedure for all combinations of endpoints sufficient to fit the model, ranging from 1 to 9 omitted communities (Supplementary Information).

### Abundance patterns

As illustrated in Fig. 2, our method predicts the endpoints of each system with high accuracy. Across all assemblages, the medians of the posterior distributions generated by our method closely track the median observed abundances. Not only does our method achieve high accuracy for within-fit data (Figs. 2a, c, e); it also accurately estimates the out-of-fit endpoints and captures the observed variability in abundances (Figs. 2b, d, f).

**Figure 2.**
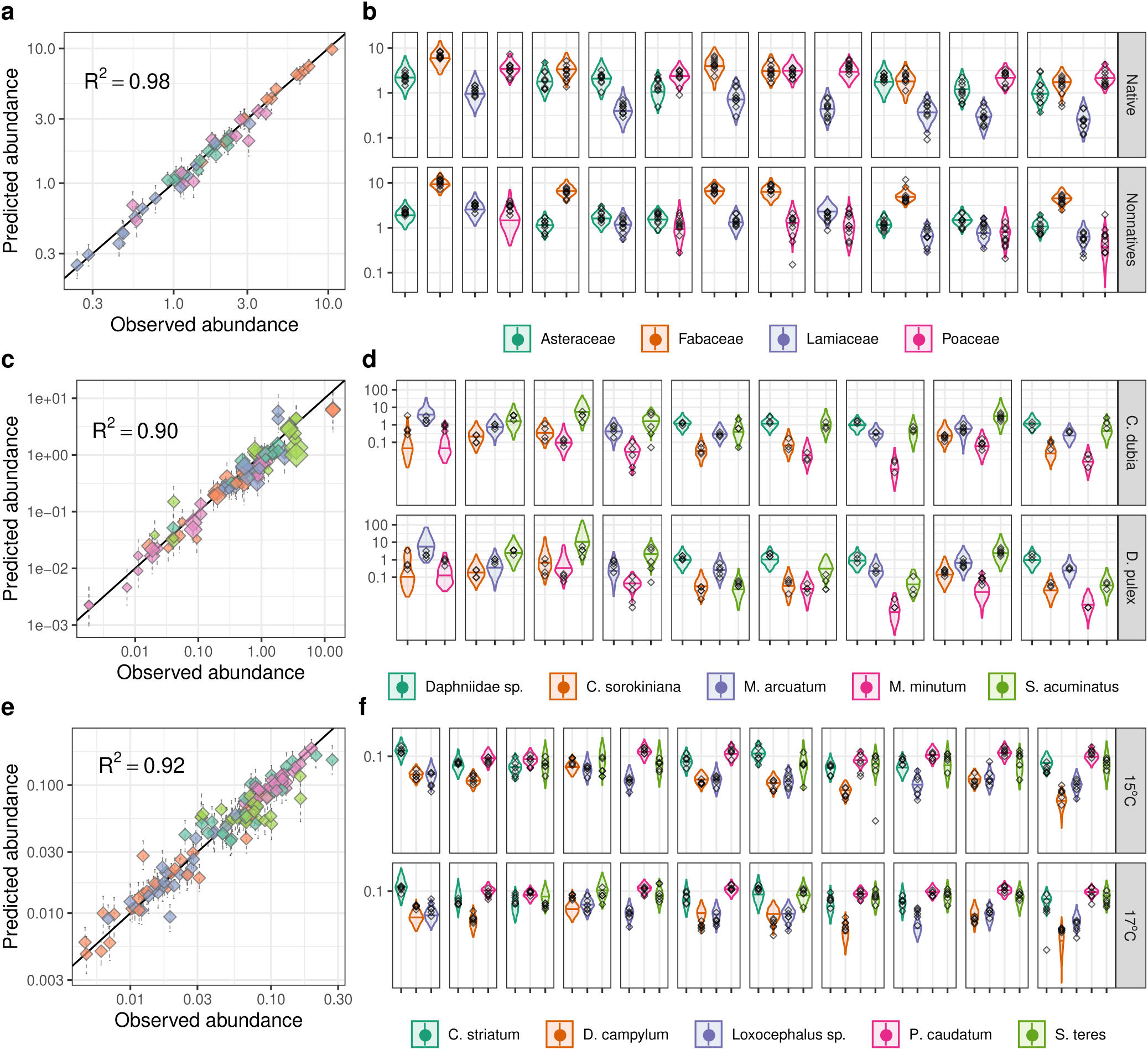
Results for three experimental systems. (a, b) The old-field plant system comprising native and non-native communities; (c, d) the herbivore-algae system with two different Daphniidae consumers; (e, f) the ciliated protist system, assayed under 6 different temperature conditions, with 15*^◦^* and 17*^◦^* C shown here. (a, c, e) The observed vs. predicted abundances for each experimental system, with vertical error bars denoting the 95% prediction interval of the posterior distribution; the size of the diamonds reflects the number of replicates per measurement. (b, d, f) The out-of-fit predicted abundances for each assemblage, using a “jackknife” leave-one-out approach. Colors correspond to species, and each box corresponds to a different assemblage. The small black diamonds show the measured abundance of each species in each replicate for each assemblage. Violin plots show the 95% prediction intervals for the posterior distribution of the abundance, with the horizontal bars indicating the median predicted abundance. Note that only assemblages containing more than two replicates and species are shown in d and f; see Supplementary Information for the complete results.

By tallying the proportion of posterior draws of *B* that result in coexistence (Methods), we can also quantify the probability that an unobserved or out-of-fit assemblage will go extinct. For example, in both algae systems, our results suggest that the unobserved two-species community comprising *C. sorokiniana* and *M. minutum* has a high probability of extinction, regardless of the identity of the herbivore (Supplementary Figures 5-6). Our findings also suggest that the four-species protist system comprised of *C. striatum*,*D. campylum*, *P. caudatum*, and *S. teres* has a non-zero risk of extinction at 19, 21, and 23° C, driven in part by high variation and relatively low abundance of *D. campylum* which causes the posterior abundance distribution to overlap zero (Supplementary Figures 7-12). Verifying these predictions would require additional experimentation; yet such information can help identify which unobserved assemblages one might avoid in future experiments to minimize the risk of extinctions (see Study Design, below).

For the plant data, a *k*-fold cross validation approach demonstrates that only a few experimental endpoints are sufficient to predict abundances in all 14 assemblages with high accuracy (Fig. 5, Supplementary Information). However, the choice of experiments used to fit the model (i.e., the experimental design) is critical, as discussed in detail in the Study Design section.

Lastly, for the protist system we have access to the full time series describing the dynamics, allowing us to compare the quality-of-fit of our method against traditional methods based on timeseries data. The estimates obtained using only endpoint data compare favorably to those obtained using standard trajectory matching to fit a Generalized Lotka-Volterra dynamical model to the full time series (Supplementary Information). Yet, by foregoing the need to model the dynamics, our method predicts species abundances with higher accuracy while requiring a fraction of the data.

### Structure of the matrix B

Because our approach does not attempt to model the dynamics of these systems, the matrix *B* has no direct mechanistic interpretation—it simply encodes the patterns in the endpoint abundances (Methods). We also reiterate that the experimental studies used here were not designed to test our method, such that their use is intended only to illustrate this method. Yet, by exploring the structure of *B*, we can demonstrate that our estimates are internally consistent and align with the known biology of the experimental systems, verifying that our method captures meaningful patterns in the data and is not over-fitting the endpoints (Figs. 3-4, Supplementary Information).

For example, we find that the coefficients of *B*, which summarize the effect of one species’ end-point abundance on another’s, are phylogenetically conserved between the native and non-native plant data sets (Fig. 3a). In the herbivore-algae system, the identity of the herbivore has little effect on interactions among the algae, suggesting weak or non-existent herbivore-mediated higher-order effects (Fig. 3b). And, in the protist system, the species-by-species coefficients exhibit consistent signs and magnitudes across temperatures, with many of these coefficients varying smoothly (Fig. 4).

**Figure 3.**
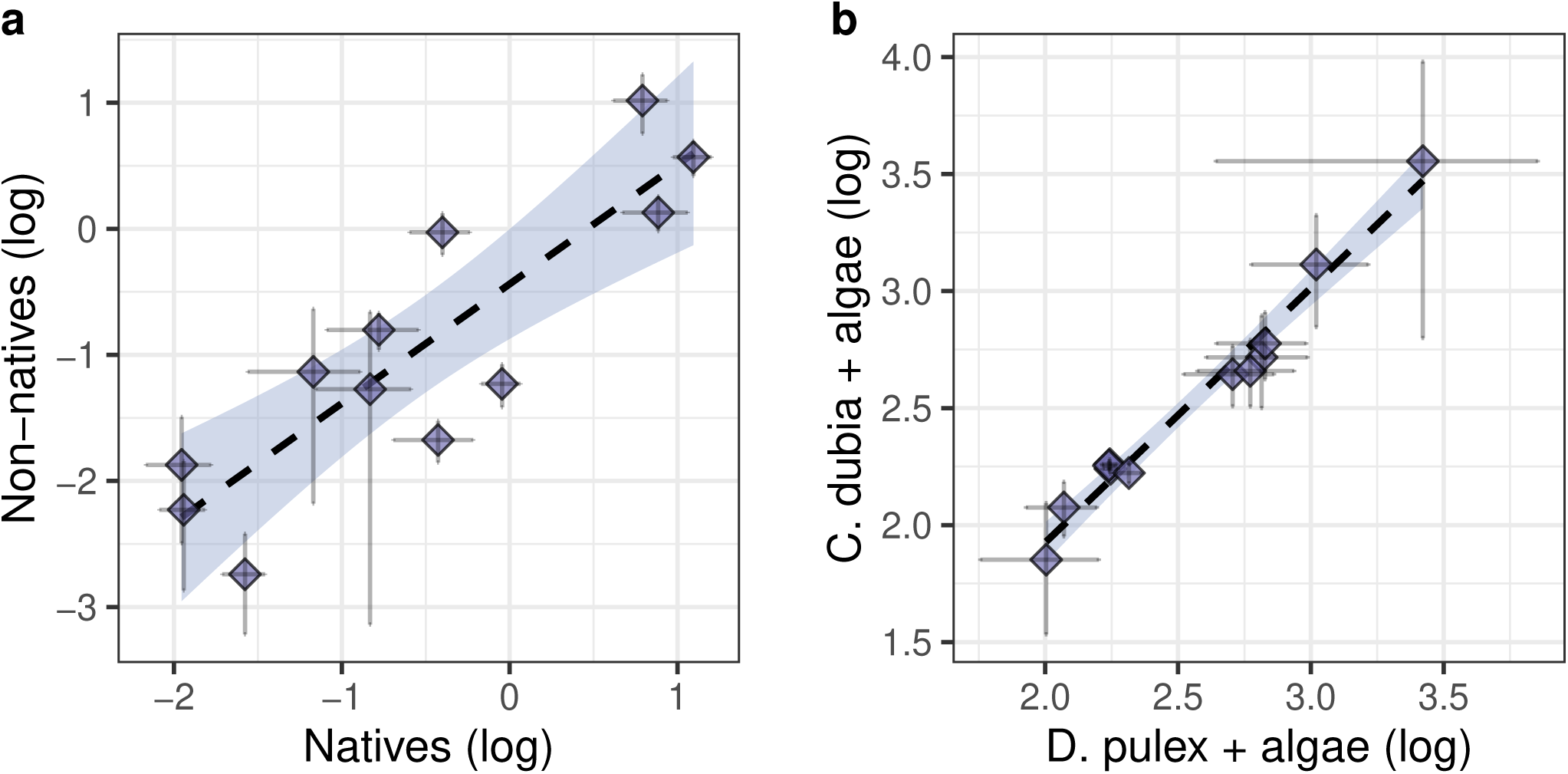
Comparison of the matrix *B* within the plant and herbivore-algae datasets. Internal consistency in the estimated *B* for each system highlights that the method is not over-fitting the data, and illustrates how this approach can offer basic ecological insight. (a) A plot of the log-transformed coefficients for the native (x-axis) vs. non-native (y-axis) plant systems, paired by family into Asteraceae, Fabaceae, Lamiaceae, and Poaceae, revealing that the effects of species from one family on species from another are conserved across the two systems. (b) A plot of the log-transformed algaeby-algae coefficients for the herbivore-algae system, estimated for the communities containing *D. Pulex* (x-axis) vs. those containing *C. dubia* (y-axis). The coefficients are strongly consistent across both herbivore systems, suggesting minimal herbivore-induced changes in algae-by-algae interactions (i.e., negligible higher-order effects on the endpoints). The dashed line gives the mean regression trend, with the blue shaded region showing 95% confidence bands. Horizontal and vertical bars show the posterior 95% prediction interval for each coefficient. The log transformation includes the addition of an offset to ensure all coefficients are positive; see Supplementary Information for details.

**Figure 4.**
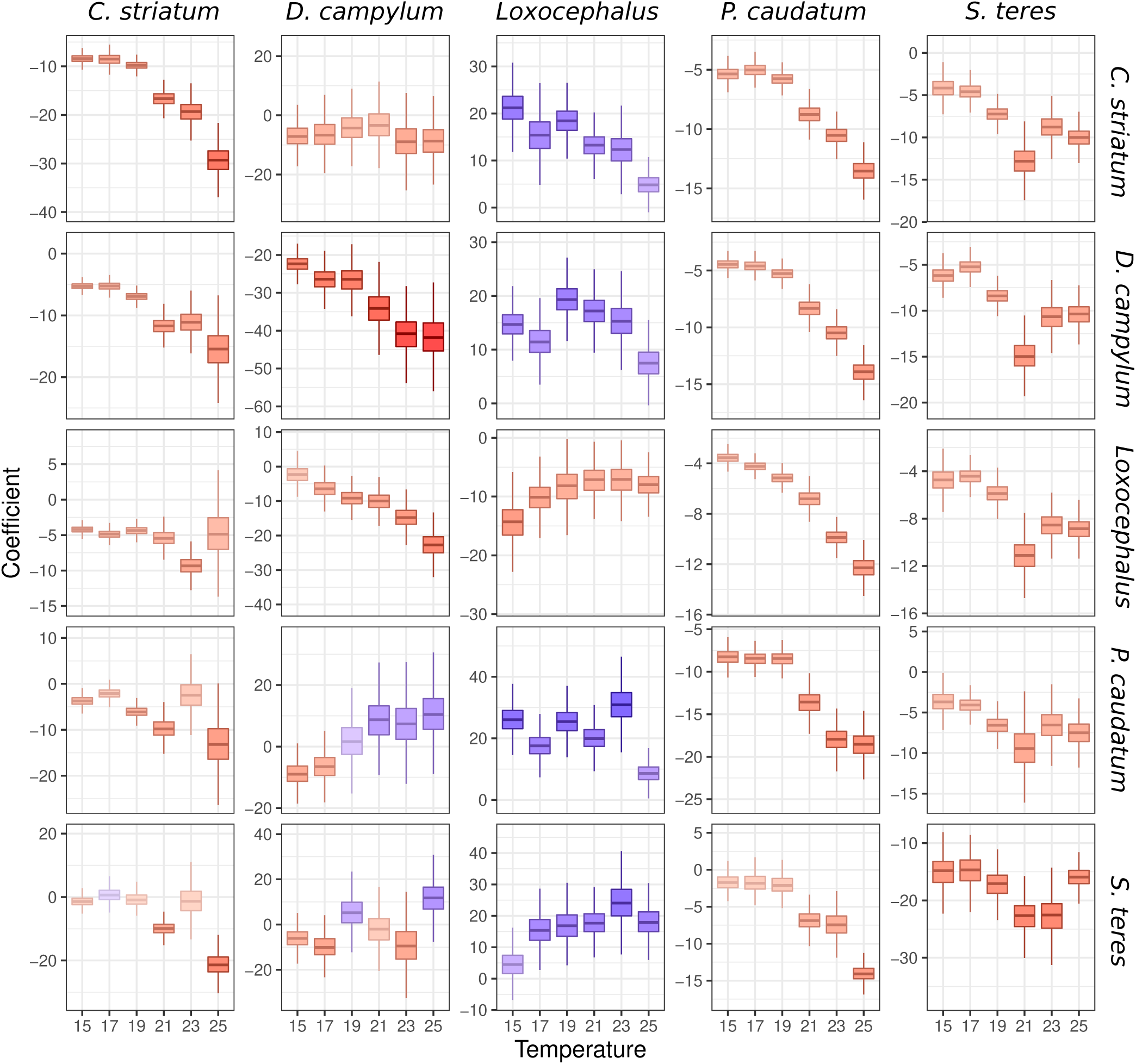
Comparison of the matrix *B* in the protist system across the six temperatures. Each panel depicts the change in the respective *B_ij_* coefficient across temperature treatments, organized with *B*_11_ in the upper left, down to *B*_55_ in the lower right. The boxplots within each panel report the medians and interquartile ranges for the posterior distribution of each coefficient at each temperature. Colors reflect the median coefficient value, with blue indicating positive effects and red indicating negative effects. Our method reveals generally consistent changes in these coefficients across the temperature gradient, further highlighting that this method is not over-fitting the data and is identifying a biologically reasonable pattern. Some coefficients exhibit non-linear fluctuations—such as the effect of *P. caudatum* on the abundance of *Loxocephalus* sp.—but these parameters exhibit consistent signs, similar magnitudes, and overlapping prediction intervals, indicative of statistical noise. Nevertheless, because our method is statistical in nature, we caution against any mechanistic interpretation of these coefficients; they are depicted here to illustrate the consistency of the model fit across different experimental conditions.

**Figure 5.**
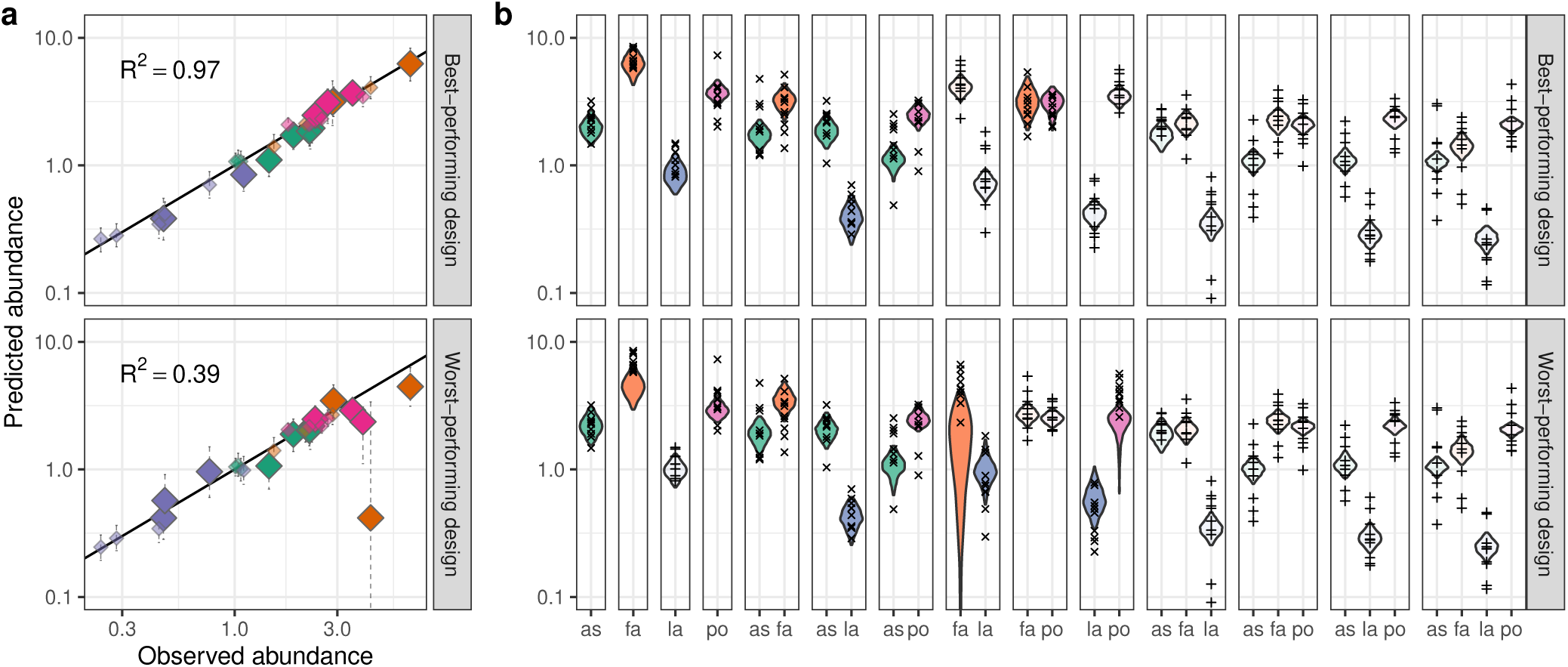
Predicting multiple endpoints out-of-fit. The best-performing (top) and worst-performing (bottom) experimental designs using 6 assemblages to fit the model and predict the remaining 8 assemblages. (a) Observed vs. predicted abundances for each experimental system, with vertical error bars denoting the 95% prediction interval of the posterior distribution. Points used to fit the model are in lighter shades and smaller size; larger points with solid color represent out-of-fit predictions. The *R*^2^ is calculated using only out-of-fit data. (b) Predicted abundances for each assemblage. Unshaded violin plots show the distribution of the predictions for in-fit data used to fit the model (+); violin plots with solid colors show the distributions of the out-of-fit data (*×*). The choice of which assemblages to use to fit the model (i.e., the experimental design) is crucial, as made clear by the difference in quality of fit between the worst (bottom) and best (top) designs making use of 6 assemblages. Similar results are found for the non-native plants, and when more assemblages are used to fit the model (Supplementary Figures 24-26). For larger communities, an efficient design is the choice of all monocultures, all ‘leave-one-out’ communities, and the full community (Extended Data 7).

These results demonstrate the type of ecological insight that can be gained, even without making mechanistic interpretations about the parameters. Moreover, the fact that these estimates of *B* are internally consistent—for example, largely consistent across changing temperature—highlights that this method is not over-fitting the data, in which case we would expect our estimates to fluctuate dramatically with small changes to the system.

### Simulations

To explore the robustness and generality of these findings, we fit simulated endpoint data generated from a variety of non-linear, non-equilibrium, and non-pairwise dynamical models, including Lotka-Volterra dynamics with limit cycles, competition with Allee effects^15^, facultative mutualism with saturation^16^, consumption with saturation^17^, and competition with high-order interactions^18^(Supplementary Information).

In all cases, our approach recovers the model endpoints quite accurately (Extended Data 1-6), demonstrating that, despite the complex and nonlinear dynamics, the endpoint structures of these systems are approximately additive and linear (as in Eqn. 1). The method also successfully predicts which combinations of species will be unable to coexist. For example, in the model with mutualism and saturation, our method correctly identifies which assemblages cannot coexist (Extended Data 4-5), and, for the Allee effect model, it correctly identifies the unstable fixed points even though they are never observed (Extended Data 3).

We intentionally selected dynamical systems that can exhibit oscillatory dynamics or alternative states—features that are difficult to reconcile with the assumption of a single “endpoint” (Eqn. 1). In these cases, our approach estimates the “average” of the distribution of abundances quite accurately (e.g., the centroid of the oscillations; Extended Data 2, 5). Certainly, the usefulness of this type of estimate will be context dependent. In many ecological applications, for example, we do not need to know the precise dynamics of a community; rather, all we want to know is whether a given set of species can coexist (i.e., if the centroid is far enough from zero to avoid stochastic extinctions), or what species’ average abundances will be across a landscape (e.g., for diversity-function or conservation questions). In such settings, our method can offer insight even though the predicted endpoint (the center of the oscillations) may never be observed experimentally.

Our method struggled most with systems containing strong higher-order interactions (HOIs), where the effect of one species on another varies depending on the abundance of a third species (Extended Data 6). These nonlinear relationships, however, can be easily incorporated into our method by including additional terms in Eqn. 1, such as interactions between pairs of species, 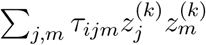, exactly as in standard polynomial regression models. In this way, exploring more complex models does not require more complex experimental designs or substantially more data^19, 20^. Thus, in settings where the baseline model performs poorly, different choices for Eqn. 1 can be tested, allowing one to adjust the study design accordingly and benchmark the model formulations using out-of-fit or *k*-fold cross validation.

### Study Design

This approach suggests new study designs for parameterizing community models and building large experimental systems. Because multispecies endpoints contribute to multiple equations used to estimate the matrix *B*, diverse communities contain more information about *B* than do small communities (e.g., the endpoint for the 3-species communities in Fig. 1b [triangles] appears in all three systems of equations shown in Fig. 1c). Leveraging this fact, our method provides an approach to experimental design that scales linearly with the size of the species pool.

Rather than growing species in monoculture and in all pairwise combinations^7, 8, 21–23^, one could first grow the full community of *n* species along with all ‘leave-one-out’ communities comprised of *n −* 1 species each. Using this approach, it is possible to estimate *B* using only *n* + 1 experiments (as opposed to *n*^2^*/*2). This design, however, is not robust in practice; these large communities contain little information about species-poor assemblages, potentially leading to substantial prediction error for smaller assemblages. Yet, by also measuring a selection of species-poor assemblages—for example, the *n* monocultures—one can efficiently “anchor” the model, providing high quality-of-fit using only 2*n* + 1 experiments.

To examine the practical impact of experimental design, we fit the model for both plant systems using six assemblages to predict the remaining eight endpoints out-of-fit. There are fifteen possible designs (combinations of assemblages) that make use of six endpoints and allow for parameterization of the model: in Fig. 5 we show the best- and worst-fitting designs for the native plant pool. The prediction accuracy shows a wide range, highlighting the importance of selecting a robust design. However, the best 6-assemblage design has a goodness-of-fit (*R*^2^) on par with that obtained using the full dataset (Fig. 2), despite using less than half of the endpoints. The same qualitative outcome is found for the invasive plant community. Further varying the number of endpoints used to fit the native and nonnative plant systems (from six to thirteen) reinforces this pattern: the best designs for any number of in-fit assemblages fare almost as well as the design using all experiments (Supplementary Information). However, across nearly 3000 k-fold cross validation assemblages, there is substantial variance in goodness-of-fit, with many designs performing well, and a few performing very poorly.

Using simulated endpoint data, we show that designs using a mix of species-rich and species-poor communities fare better than those using only small or large communities to fit the data (Extended Data 7). In particular, the design using all monocultures and all pairs of species performs poorly relative to random designs with the same number of experiments. By contrast, the design using all monocultures and all leave-one-out communities is among the best designs for any number of experiments.

In practice, an experimental design may also fail because some assemblages collapse to smaller communities, leading to fewer unique assemblages than desired. To address this challenge, we propose a simple, iterative scheme for experimental design: first, one conducts a minimal set of experiments sufficient to obtain a draft estimate of *B*. This matrix is then used to predict the outcomes of all unperformed experiments. The assemblages with the highest inferred probability of coexistence are selected for a second round of experiments, increasing the chances that these new communities yield useful data. One might repeat this process several times, updating the estimate of *B* after each iteration. At each stage, it is possible to compute the out-of-fit accuracy of the previous *B* matrix, giving a real-time measure of the model performance. If the quality-of-fit remains poor even after several iterations (or, for example, after annual sampling points in a biodiversity-ecosystem function experiment), then one can adjust the model assumptions (e.g., by adding HOIs or quadratic terms in Eqn. 1) or consider alternative approaches. By iteratively updating *B*, one maximizes the utility of each round of experiments, avoiding “wasted” experiments, while ensuring adequate model fit. This approach can help experimentalists navigate the enormous space of possible assemblages, providing an efficient way to explore, build, and quantify large experimental systems.

## Discussion and conclusions

From a small set of observed experimental assemblages, we show how to estimate a matrix *B*, which can be used to predict the outcomes of all possible assemblages. We have successfully applied this approach to three independent experimental systems, obtaining high-quality predictions for out-of-fit data, despite the fact that our method completely neglects functional responses, behavioral changes, spatial effects, resource depletion, etc.—all of which have been documented in these systems to varying degrees^24–28^. The simulation results further illustrate that nonlinear and non-equilibrium dynamics need not produce nonlinear endpoint structures. Together, these results demonstrate that a simple additive model and a small set of experiments can provide robust out-of-fit predictions for complex dynamical systems.

In applying this approach, an important practical consideration is deciding when to end an experiment and sample species abundances. In some cases, there may be a biologically-relevant time point (e.g., the end of the growing season for annual plants). In other settings, one might use some inexpensive, non-invasive technique to monitor the state of the community (e.g., optical density, chlorophyll fluorescence, or total respiration) and track this measure until it converges to a stationary distribution (as was done here for the protist experiment; Supplementary Information). In general, our results suggest that one simply needs to ensure that the dynamics have passed through any transients, allowing species sufficient time to go extinct. In the protist system, for example, the model fit was poor between days 1 and 10, but increased sharply in quality at day 11 and remained good for all days between 11 and 30 (Extended Data 8, Supplementary Information), highlighting that this method is not overly sensitive to the choice of when to end the experiment. In systems where the endpoints cannot be observed, or where there is no principled way to determine when to end the experiments—as in communities that cannot be easily manipulated or systems with dynamics that play out over very long time periods—alternative methods might be more suitable.

A key benefit of this approach is that the minimum number of required experiments scales linearly with the size of the species pool, helping to overcome a central challenge in studying large experimental systems. In theory, this method becomes increasingly efficient as the size of the species pool increases, with the minimum sufficient proportion of experiments scaling as (*n* + 1)*/*(2*^n^ −* 1).

In practice, however, large communities may present computational challenges, since the number of parameters to be estimated still scales with *n*^2^. In large experimental systems, it is also likely that some species will persist in fewer than *n* unique endpoints, preventing estimation of their associated parameters. Invoking strict Bayesian priors or employing more advanced search algorithms may help overcome some of these limitations. Nevertheless, improving the computational efficiency of this approach is a challenge, and one that would help extend this method to highly speciose systems.

Because this method requires relatively few experiments, it can easily be integrated into traditional studies (e.g., biodiversity-ecosystem function studies) without requiring a substantial overhaul of the experimental design. That is, we envision that this approach can be used to complement—rather than replace—current experimental methods. However, to ensure robust predictions, we emphasize that it may be necessary to use higher replication in field-based experiments to overcome stochasticity and experimental noise. Indeed, when external factors (including, especially, dispersal and recruitment) cannot be adequately controlled, this approach may not be appropriate. However, one advantage of our method is that it provides a straightforward and principled way to assess performance using out-of-fit predictions. Additionally, because this approach permits very efficient experimental designs, high replication may be more feasible.

Central to our method is that it uses a single measurement for each assemblage. By focusing exclusively on the abundances of species present in a final snapshot, this approach requires drastically fewer measurements than time-series methods. Yet this method is also completely blind to initial conditions: two experiments that start with different species compositions but collapse to the same assemblage are considered to be ‘replicates’ of this endpoint. Extending the model to account for initial conditions is an important next step, as doing so should help improve fit and prediction by incorporating knowledge about which assemblages have collapsed due to extinctions.

While many challenges remain in the study of speciose ecological communities, our framework provides a powerful approach for exploring these systems. We adopt a simplified, statistical method that robustly predicts the outcomes of unobserved experiments using relatively few observations. By foregoing dynamical modeling, we gain tractability, thereby providing a principled and efficient means for studying, navigating, and building diverse experimental systems.

## Methods

### Experimental setting

We consider a pool of *n* species, which can give rise to as many as 2*^n^ −* 1 distinct combinations of species’ presence/absence. For any combination (henceforth, *assemblage*), we consider an experiment in which each of the selected species is inoculated at some initial density in controlled conditions, and the dynamics of the system (i.e., inter- and intraspecific interactions) are allowed to play out. Once sufficient time has elapsed, the abundance of every species is measured. As explained in the main text, several methods could potentially inform us on when to sample the abundances, and in Supplementary Information we show the consequences of sampling during the transient phase. Here, we simply assume that the system has settled in some dynamical attractor, including the possibility of stochastic fluctuations around an equilibrium, deterministic chaos, periodic oscillations, etc. This experimental design permits destructive sampling methods (e.g., high-throughput sequencing of the endophytic microbial assemblage in plants), and does not require any observation of the transient dynamics or initial conditions.

We refer to the set of abundance measurements for each assemblage as an “endpoint” of the dynamics, denoted by *x*^(^*^k^*^)^, where *k* is an index referring to the set of species present at non-zero abundance in the endpoint measurement. Note that there is not a bijective mapping between assemblages and endpoints in general. For example, a particular assemblage of species may not coexist, in which case the system will collapse to a subset of species and reach the same endpoint as if it had been seeded with only the sub-assemblage. Conversely, identical initial assemblages may result in different endpoints. Here, we distinguish between two cases: first, the endpoints might be sampled from the same attractor, but may differ because of cycling or sampling error; second, we might have “true multi-stability”^29^—the replicate systems have reached distinct attractors, possibly depending on the initial abundance of each species. In this work, we do not explicitly account for the latter case, assuming that replicate endpoints are drawn from the same stationary distribution (as such, multistability is conflated with “sampling error”). Instances of true multi-stability in ecological dynamics are well-documented, but should be identifiable in data with sufficient replication^30–32^.

Given a set of experimental endpoints, we attempt to predict which unobserved assemblages can coexist and, if so, at what endpoint abundances. To accomplish this, we assume that the end points are related by a simple linear model. As noted in Box 1, this model takes the form:

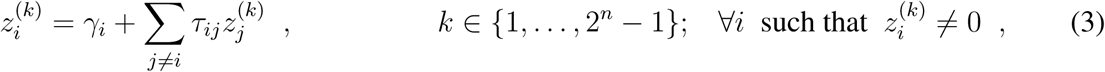

where *k* denotes the assemblage (as above) and *z_i_* denotes the average (across replicates) endpoint abundance of species *i*. In cases where the stationary distribution is not a fixed point, 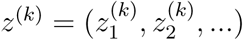 can be thought of as the “centroid” of the attractor for assemblage *k*. The coefficients *γ* and *τ* are statistically-determined constants that relate the endpoint abundances to each other. Intuitively, *γ_i_* models the average abundance of species *i* when grown in isolation, and therefore we expect *γ_i_* to be positive for producers—reflecting their carrying capacity—and zero (or negative) for consumers and predators. The *τ_ij_* can be seen as the average per-capita effect that members of species *j* have on the endpoint abundance of species *i*. We assume each *τ_ij_* is constant across communities (i.e., *τ_ij_* has no dependence on *k*).

Manipulating Eqn. 3 slightly, we can obtain

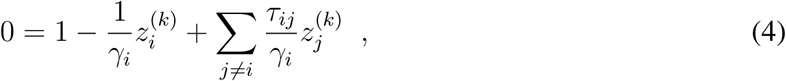

and, letting 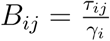 for *i ≠ j* and 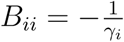,

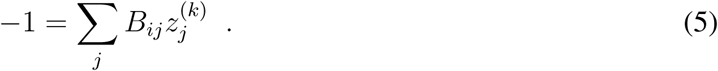

Eqn. 5 can be written more compactly in matrix form as

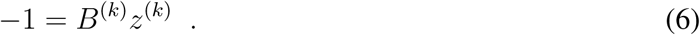

Here, if the *k*^th^ endpoint contains *w* species, then the left-hand side of Eqn. 6 is an *w × w* vector of 1s, and *B*^(^*^k^*^)^ is the *w × w* submatrix obtained by selecting only the elements of *B* (the full *n × n* matrix of coefficients) whose rows and columns correspond to species that have nonzero density in endpoint *k*.

Having estimated the matrix *B*, it is possible to predict any of the 2*^n^ −* 1 endpoints by solving Eqn. 6 for a desired assemblage *k*. If the estimated solution, 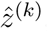, contains any negative elements, we interpret this as an indication that the species in assemblage *k* cannot coexist. If every element of 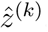 is positive, these elements give the endpoint abundances at which the species may coexist. This condition (non-negativity, or “feasibility”) is necessary for the coexistence of community *k*, but not sufficient, because the endpoint (or the attractor associated with it) may not be attractive or stable.

### Inferring *B* from a set of endpoints

We are primarily concerned with the practical challenge of inferring the matrix *B* from endpoint data. As a first approach, we form equations relating the elements of *B* and the endpoint abundances, and use these to solve for *B* on a row-by-row (species-by-species) basis. Although this “näıve” procedure ignores the complex error structure of the model (Supplementary Information), it is illustrative of the general approach, helping to provide intuition.

To implement this first approach, we introduce the matrices *E_i_*, which contain the observed end-point abundances *x*^(^*^k^*^)^, for any *k ∈ {*1*,…,* 2*^n^ −* 1*}* such that 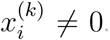. That is, each *E_i_* contains all the endpoints in which species *i* is present. Every *E_i_* has *n* columns, corresponding to the *n* species, and we fill in a zero wherever a species is not present in an endpoint. The number of rows of *E_i_* will be variable, depending on the dataset.

As a simple example, consider a case in which we have a pool of three species, and species 1 is present in 5 endpoints: two endpoints containing only species 1 (monocultures), two endpoints in which species 1 and 2 coexist, and one in which species 1 and 3 coexist. The structure of matrix *E*_1_ would be:

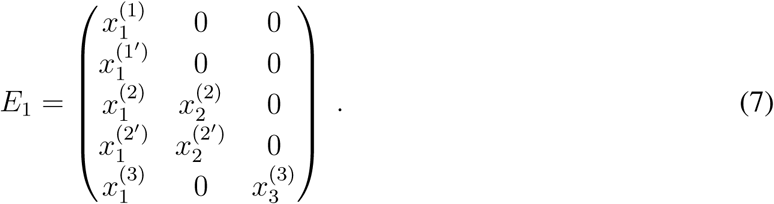

where *x*^(^*^k′^*^)^ is a replicated endpoint containing the same set of species as *x*^(^*^k^*^)^. Of course, the dataset may contain other endpoints in which species *i* is not present: these endpoints will appear in *E*_2_ or *E*_3_, but not *E*_1_. We also highlight the fact that some of these rows (endpoints) will be repeated in other *E_i_*. For example, the third row of *E*_1_ will also be present in *E*_2_, and the last row of *E*_1_ will also appear in *E*_3_. This means that endpoints containing multiple species provide more information than those containing single species, allowing for an efficient experimental design (Extended Data 7, Supplementary Information).

Having formed the matrices *E_i_*, we can recover the *i*^th^ row of *B*, denoted *B_i_*, by solving

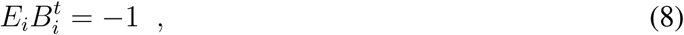

where the right-hand side of Eqn. 8 is a vector of *−*1s with as many elements as there are endpoints in *E_i_*. A general solution to this equation is given by 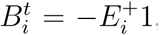, where 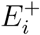 is the Moore-Penrose pseudoinverse of *E_i_*. We write the pseudoinverse rather than the inverse because the matrix *E_i_* need not be square. The use of the pseudoinverse is also convenient because, when analyzing actual experimental data, replicate experiments may yield different abundances for the same endpoint, due to measurement errors or non-point attractors. With the notation introduced above, one can simply list all experimental endpoints, including replicates, in the corresponding matrix *E_i_*, and solve. In this case, there will not be an exact solution, because the system is overdetermined, but the Moore-Penrose pseudoinverse guarantees that the matrix recovered is a maximum-likelihood (least-squares) estimate of *B* given the data.

An example may help clarify this approach. Consider the endpoints recorded in Table 1. These data were generated by adding a small amount of noise to solutions of Eqn. 8, with

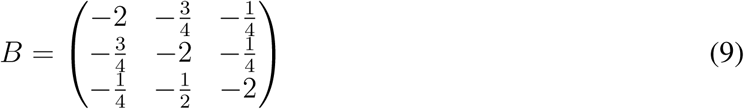

and replicating several endpoints. Constructing *E*_1_ for these data gives:

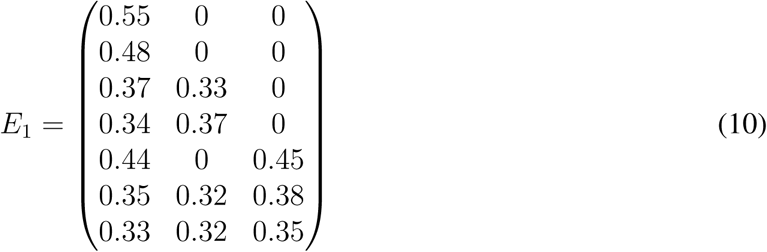

and we recover an estimate of *B*_1_ by computing:

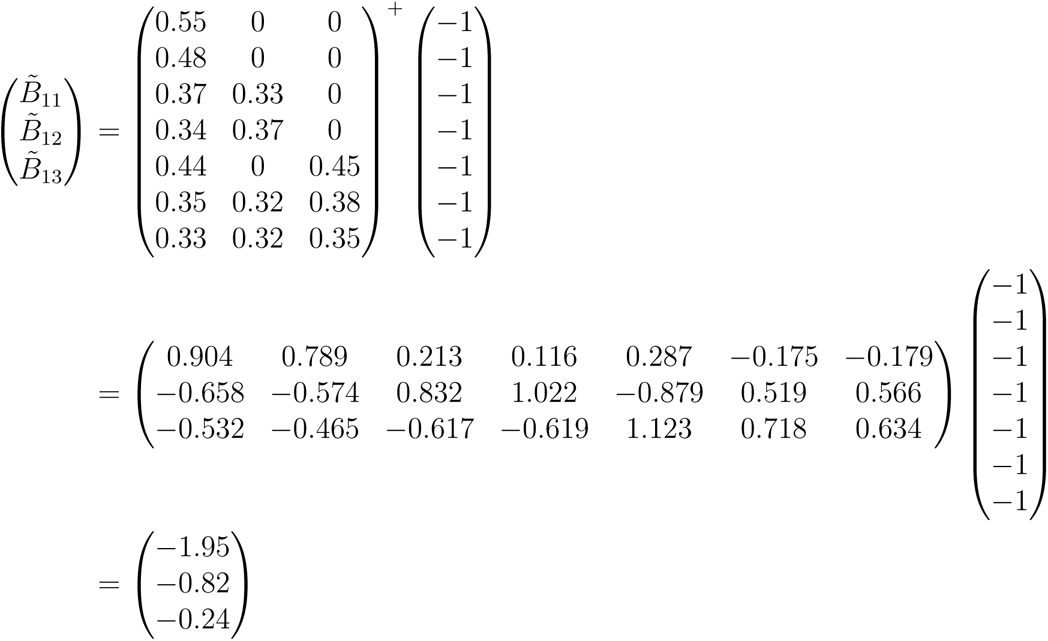

which closely approximates the true coefficients (*−*2*, −*0.74, and *−*0.25). An identical procedure yields estimates for the remaining rows of *B*.

### Requirements

In order to solve for the coefficients in a given row, *B_i_*, it is necessary that the (minimal) structural rank of *E_i_* be *n*, meaning that the rank must remain *n* even if all of the non-zero values of *E_i_* were made identical^33^. This condition is equivalent to simultaneously imposing three biologically-meaningful requirements: (i) each species must be present in at least *n* distinct endpoints, not counting replicates; (ii) each species must co-occur with each other species in at least one endpoint (i.e., for every pair of species *i* and *j*, there must be some endpoint where *i* and *j* co-occur, possibly along with other species); (iii) for each species *i* there must exist a perfect matching between the *n* species and the endpoints in which they co-occur with *i*. Put another way, for a focal species *i*, each endpoint can only “count” once toward the second condition.

These conditions place obvious limitations on the datasets and ecological systems for which our method is applicable. Coexistence among species must be reasonably widespread in order for the first and second conditions to hold. In particular, if a given pair of species *i* and *j* never co-occur, it is impossible to estimate the *B_ij_* relating their endpoint abundances. Systems with significant trophic structure are unlikely to satisfy these conditions. For example, in a linear food chain, the top consumer only occurs in a single endpoint (the full assemblage), violating the first condition. Similarly, systems with strong competitive hierarchies will usually violate these conditions. In general, we envision our method applied to systems where most species are able to persist in isolation, and where the majority of interactions are relatively weak (e.g., many plant and microbial communities).

Finally, we note that our model closely resembles a linear regression on the endpoint abundances. In fact, exactly as in linear regression, if we assume (i) that the endpoints are measured without error and (ii) that the values 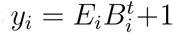 are independently, normally distributed with mean 0 and variance *σ*^2^, then by taking the pseudoinverse we minimize 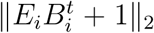 and therefore the variance *σ*^2^. The first of these assumptions is clearly incompatible with the earlier assumption that replicate endpoints are noisy samples from the same attractor. This issue is partially reconciled by imposing the structural rank condition explained above; however, the error structure of our model is still quite distinct from that of a typical linear regression. The fact that these assumptions are not fully appropriate for our setting motivates the development of more sophisticated approaches, which are explained below and employed in practice.

### Accounting for the error structure

The näıve regression approach illustrated above is straightforward, but has several drawbacks. In particular, empirical data will always have some degree of error in the measurements of the densities; moreover, each multi-species endpoint is present in multiple matrices (in the example above, *x*^(3)^ would be reported in *E*_1_ and *E*_3_), and therefore these equations are coupled. Lastly, the maximum likelihood approach above attempts to find the best-fitting *B* which yields an approximate solution, i.e., such that residuals are independent and normally distributed. In doing so, it can allow species’ true endpoints to be quite far from the observed values, such that species may be observed to be present in an endpoint but have a predicted endpoint abundance that is negative.

Thus, although the method outlined here can be näıvely solved using simple linear regression, there is no guarantee that the result is accurate, and it can allow for results that are inconsistent with the biology of the system (e.g., species “coexisting” at negative abundances). We therefore must use a method that allows us to find a matrix *B* such that the corresponding set of true endpoints *z*^(^*^k^*^)^ = *−*(*B*^(^*^k^*^)^)*^−^*^1^1 are as close as possible to the observed endpoints *x*^(^*^k^*^)^ across all replicates and communities *k*.

We present two complementary approaches for estimating *B*, both of which account for the complex error structure and prevent species from coexisting at negative abundances.

First, as detailed in Supplementary Information, one could use a sum-of-squares approach, which uses numerical optimization to minimizes the deviation between the observed and predicted endpoints to get a single estimate of *B*. The benefit of this approach is that it correctly handles the error structure by explicitly incorporating measurement error. It can also be computationally faster than the Bayesian approach, which is detailed next. A drawback to the sum-of-squares approach, however is that it may struggle to find a global maximum, particularly if the likelihood surface for *B* is relatively flat or has many local maxima—a fact that is made more complex by the need to invert *B* (or its submatrices) to calculate the endpoint abundances. This method also does not provide a measure of the uncertainty or standard error surrounding the coefficients, complicating model-selection approaches and preventing one from estimating confidence intervals for the resulting abundance predictions 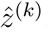. Nevertheless, because of the computational efficiency of this method, we employ it for the analysis of experimental designs for the plant systems and for the simulations, where we are primarily interested in goodness-of-fit of the median value rather than measures of uncertainty.

To address the limitations of the sum-of-squares approach, the most rigorous method for handling the complex error structure—and the one used to fit the three empirical datasets—is a Bayesian MCMC approach, as detailed next. This method allows for probabilistic inference about coexistence, while appropriately handling the complex error structure.

### Measurement error model — a Bayesian approach

The Bayesian MCMC approach is similar to the sum-of-squares approach, but it yields posterior distributions for the elements of the matrix *B* and for each endpoint *z*^(^*^k^*^)^. It allows for standard Bayesian model-selection approaches, and can more easily handle relatively flat likelihood surfaces, provided the priors are chosen appropriately.

To implement this method, we assume there is some underlying *B* which gives rise to the true endpoints, given by *z*^(^*^k^*^)^ = *−*(*B*^(^*^k^*^)^)*^−^*^1^1. The observed endpoints *x*^(^*^k^*^)^ are viewed as random variables, sampled with error from some distribution centered at *z*^(^*^k^*^)^ with vector of standard deviations *σ*^(^*^k^*^)^.

The basics of this approach are as follows:

1. Assign prior distributions for the coefficients of *B* = *{B_ij_}* and *σ* = (*σ, …, σ_n_*)*′*, with the hyperparameters for these distributions encoded in the vector *α*. For the sake of generality we assume a species-specific standard deviation, but one could of course assume a constant standard deviation across all species. Alternatively, one could make more complex assumptions about both *B* and *σ*, for example, that they vary smoothly across environments or that they are phylogenetically correlated across species.
2. Sample 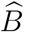 and 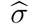 from these distributions, and predict the equilibrium abundance for each observed endpoint *k* by calculating 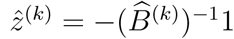.
3. If any species is predicted to have a negative abundance for an endpoint where it was observed to be present, replace it with an arbitrarily small positive value (e.g., 10*^−^*^20^) to ensure this outcome is assigned a near-zero probability of occurrence under a log-normal error structure. Without this step, the model would produce unbiological results, whereby species “coexist” at negative abundances. This step can be modified based on the specific error structure (e.g., if assuming the error are normally distributed rather than log-normal).
4. Calculate the logarithm of the posterior probability for 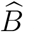 and 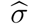 by summing the log-probabilities across all endpoints and replicates, following Bayes’ theorem:

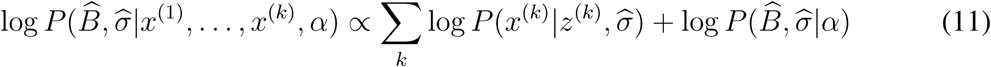 The specific error structure for the endpoints is encoded in the first probability term. Thus, to include a log-normal error structure for *x*^(^*^k^*^)^, as we do for the datasets below, we set 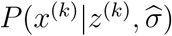 to be the density function of a log-normal distribution with parameters *z*^(^*^k^*^)^ and 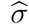.
5. To obtain a posterior distribution for *B*, repeat this process via Markov chain Monte Carlo sampling.

For the datasets analyzed here, we use the *Stan* programming language to implement the MCMC sampling, called via the *stan* function in *R* with the default “No-U-Turn” variant of Hamiltonian Monte Carlo algorithm^34, 35^. For each dataset, we ran four separate MCMC chains for 50,000 iterations each, with a warm-up of 20,000 iterations and thinning every 40 iterations. Posterior plots were investigated to ensure proper mixing of the chains, and “adaptive-delta” parameter was set to 0.85 to minimize divergent transitions. See the data-specific sections (Supplementary Information) for the exact priors and fitting details for each of the three systems.

### Probabilistic inference

The results of this method are posterior distributions for 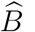, 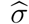, and 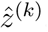which provide estimates of certainty for each value. We can use these posteriors to conduct basic probabilistic inference. For example, for each unobserved or out-of-fit assemblage, we can infer the probability of coexistence by calculating the proportion of posterior *B*s that resulted in all species within the assemblage having positive abundance. Note that this probability cannot be estimated for in-fit data because our method requires that observed assemblages are predicted to coexist (see Step 3, above). For each experimental system, for example, we sampled 1000 bootstrap estimates from the posterior for *B*, and calculated the probability of coexistence for each assemblage (Supplementary Information). With more strict assumptions about the dynamics of the system, one can also infer the probability of local and global stability of the fixed points, for both the observed and unobserved/out-of-fit assemblages (Extended Data 1-6).

## Data and code availability

The data needed to replicate the central findings of this work are available at: https://git.io/fjvON

## ACKNOWLEDGMENTS

We thank S. Kuebbing, C. Rakowski, F. Pennekamp, and O. Petchey for making their data available and for comments on earlier drafts of this manuscript. We thank C. Serván, M. Pascual, J. Bergelson, E. Baskerville, G. Barabás, and E. Friedlander for assistance and suggestions throughout this study. Four anonymous referees provided constructive comments and suggestions.

## Author contributions

DSM, ZRM, and SA conceived of this study and developed the methods. DSM collected and analyzed the data, and wrote the supplemental data analysis. SA and ZRM wrote the supplemental materials and methods and implemented the simulations. All authors contributed to the writing of the manuscript and assisted with revisions.

## Competing interests

The authors declare no competing interests.

---

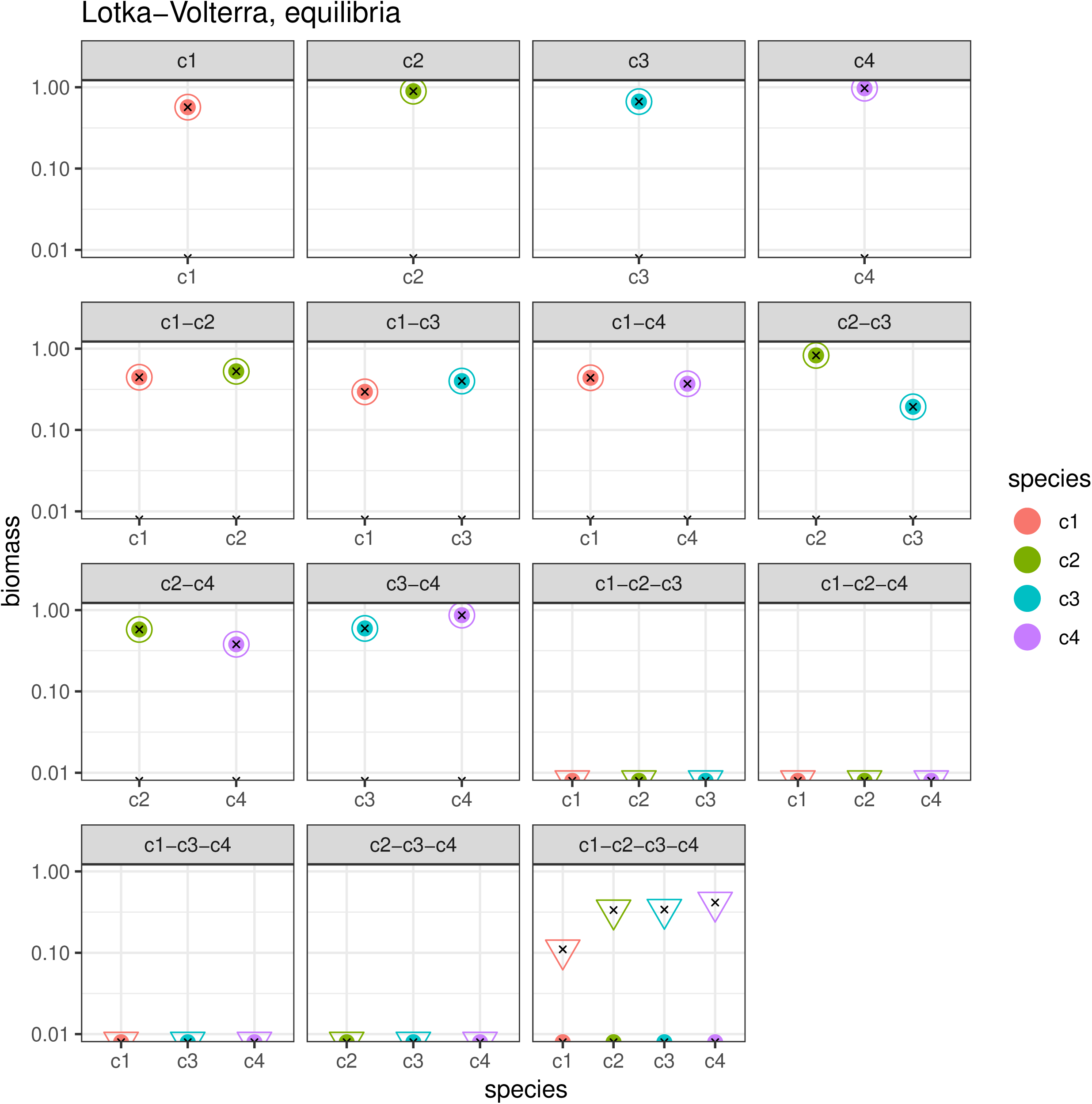

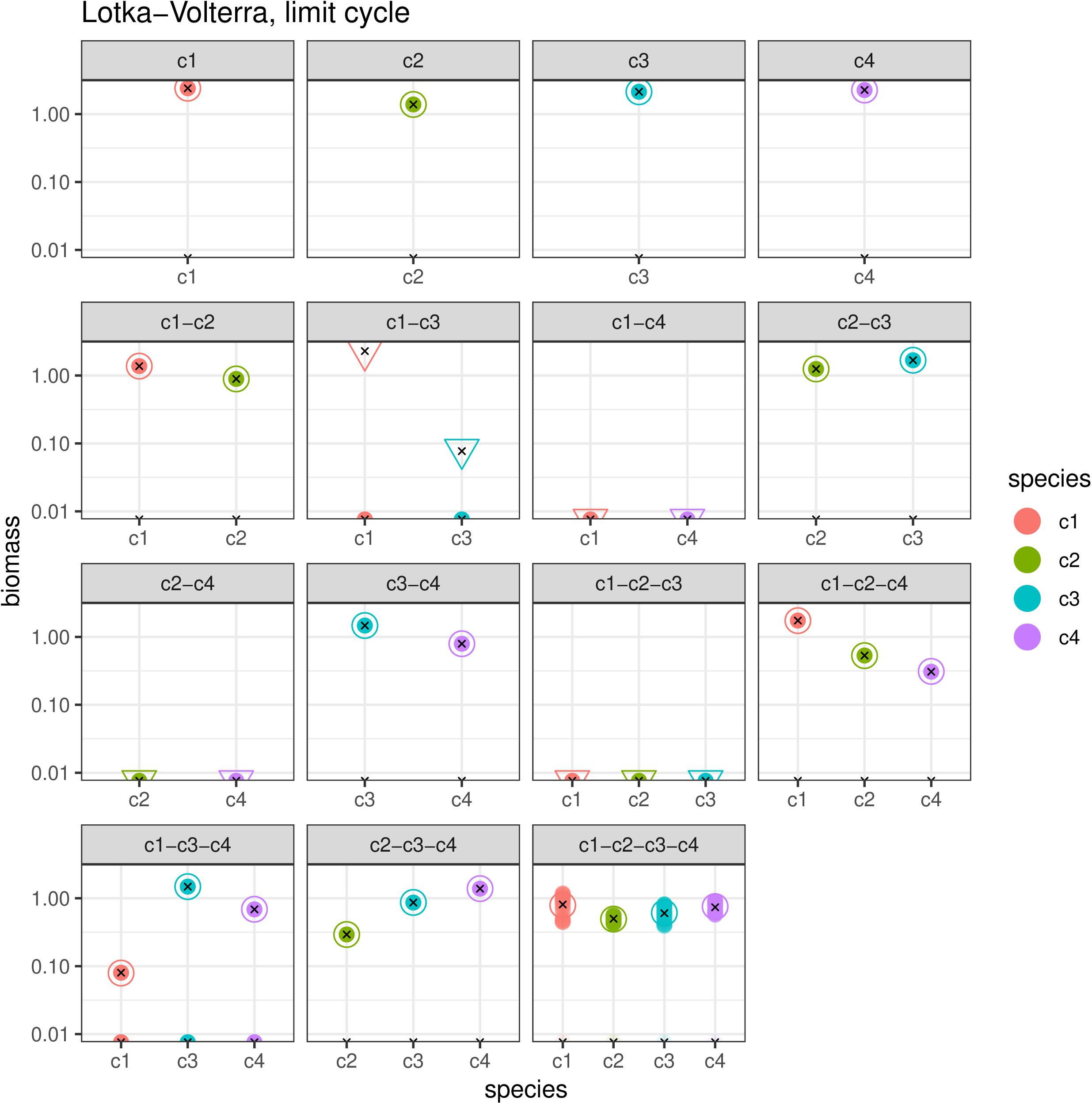

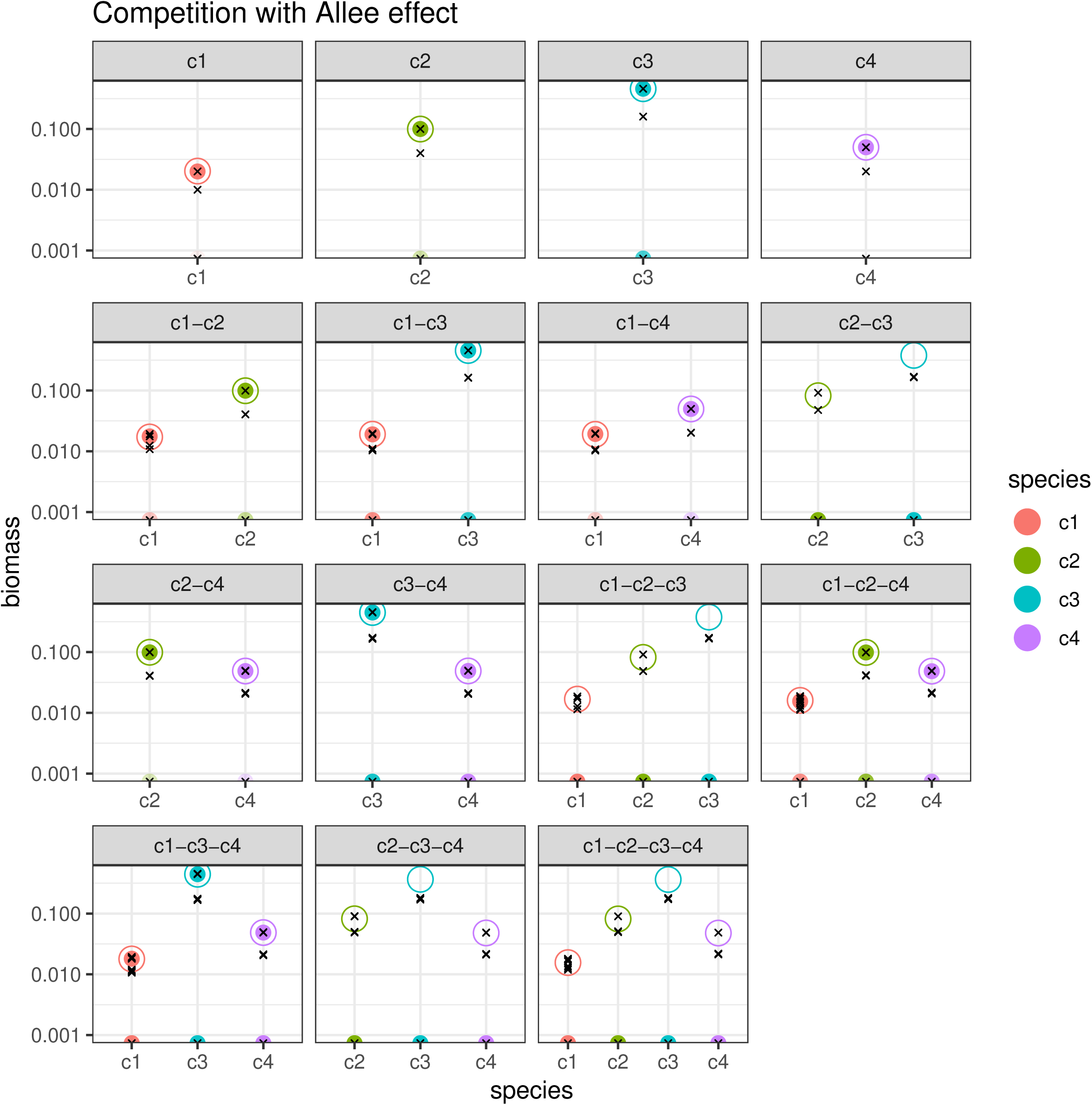

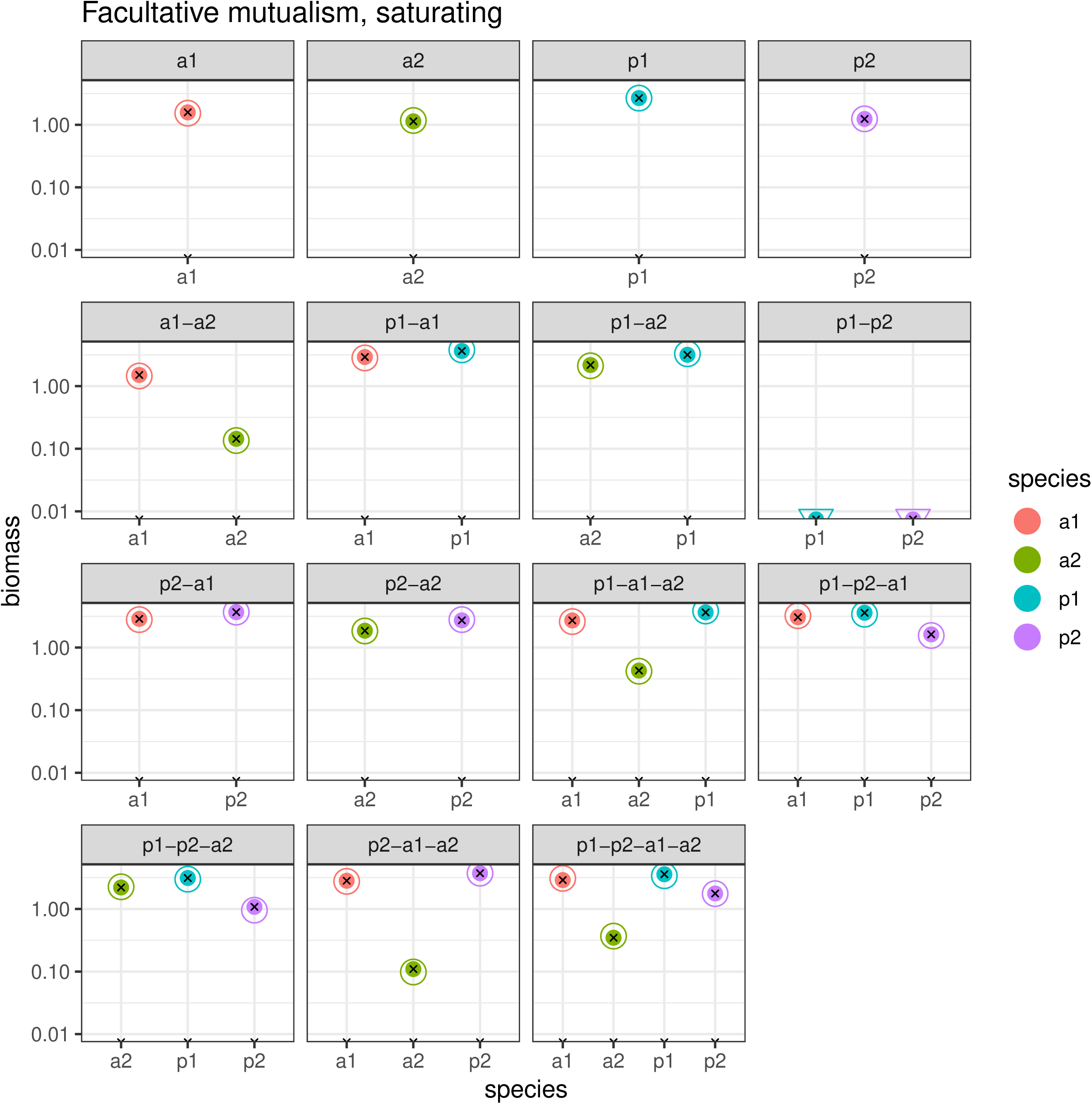

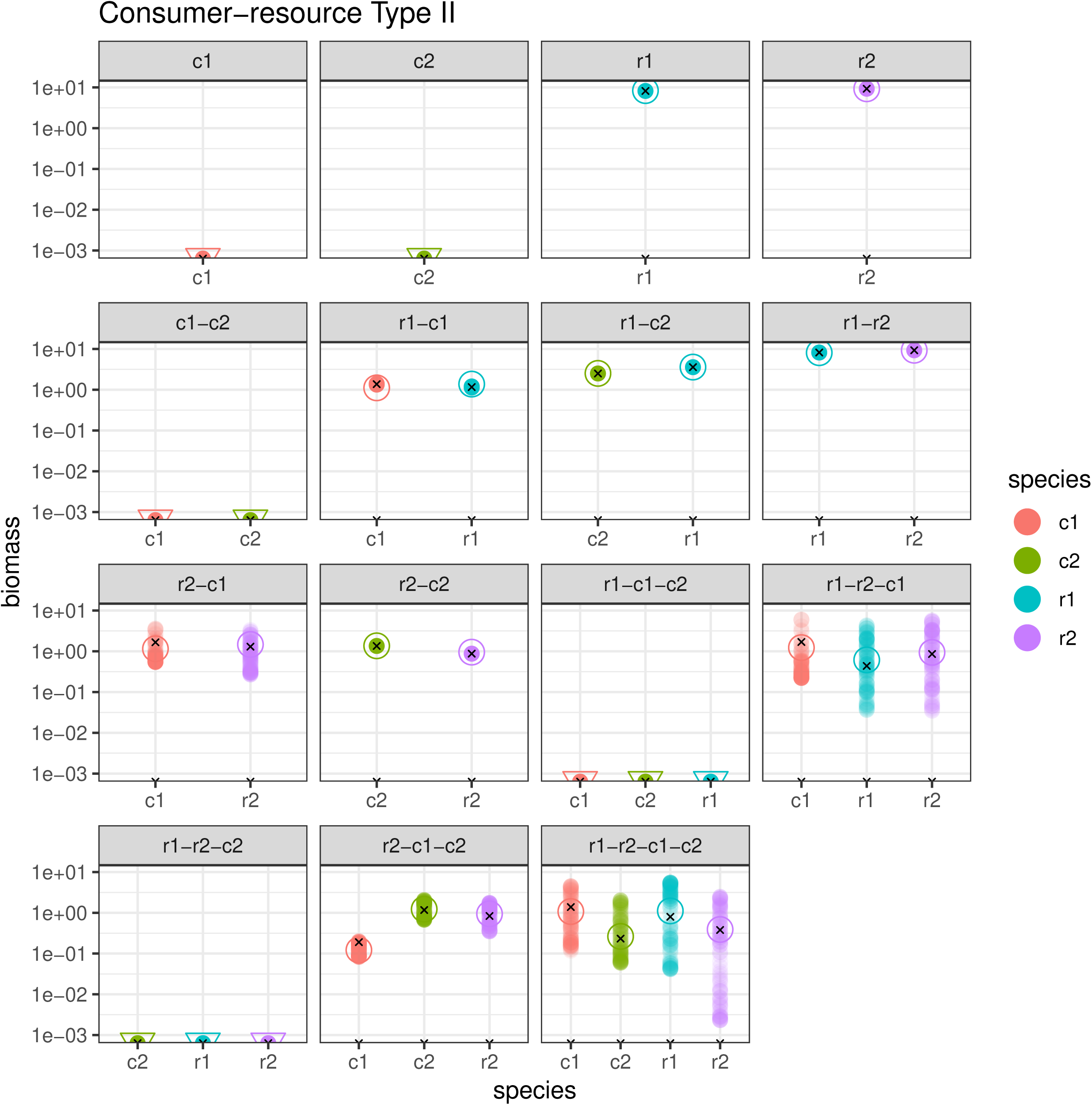

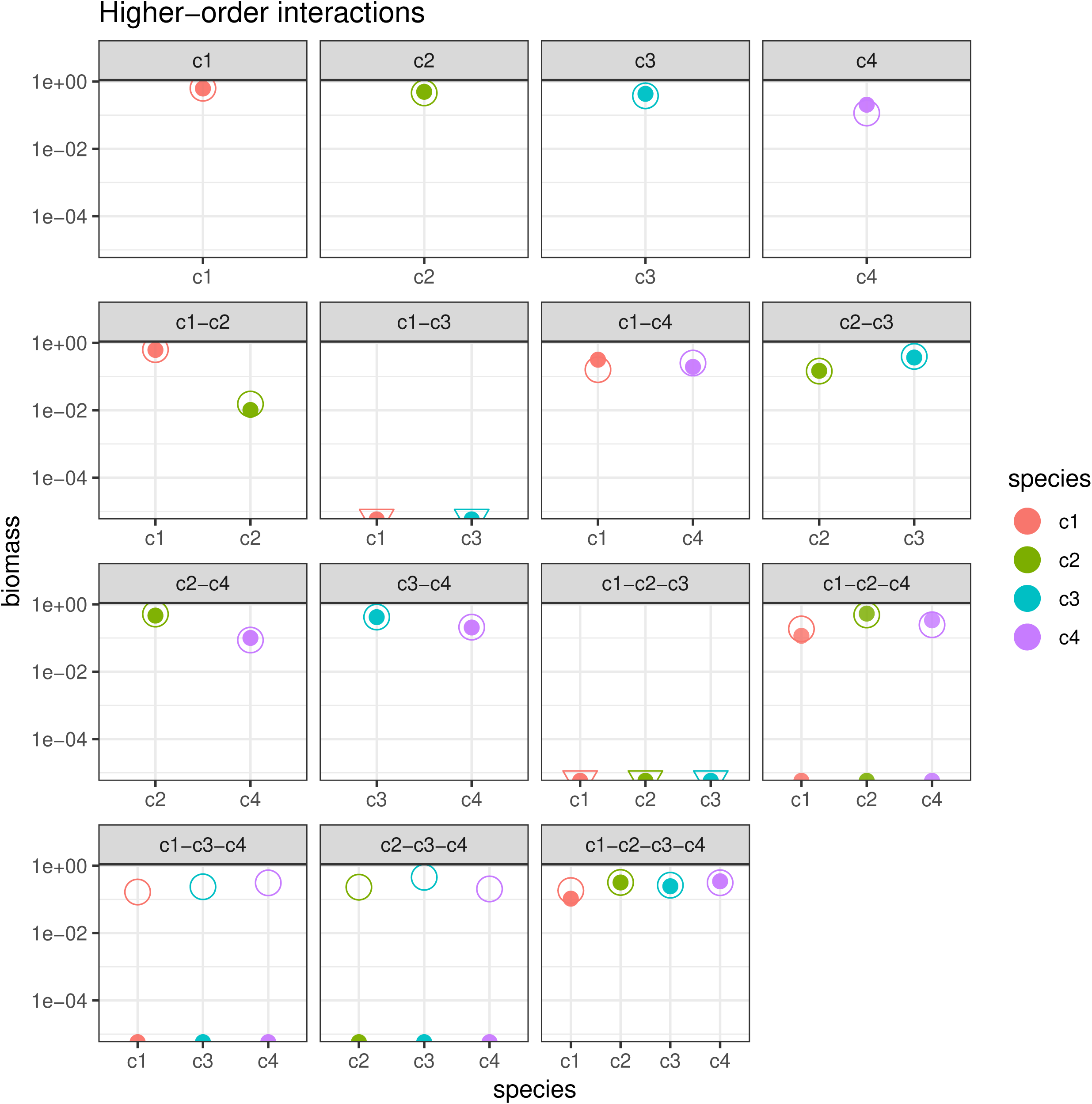

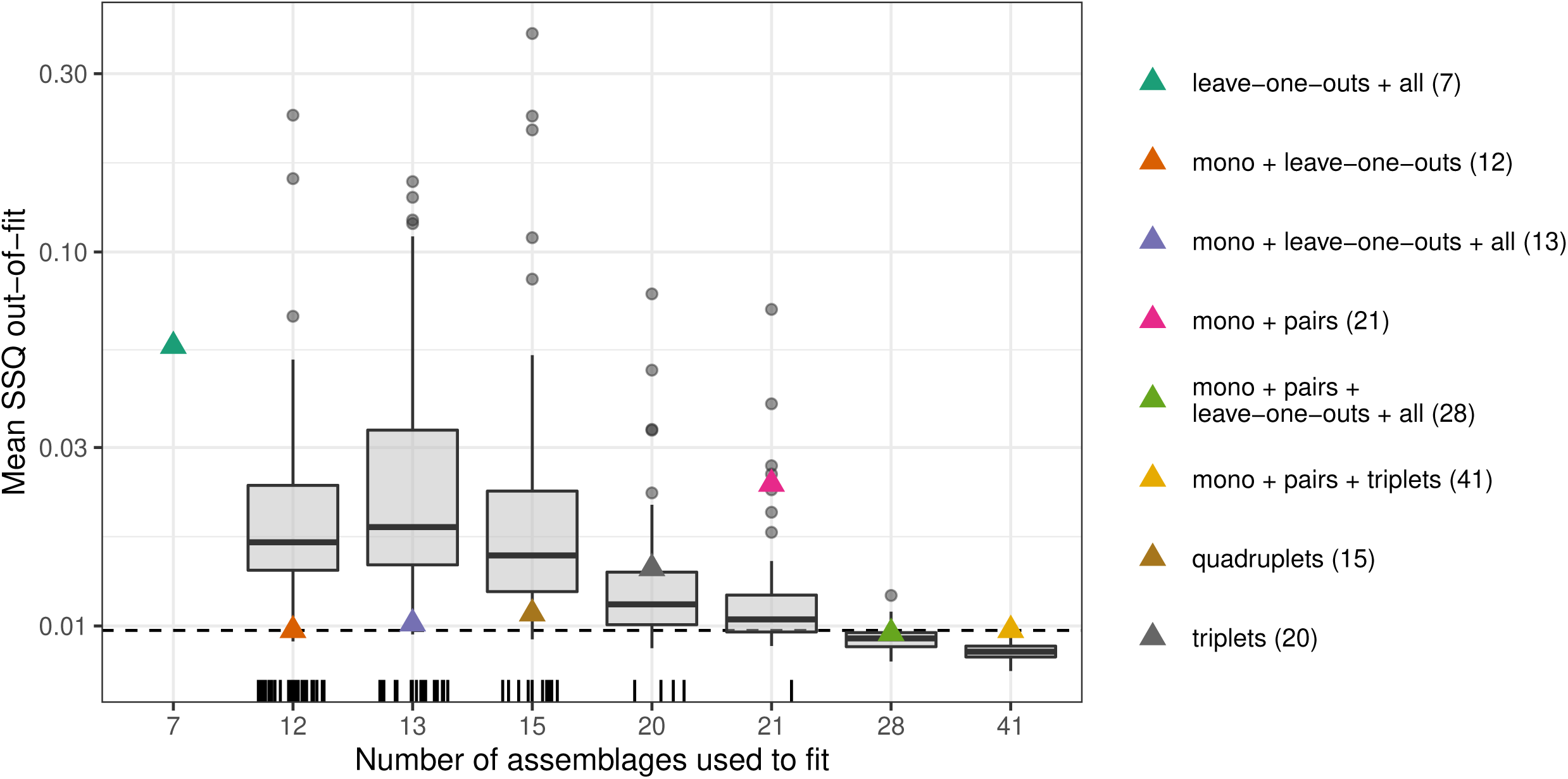

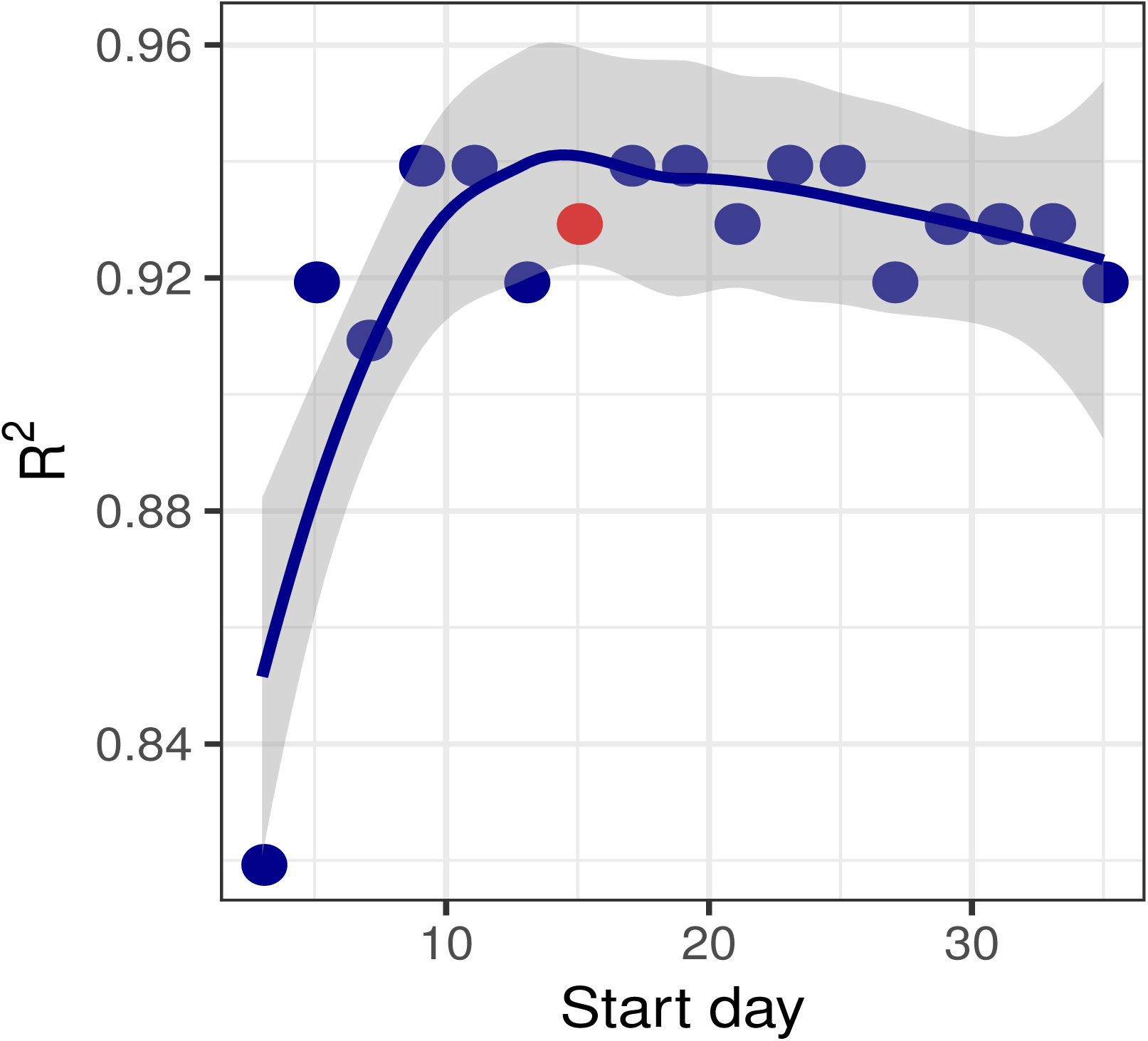

## Notes

https://github.com/dsmaynard/endpoints

