## Supplemental Materials for "Predicting coexistence in experimental ecological communities"

### Predicting coexistence in experimental ecological communities
### Supplementary Information

Daniel S. Maynard,[1,3,*] Zachary R. Miller,[1] Stefano Allesina[1,2]

[1]Department of Ecology & Evolution, University of Chicago
[2]Northwestern Institute on Complex Systems
[3]Institute of Integrative Biology, Dept. of Environmental Systems Science, ETH Zürich, Universitätstrasse 16, 8092 Zürich, Switzerland

### A Supplemental Methods

#### A1 The error structure

Despite the parallels between our inference procedure and standard linear regression, important differences in the error structure necessitate the use of alternative approaches. To illustrate exactly where the regression approach fails, we can consider the standard linear model, in which the endpoint equations would be of the form:

$$y_i^{(k)} = \sum_j B_{ij} z_j^{(k)} + \epsilon_i^{(k)} \tag{1}$$

where $y_i^{(k)} = -1$ and $\epsilon_i^{(k)} \sim \mathcal{N}(0, \sigma^2)$ for all $k, i$.

The main assumption of standard regression is that the predictors, $z_j^{(k)}$, can be measured exactly, such that the error term reflects measurement error in the outcome $y_i^{(k)}$. In the context of endpoint estimation, however, this assumption is inappropriate for two reasons. First, in our model the outcomes $y_i^{(k)}$ are "known" exactly, as they are by definition equal to $-1$. The problem with assuming the $y$ are normally distributed is that it would allow for negative endpoint abundances, which makes no biological sense.

Second, in experimental systems we do not know $z_j^{(k)}$ exactly, but rather we measure it with some error. Thus, the error must be placed on the $z_j^{(k)}$ terms, not on the $-1$ term. If we let $x_j^{(k)}$ denote the noisy, experimentally measured endpoint, we have that $x_j^{(k)} \sim \mathcal{N}(z_j^{(k)}, \sigma_j^2)$, such that the correct error structure for this model is:

$$-1 = \sum_j B_{ij} x_j^{(k)} = \sum_j B_{ij}(z_j^{(k)} + \epsilon_j^{(k)}) = \sum_j B_{ij} z_j^{(k)} + \left[\sum_j B_{ij} \epsilon_j^{(k)}\right] \tag{2}$$

where $\epsilon_j^{(k)} \sim \mathcal{N}(0, \sigma_j^2)$. Now, the error term, $\sum_j B_{ij} \epsilon_j^{(k)}$, is a function of the parameters that we are trying to estimate, contradicting a basic assumption of linear regression.

The problem is even more severe under a log-normal error structure, in which the magnitude of error is proportional to the abundance, as is common when estimating population sizes, and which holds for the datasets analyzed in the main text (e.g., if abundances are measured in a small sample and then extrapolated to the whole system; for example counting cell density using a few *ml* of culture to determine abundance for a culture of 250 *ml*). In this setting, we have $x_j^{(k)} \sim LN(z_j^{(k)}, \sigma_j^2)$, and the error structure for this model becomes:

$$-1 = \sum_j B_{ij} x_j^{(k)} = \sum_j B_{ij} z_j^{(k)} \exp(\epsilon_j^{(k)}) = \sum_j B_{ij} z_j^{(k)} + \left[ \sum_j B_{ij} z_j^{(k)} (\exp(\epsilon_j^{(k)}) - 1) \right] \quad (3)$$

where $\epsilon_j^{(k)} \sim \mathcal{N}(0, \sigma_j^2)$. Here, the so-called error term under a regression framework would be $\sum_j B_{ij} \tilde{z}_j^{(k)} (\exp(\epsilon_j^{(k)}) - 1)$, which is a function of the parameters and the data, preventing any sort of valid, unbiased estimate of $B$ using standard linear regression approaches.

Thus, although the method outlined here can be naïvely implemented using simple linear regression, there is no guarantee that the result will be accurate, and it can allow for results that are inconsistent with the biology of the system (i.e., species "coexisting" at negative abundances). We therefore must use a method that allows us to find a $B$ such that our empirical estimates of the "true" endpoints, given by $\hat{z}^{(k)} = -(B^{(k)})^{-1} 1$, are as close as possible to the set of empirically measured $x^{(k)}$ across all replicates and communities $k$.

We present two complementary approaches for estimating $B$, both of which account for the complex error structure and prevent species from coexisting at negative abundances. In the main text we present a Bayesian MCMC approach that constructs a posterior distribution for the entries of $B$ and the corresponding endpoint abundances. Here, we present an alternative sum-of-squares approach that relies on numerical optimization to find the $B$ that minimizes the deviation between the observed and predicted endpoints.

#### A2 Sum-of-Squares approach.

Because we can estimate all the endpoints $\hat{z}^{(k)}$ for a given $B$, calculated as $\hat{z}^{(k)} = -(B^{(k)})^{-1}1$, the simplest way to allow for complex error structures is to implement a search optimization approach to find the best-fitting $B$ that minimizes the sum of squares (SSQ) of the error between $\hat{z}^{(k)}$ and $x^{(k)}$, taken across all $k$ and all replicates.

By varying how we calculate the SSQ, we can account for various error structures present in the data. For example, if the measurement errors were normally distributed, then we would want to find $B$ that minimized the standard sum of squares (the summation over replicates is omitted for notational simplicity):

$$\min_B ||\hat{z}^{(k)} - x^{(k)}|| = \min_B || - (B^{(k)})^{-1}1 - x^{(k)}|| = \min_B \sum_k \left[-(B^{(k)})^{-1}1 - x^{(k)}\right]^2 \tag{4}$$

Under a log-normal error structure, the maximum likelihood estimate is obtained by taking the sum-of-squares of the difference of the log:

$$\min_B || \log \hat{z}^{(k)} - \log x^{(k)}|| = \min_B \sum_k \left[\log(-(B^{(k)})^{-1}1) - \log(x^{(k)})\right]^2 \tag{5}$$

To implement this approach, we proceed as follows:

1. Initialize $B$ with some starting value, such as a negative diagonal matrix or mean-field initial guess.
2. Calculate the predicted endpoint abundance for all observed subsets of species, given by $\hat{z}^{(k)} = -(B^{(k)})^{-1}1$ for all $k$.
3. If any species is predicted to have a negative abundance for an endpoint where it was observed to be present, replace it with an arbitrarily large value (e.g., $10^{20}$) to penalize this solution to have a large SSQ.

4. Calculate the sum of squares of the deviation between the observed and predicted endpoints, e.g., $\sum_k \left[\log \hat{z}^{(k)} - \log x^{(k)}\right]^2$ under a log-normal error structure.

5. Use an optimization approach to sequentially search for a $B$ that minimizes the sum-of-squares. For example, for the simulations as detailed below, we use the `optim` function in R with the "Nelder-Mead" search algorithm.

The result is that we obtain a best-fitting $B$ encoding endpoints that are most consistent with the observed data under the given error structure. Because all endpoints come from a single instance of $B$, the endpoints are naturally coupled across equations, and because we heavily penalize $B$ matrices where species would incorrectly have negative abundances (i.e., be unable to coexist despite being observed to do so) we ensure that the coexistence patterns given by the resulting $B$ are fully compatible (in terms of presence/absence) with the observed set of endpoints.

The benefit of this approach is that it correctly handles the error structure by explicitly incorporating measurement error. It can also be computationally faster than the Bayesian approach, which is detailed in the main Methods. A drawback with this approach is that it can struggle to find a global maximum, particularly if the likelihood surface for $B$ is relatively flat or has very many local maxima—an issue that is made more acute by the need to invert $B$ (or its sub-matrices) to calculate the endpoint abundances. This method also provides a single best-fitting $B$, without any measure of the uncertainty or standard error surrounding the coefficients. This complicates model-selection approaches, and prevents one from estimating confidence intervals for the resulting abundance predictions $z^{(k)}$. To address these limitation, in the main text we have outlined a Bayesian MCMC approach, which was used for the analysis of the empirical datasets presented in the main text.

#### A3 Inference under Lotka-Volterra dynamics

One might recognize that the set of endpoints encoded by Eqn. 1 (Main text) is identical in structure to the set of equilibria of a generalized Lotka-Volterra (GLV) model. This model has been studied for almost a century[1,2], and many of its properties are well-known (for an excellent survey, see[3]). In the GLV model, the dynamics of species' abundances are modeled by the set of differential equations:

$$\frac{\mathrm{d}x_i(t)}{\mathrm{d}t} = x_i(t)\left(r_i + \sum_j A_{ij}x_j(t)\right) = r_i x_i(t)\left(1 + \sum_j \tilde{A}_{ij}x_j(t)\right) \quad , \tag{6}$$

Here, as above, $x_i$ is the abundance (often density or biomass) of species $i$, now indexed by time, $t$. The $r_i$ are intrinsic growth rates for each species, describing the rate of growth of species $i$ when grown in isolation at low abundance. The $A_{ij}$ are interaction coefficients that specify the effect of species $j$ on the per-capita growth rate of species $i$. The coefficients $\tilde{A}_{ij} = A_{ij}/r_i$ are composite parameters, interpretable as interaction strengths normalized by growth rate. This reformulation (in terms of $\tilde{A}$) has been exploited before to infer interaction strengths from experimental data (e.g.,[4,5]).

Fixed points (equilibria) of the GLV system can be found by setting the left-hand side of Eqn. 6 to $0$. For all species not present at the fixed point, the resulting equations are satisfied trivially. For species present at non-zero abundance, and assuming non-zero growth rates, one can divide both sides of the corresponding equation by $r_i x_i$, yielding

$$0 = 1 + \sum_j \tilde{A}_{ij}x_j^\star \ , \quad \forall\, i \ \text{ such that } \ x_i^\star \neq 0 \ \ . \tag{7}$$

The form of Eqn. 7 is precisely the same as Eqn. 1 (Main), with $\tilde{A}$ playing the role of $B$. This correspondence indicates that whenever the dynamics of a system are well-described by a GLV model, our assumptions regarding the relationships between endpoints will hold, and our method should perform well. However, we emphasize that our method is agnostic to the dynamics underpinning the system, and does not assume or require either GLV dynamics or equilibrium conditions. And while these conditions should lead to high performance by our method, high performance does not necessarily imply that the underlying dynamics are GLV. In our modeling framework, the coefficients of $B$ are interpretable only in relation to endpoint abundances; away from the system's attractors (e.g., in the transient phase), the interactions between species may be very different. In a GLV model, on the other hand, one must assume that these interaction coefficients describe the dynamics at all times.

With these cautions in mind, the correspondence between our model and the GLV equilibrium structure motivates us to ask whether additional inference is possible under the more stringent assumption of GLV dynamics. That is, if we now assume the dynamics are specified by Eqn. 6, with $\tilde{A} = B$, what else can we learn from endpoint data? We find that the matrix $B$ is informative about the sign pattern of the matrix $A$ (the unnormalized interaction strengths), the relative magnitudes of the $A_{ij}$, and the invasibility of the system's endpoints, even with no knowledge of the growth rates $r_i$. $B$ also provides more limited information regarding the (local) stability of endpoints. We briefly explain the inference procedure for each of these properties below.

The growth rate of a species in isolation, $r_i$, may be positive (e.g., for producers) or negative (e.g., for consumers). As such, it is not possible to directly infer the sign of the interaction coefficient $A_{ij}$ from the estimate of $B_{ij}$. However, one may reasonably assume that all species display some form of self-regulation at sufficiently high density, which implies that the diagonal elements $A_{ii}$ are negative. Because the matrix $B$ is related to $A$ by a rescaling of each row by the corresponding growth rate, a nonnegative value for $B_{ii}$ is likely to indicate that $r_i < 0$. Using this information, it is straightforward to "flip" all of the signs in the appropriate rows to recover the sign pattern of $A$. In the GLV setting, where interactions are consistent in time, the signs of the pair $(A_{ij}, A_{ji})$ specify the type of interaction between species $i$ and $j$ (e.g., competition, mutualism, predation, etc.).

By the same observation that $B$ is related to $A$ by a rescaling of the rows, it is clear that the elements of $B$ can be meaningfully compared within rows. For two values, $B_{ij}$ and $B_{ik}$, the growth rate $r_i$ cancels and $\frac{B_{ij}}{B_{ik}} = \frac{A_{ij}}{A_{ik}}$. The value of $A_{ij}$ indicates the strength of the per-capita effect of species $j$ on species $i$. Comparing values within row $i$, one can see the relative effect of all species on $i$; for example, if $B_{ij}$ is twice as large as $B_{ik}$, then the effect of $j$ on the growth of $i$ is twice as strong than that of $k$. We cannot, however, compare elements across different rows (as they have been divided by different quantities). This means it is impossible to infer which other species a focal species $i$ affects more or less strongly.

Having inferred the signs of the $r_i$, one can also assess the invasibility of any equilibrium community (observed or unobserved). In community ecology, the invasibility of equilibria is often used to draw conclusions on coexistence. Typically, one considers a community $S$ resting at the equilibrium $x^\star$ with species $i$ absent, and asks whether $i$ can grow when introduced at low abundance. This translates into the criterion:

$$r_i + \sum_{j \in S} A_{ij} x_j^\star > 0 \tag{8}$$

or

$$\operatorname{sign}(r_i) \left(1 + \sum_{j \in S} B_{ij} x_j^\star \right) > 0 \tag{9}$$

This means that invasibility analysis requires knowing only the sign of $r_i$, and not its magnitude (if $r_i < 0$, the inequality is simply reversed). Therefore, using the procedure described above to infer the signs of $r$, invasibility can be inferred solely from the matrix $B$.

Finally, one is very often concerned with the stability of equilibria. As noted above, even if an endpoint (in the GLV setting, an equilibrium point) is feasible, it might be unstable, in which case it may be impossible to reach experimentally (note, however, that unstable equilibria may still be associated with observable attractors, including stable limit cycles or chaotic attractors). Therefore, it is desirable to be able to predict the stability of endpoints in order to accurately predict coexistence in practice.

For GLV systems, the stability of an equilibrium $x^\star$ is determined by the community matrix $M = D(x^\star)A = D(r)D(-B^{-1}1)B$ (where $D(y)$ denotes the diagonal matrix with the vector $y$ on the diagonal). If all of the eigenvalues of $M$ have negative real part, then the feasible equilibrium $x^\star = -B^{-1}1$ is *locally asymptotically stable*, meaning that when the system is initialized sufficiently close to the equilibrium, it will eventually reach it. A stronger notion of stability is *global stability*, meaning that the equilibrium will be reached whenever all the species are initialized at positive densities. A sufficient condition for the global stability of an equilibrium of the GLV model is Lyapunov diagonal stability of $M$. In turn, a sufficient condition for Lyapunov diagonal stability is $M + M^t$ having all negative eigenvalues.

In general, then, it is necessary to know the growth rates $r_i$ to compute $M$ and determine the stability of an equilibrium. However, in certain cases the matrix $B$ is sufficient. To see this, note that if a matrix $Y$ is Lyapunov diagonally stable, then $D(z)Y$ is also Lyapunov diagonally stable, for any positive vector $z$. We can exploit this fact in the following way: Flip the signs of $B$ to obtain a matrix $C$ where $A = D(\text{abs}\,(r))C$ (where abs denotes the absolute value operator, applied element-wise). Now, if $C + C^t$ has all negative eigenvalues (or, in fact, if one can find any strictly positive $z$ such that $D(z)C + C^t D(z)$ has all negative eigenvalues), then $C$ is Lyapunov diagonally stable, and consequently so are $A$ and, by the feasibility of $x^\star$, $M$. In short, if one can demonstrate the Lyupanov diagonal stability of $C$, which is obtained from $B$, then the corresponding feasible equilibrium is globally stable regardless of the magnitudes of the $r_i$.

Even in cases where $C$ cannot be shown to be Lyupanov diagonally stable, it may still be possible to learn about the stability of an equilibrium. One approach is to simply check whether $D(-B^{-1}1)C$ has all negative eigenvalues. If so, this implies that the equilibrium will be stable for growth rates of equal magnitude (i.e. $|r_i| = |r_j|$ for all $i$ and $j$). Because the eigenvalues are a continuous function of the coefficients, this inference is somewhat robust to small deviations from strict equality. In other words, if the growth rates differ but have low variance, the equilibrium is likely to be stable. As another approach, if one assumes some distribution for the $r_i$, then the probability of stability can be estimated numerically by drawing many random growth rate vectors, using these to form $M$ matrices, and checking their stability.

### B Data analysis

#### B1 Methods Overview

For each of the three systems detailed below, encompassing 10 unique datasets, we implemented a Bayesian approach to obtain a posterior distribution for $\hat{B}$ and each endpoint $\hat{z}^{(k)}$. In every case, we first conducted the analysis using the full set of endpoints, providing a best-fit estimate of $B$. Second, we tested the ability of each model to predict unobserved assemblages using a jackknife approach: we sequentially removed each endpoint $k = 1, \ldots, m$, one at a time, and used the remaining $m - 1$ communities to estimate $B$ in the absence of endpoint $k$. This 'leave-one-out' dataset was then used to predict the omitted endpoint(s), as outlined above, providing a probability of feasibility of such assemblages, as well as a posterior distribution for each species' abundance in the omitted assemblage.

***Note:*** Species abbreviations refer to the first two letters of genus name (protist), the family name (plants), or the initials of the Latin binomial (herbivore-algae).

#### B2 Plant system — data from Kuebbing *et al.*, 2015

**Description of the experiment.** Kuebbing *et al.*[6] explored how species' richness and identity affect plant productivity and seedling establishment. The experimental system comprised four native plant species and four non-native plant species that are common in eastern Tennessee old-fields. The plants were phylogenetically paired across the native and non-native subsets, resulting in one species from the families Asteraceae, Fabaceae, Lamiaceae, Poaceae in each pool (Table S1).

Table S1: **The eight old-field plant species, split into phylogenetically paired native and non-native plant pools. Adapted from Table 1 of Kuebbing *et al.* (2015)**.

| Family | Abbrv. | Native species | Non-native species |
|---|---|---|---|
| Asteraceae | as | *Achillea millefolium* L. | *Leucanthemum vulgare* Lam. |
| Fabaceae | fa | *Lespedeza capitata* Michx. Hornem. | *Lespedeza cuneata* (Dum. Cours.) G. Don |
| Lamiaceae | la | *Pycnanthemum virginianum* Schrad. | *Prunella vulgaris* L. var. *vulgaris* |
| Poaceae | po | *Sorghastrum nutans* (L.) Nash | *Phleum pratense* L. |

Native species were grown only with other natives, and non-natives were grown only with other non-natives, resulting in two distinct pools of four species, each giving rise to 15 (i.e., $2^4 - 1$) possible assemblages. For each pool of plants, species were grown in monoculture and in all but 1 of the 15 multi-species combinations, which was omitted due to a limited number of viable seedlings.

Assemblages were grown in square pots ($13 \times 13 \times 17$ cm) consisting of 1:1 volumetric ratio of autoclaved sand and field soil, with each assemblage replicated 20 times (10 replicates used here for biomass estimates). Each replicate consisted of 12 individual seedlings in a $3 \times 4$ grid, with locations randomly assigned. Seedlings that died within one week of initial planting were replaced to remove the effect of transplant stress. Pots were fertilized at days 50 and 100 (20:20:20 nitrogen:phosphorus:potassium), and watered biweekly or more often if needed. Pots were incubated in a glasshouse at the University of Tennessee, Knoxville, TN, USA.

Ten replicates of each assemblage were harvested after 112 days of growth. Aboveground biomass was clipped, sorted, and dried at $60°$C for 48 hours. The resulting per-species biomass for each pot is used here to estimate endoints, given in grams ($g$) dry biomass per pot. See Kuebbing *et al.*[6] for a more detailed description of the study and experimental design.

**Stan fitting details.** To properly scale the data, we first estimated the maximum abundance for each species, calculated as the upper $95^{th}$ percentile across all communities. We then divided each species' abundances in all assemblages by their respective maximum value, such that their self-regulation term $B_{ii}$ would be approximately -1 (assuming competitive or neutral interactions). Under the null hypothesis that species are neutral with respect to each other, the MCMC chains were initialized with $B$ equal to the negative identity matrix, i.e.:

$$\mathbf{B} = \begin{array}{c} \\ as \\ fa \\ la \\ po \end{array} \begin{array}{c} \begin{array}{cccc} as & fa & la & po \end{array} \\ \begin{pmatrix} -1.00 & 0 & 0 & 0 \\ 0 & -1.00 & 0 & 0 \\ 0 & 0 & -1.00 & 0 \\ 0 & 0 & 0 & -1.00 \end{pmatrix} \end{array}$$

For both plant pools, we used non-informative priors for the coefficients $B_{ij}$, with each entry sampled from a Normal $\mathcal{N}(0, 10)$ distribution. We additionally constrained diagonal entries to be negative, to reflect the fact that this is a competitive system where each species exhibits positive growth in monoculture. The species-specific standard errors were sampled from a normal $\mathcal{N}(0, 0.25)$ distribution.

**Results.** The median posterior $B$ values for the native plant system were:

$$\mathbf{B} = \begin{array}{c} \\ as \\ fa \\ la \\ po \end{array} \begin{array}{c} \begin{array}{cccc} as & fa & la & po \end{array} \\ \begin{pmatrix} -0.45 & -0.05 & -0.13 & -0.20 \\ -0.28 & -0.16 & -0.43 & -0.18 \\ -0.28 & -0.06 & -0.96 & -0.18 \\ -0.28 & -0.04 & -0.06 & -0.29 \end{pmatrix} \end{array}$$

And for the nonnative plant system:

$$\mathbf{B} = \begin{array}{c} \\ as \\ fa \\ la \\ po \end{array} \begin{array}{c} \begin{array}{cccc} as & fa & la & po \end{array} \\ \begin{pmatrix} -0.48 & -0.07 & -0.15 & -0.22 \\ -0.30 & -0.10 & -0.22 & -0.25 \\ -0.34 & -0.07 & -0.36 & -0.23 \\ -0.32 & -0.07 & -0.21 & -0.48 \end{pmatrix} \end{array}$$

The full sets of predictions for these two systems are depicted in Figs 3-4. As noted in the main text, this approach is able to recover with high accuracy the median endpoint abundances ($R^2 = 0.98$ for observed vs. predicted), and exhibits strong agreement between the predicted and observed variation in abundances for each species. Both systems show high likelihood of feasibility for all subsets—consistent with no species going extinct in any replicate—with only the full four-species nonnative system having an out-of-fit predicted probability of coexistence of $< 95\%$.

These results show a few interesting biological features, such as the Fabaceae species in both systems having the smallest negative effects (column $Fa$), consistent with the species being nitrogen fixers. In order to directly compare the structure of these matrices, we need to scale each row by the inverse of the magnitude of the diagonal, $|1/B_{ii}|$, so as to remove the effect of self-regulation (carrying-capacity) on the magnitude of the interactions. This scales the interspecific effects relative to the magnitude of the intraspecific effect.

For the native, this yields:

$$\mathbf{B_{scaled}} = \begin{array}{c} \\ as \\ fa \\ la \\ po \end{array} \begin{array}{c} \begin{array}{cccc} as & fa & la & po \end{array} \\ \begin{pmatrix} -1.00 & -0.11 & -0.29 & -0.45 \\ -1.76 & -1.00 & -2.74 & -1.13 \\ -0.29 & -0.06 & -1.00 & -0.19 \\ -0.97 & -0.15 & -0.22 & -1.00 \end{pmatrix} \end{array}$$

And for the nonnative plant system:

$$\mathbf{B_{scaled}} = \begin{array}{c} \\ as \\ fa \\ la \\ po \end{array} \begin{array}{c} \begin{array}{cccc} as & fa & la & po \end{array} \\ \begin{pmatrix} -1.00 & -0.14 & -0.31 & -0.46 \\ -2.98 & -1.00 & -2.20 & -2.42 \\ -0.95 & -0.21 & -1.00 & -0.65 \\ -0.67 & -0.14 & -0.43 & -1.00 \end{pmatrix} \end{array}$$

Because the error structure for the coefficient is log-normal, we must take the $\log$ of the coefficients to compare them on the correct scale. Since the coefficients are all negative, we thus take the $\log(-B_{ij})$, ensuring that we are taking the $\log$ of positive numbers. Comparing these two systems, we see a close relationship between the structure of the scaled $B$ matrices (Fig. 3a, main text).

This result suggests that the effect one species has on another is phylogenetically conserved across the two data sets, such that, for example, the relative effect that a Lamiaceae has on the abundance of a Poaceae is approximately the same across the two systems.

As discussed above, if we are willing to assume Lotka-Volterra dynamics for these systems, we can then make additional assumptions about the stability and invasibility of endpoints. For example, in these systems, all communities exhibit high probability of local and global stability

(Figs 3-4c), albeit with the non-native community showing slightly lower probability of global stability compared to the natives. These results also suggest that nearly every species can invade into every community (Figs 3-4d), likely due to strong self-limiting (diagonal) terms in $B$. Though these results are speculative, due to the fact that these dynamics likely deviate from GLV dynamics to some extent, such information could help design and select future assemblages for experimentation.

#### B3 Herbivore-algae system — data from Rakowski and Cardinale, 2016

**Description of the experiment.** Rakowski and Cardinale[7] tested how herbivory affects the relationships between species richness and biomass. The experimental system consisted of five species of freshwater algae: *Chlorella sorokiniana, Scenedesmus acuminatus, Pediastrum duplex, Monoraphidium minutum* and *Monoraphidium arcuatum*; and two species of freshwater herbivores from the family Daphniidae: *Ceriodaphnia dubia* and *Daphnia pulex*. Here we focus only on those communities in which the algae *P. duplex* was absent (Table S2), as this species did not survive in enough microcosms to estimate its interaction parameters.

Table S2: **The two Daphniidae species and four algae species use for analysis here, from Rakowski and Cardinale (2016).**

| Abbrv. | Species | Functional group |
|---|---|---|
| dp | Daphnia pulex | Daphniidae (herbivore) |
| cd | Ceriodaphnia dubia | Daphniidae (herbivore) |
| cs | Chlorella sorokiniana | Chlorophyta (algae) |
| ma | Monoraphidium arcuatum | Chlorophyta (algae) |
| mm | Monoraphidium minutum | Chlorophyta (algae) |
| sc | Scenedesmus acuminatus | Chlorophyta (algae) |

The algal species were grown in monoculture (5 unique communities) and in all four-species combinations ($\binom{5}{4} = 5$ unique multi-species combinations). These 10 unique assemblages were each replicated 15 times, resulting in 150 total microcosms. All replicates were assembled in

1-liter bottles filled with 750-ml nutrient-rich COMBO medium, and inoculated either with 400,000 cells of one species for the monoculture treatment, or with 100,000 cells for each of the four species. Microcosms were incubated at 20°C on a 16:8 hour light:dark cycle. After 12 days, 23 adult individuals of *C. dubia* were added to 5 of the 15 replicates for each assemblage, and 15 adult individuals of *D. pulex* were added to a different 5 replicates; the remaining 5 replicates of each algal assemblage had no herbivores added. The different initial densities of the two herbivores were selected to ensure approximately equal initial biomass.

Starting upon herbivore addition (day 0) and every 2 days afterward, 8% of the media was exchanged with fresh media. The abundance of each algal species was estimated every six days, starting at day 0 and ending at day 24 (5 measured time points) by counting the number of cells of each species per sample, stopping when either 1.8 $\mu$l of sample was analyzed or 400 cells were counted. Likewise, herbivore abundances were estimated every 6 days, starting at day 10 and ending at day 28 (4 sampling time points) by counting the number of herbivores visible in the bottle, estimated to the nearest multiple of 5.

Because densities were estimated using a small subsample of the microcosm, species that were recorded as "extinct" were typically present in very low abundances in the full community (C. Rakowski, personal communication), such that recorded absences reflect "effective" extinctions. For our purposes, mistaking effective extinctions as true extinctions (false negatives) is far less problematic than incorrectly assuming a species is present when in fact it is on the path to extinction (false positive). False negatives add noise to the true endpoint solution, whereas false positives force the fitted hyperplanes through a non-existent endpoint, which may strongly alter the overall structure of endpoints. We thus used the reported raw counts of the number of herbivores and algae in each microcosm at the end of the experiment (day 28 for the herbivores, day 24 for the algae) as the endpoint abundances, assuming zero abundances represented true absences. To further minimize the presence of false positives, we removed an additional 6 assemblages (9 total microcosms) which had fewer than three replicates each.

The remaining 116 microcosms encompassed 26 of the $62 = 2 \times (2^5 - 1)$ possible endpoint patterns, 9 of which contained algae only (47 total microcosms), 10 of which contained *D. pulex* + algae (42 total microcosms), and 7 of which contained *C. dubia* + algae (27 total microcosms). These set of endpoints were sufficient to estimate $B$ for the *D. pulex* and *C. dubia* sub-communities, separately.

**Stan fitting details.** To properly scale the data, we first estimated the maximum abundance for each species, calculated as the upper $95^{th}$ percentile across all communities. We then divided each species' abundances in all assemblages by their respective maximum value, such that their self-regulation term $B_{ii}$ would be approximately -1.

For the algae, we constrained the diagonals of $B$ to be negative, consistent with these species having positive growth in monoculture. For each herbivore $i$, we constrained $\gamma_i \leq 0$, consistent with this species going extinct in the absence any resources; and we constrained $\tau_{ij} > 0$ for all algae $j$, consistent with the herbivore feeding on the algae. When rescaling the herbivore coefficients for $B$ (indexed by $i$), we therefore have that $B_{ii} = -1/\gamma_i > 0$, and that $B_{ij} = \tau_{ij}/\gamma_i < 0$. That is, even though the herbivore feeds on the algae, the non-diagonal row coefficients $B_{ij}$ are negative because the $\tau_{ij}$ (which are positive) are then scaled by the herbivore's monoculture carrying capacity $1/\gamma_i$ (which is negative). This fact highlights why mechanistic inference of $B$ can be difficult, as $B$ is not identical to a traditional interaction matrix—which, under standard Lotka-Volterra dynamics, for example, would have positive row coefficients for the effect of algae on the herbivore.

Finally, we assumed a mean-field consumptive effect to initialize $B_{ij}$, such that the algae are interchangeable resources for the herbivore and thus have the same magnitude of coefficients. Additional, we guess initially that algae interact neutrally with one another. Together, these assumptions entail that differences in abundances between communities of the same size are purely an artifact of measurement error.

Under these assumptions, $B$ was initialized as:

$$\mathbf{B} = \begin{array}{c} \\ dp \\ cs \\ ma \\ mm \\ sa \end{array} \begin{array}{c} \begin{array}{ccccc} dp & cs & ma & mm & sa \end{array} \\ \begin{pmatrix} 1.00 & -1.00 & -1.00 & -1.00 & -1.00 \\ -1.00 & -1.00 & 0 & 0 & 0 \\ -1.00 & 0 & -1.00 & 0 & 0 \\ -1.00 & 0 & 0 & -1.00 & 0 \\ -1.00 & 0 & 0 & 0 & -1.00 \end{pmatrix} \end{array}$$

We used non-informative priors for the coefficients $B_{ij}$, with each entry sampled from a Normal $\mathcal{N}(0, 10)$ distribution. We additionally constrained diagonal entries to be negative, to reflect the fact that this is a competitive system where each species exhibits positive growth in monoculture. The species-specific standard errors were sampled from a normal $\mathcal{N}(0, 0.25)$ distribution.

**Results.** The median posterior $B$ values for the *D. pulex* system were:

$$\mathbf{B} = \begin{array}{c} \\ dp \\ cs \\ ma \\ mm \\ sa \end{array} \begin{array}{c} \begin{array}{ccccc} dp & cs & ma & mm & sa \end{array} \\ \begin{pmatrix} 1.19 & -7.66 & -6.28 & -2.04 & -13.23 \\ -0.87 & -0.17 & -0.38 & 2.95 & -0.31 \\ -0.81 & 4.95 & -1.10 & 6.95 & -0.55 \\ -0.94 & 5.66 & -0.55 & -1.03 & -0.59 \\ -1.15 & 6.67 & 0.09 & 10.19 & -1.01 \end{pmatrix} \end{array}$$

And for the *C. dubia* system:

$$\mathbf{B} = \begin{array}{c} \\ cd \\ cs \\ ma \\ mm \\ sa \end{array} \begin{array}{c} \begin{array}{ccccc} cd & cs & ma & mm & sa \end{array} \\ \begin{pmatrix} 3.79 & -16.93 & -9.70 & -1.69 & -4.39 \\ -0.61 & -0.16 & -0.53 & 3.39 & -0.29 \\ -0.54 & 3.38 & -0.89 & 4.39 & -0.51 \\ -0.58 & 5.38 & -0.52 & -1.35 & -0.60 \\ -0.57 & 4.30 & -0.56 & 8.92 & -0.74 \end{pmatrix} \end{array}$$

The full sets of predictions for these two systems are depicted in Figs 5-6. As noted in the main text, this approach is able to recover with high accuracy the median endpoint abundances ($R^2 = 0.94$ for observed vs. predicted) and exhibits strong agreement between the predicted and observed variation in abundances for each species.

In contrast to the plant systems, there are some subsets of species that have a non-negligible probability of extinction (Figs 5-6b). For example, in both systems, there is a moderate probability that two-species assemblage comprising *M. minutum* and the herbivore, and the four-species assemblage comprising an herbivore and all but *C. sorokiniana*, cannot coexist, with the probability that at least one species goes extinct above 50% for both assemblages. These lower probabilities of coexistence are consistent with experimental data—in fact, though all experiments were initialized with all 4 algae species, the endpoints typically contain only a subset of the species, meaning that the remaining species went extinct.

As with the plant system, these results suggest interesting biological features. Once again, the error structure for the coefficients is log-normal, such that we must take the $\log$ of the coefficients to compare them on the correct scale. However, because this system has a mix of positive and negative coefficients, we cannot take the $\log$ (or the $\log$ of the negative values, as with the plants). Thus, here we compare $\log(B_{ij} + 10)$, where the value $10$ is chosen to ensure that $B_{ij} + 10 > 0$, allowing us to then take the $\log$. When we plot these algae-by-algae coefficients from the *D. pulex* system versus those form the *C. dubia* system, we see a very tight relationship (Fig. 3b, main text).

This result suggests that, in general, the competitive effect of one algae on another is independent of the identify of the herbivore—as such the data do not seem to support the existence of strong herbivore-mediated higher-order interactions.

If we are willing to assume Lotka-Volterra dynamics for these systems, then we can draw additional conclusions about the stability and invasibility of these communities. In contrast to the plant systems, we see relatively low evidence of stability, apart from the two-species algal systems comprising *M. arcuatum* and *S. acuminatus* or *C. sorokiniana* and *S. acuminatus*, which exhibit high probability of global stability in both herbivore systems (Figs 5-6c). Otherwise there is sparse evidence for global stability, though some evidence for local stability, which varies depending on the identity of the herbivore. In terms of invasibility, we find that nearly every species can invade into every other community (Figs 5-6d). For the herbivores, this results is obvious; for the algae, this result reflects the fact that the positive off-diagonal terms tend to be large relative to the diagonal, facilitating invasion by rare species. Though these results are speculative due to the fact that these dynamics likely deviate from GLV dynamics to some extent, such information can help design and select future assemblages for analysis.

#### B4 Ciliated protist system — data from Pennekamp *et al.*, 2018

In the previous two examples, we used data in which we had information on the density of the species at the end of the experiment. To contrast our method with time-series trajectory matching, we turn to the data published by Pennekamp *et al.*[8], who grew ciliates on bacteria at different temperatures, recording a detailed time-series in replicate for all experiments.

**Description of the experiment.** Pennekamp and colleagues grew six species of ciliates (*Colpidium striatum*, *Dexiostoma campylum*, *Loxocephalus* sp., *Paramecium caudatum*, *Spirostomum teres* and *Tetrahymena thermophila*) in microcosms incubated at different temperatures (from 15°C to 25°C in steps of two degrees). The ciliates were grown in bacterized cultures containing protist pellets and the bacterium *Serratia fonticola*. Of the 63 possible combinations of presence/absence of each species at time 0, the authors experimentally tested 53—all possible combinations besides the triplets of species, for which they chose at random 10 combinations out of 20. Here, we focus on the five-species subsystem without *Tetrahymena thermophila* (Table S3), as this species did not consistently survive in enough communities to allow estimation of its coefficients.

Table S3: **The five-species protist systems analyzed here, using data from Pennekamp *et al.*[8].**

| Abbrv. | Species |
|---|---|
| co | Colpidium striatum |
| de | Dexiostoma campylum |
| lo | Loxocephalus sp. |
| pa | Paramecium caudatum |
| sp | Spirostomum teres |

Although Pennekamp *et al.* tracked the density of the species for 57 days, the data accompanying the article reports values for only the first 37 days, which were obtained by interpolating the data through a cubic spline (F. Pennekamp, personal communication) and which should match the original data quite closely. To match the initial conditions of the more speciose assemblages, Pennekamp and colleagues shifted the time series for the monocultures so that they started at 20% of the respective carrying capacities. However, this resulted in most monocultures displaying a monotonic decline in abundances (perhaps signaling nutrient depletion), which prevented us from identifying the stationary phase and thus rendered these data unusable for our purposes.

The densities were obtained by analyzing videos of each community recorded using a microscope[9]. Organism counts were then scaled to the volume of the microcosm to estimate the densities. The classification success of this method is excellent for pairs and triplets of species, while at high temperatures, it drops considerably for ensembles of five or six species[9]. We therefore analyzed all the available pairs, triplets and quadruplets which included four of the six species (*Colpidium striatum*, *Loxocephalus* sp., *Paramecium caudatum* and *Spirostomum teres*). The time-series show robust coexistence for all available combinations, with the exception of *Colpidium* and *Paramecium*, and possibly the triplet including also *Spirostomum*: in both cases, *Paramecium* seems to decline toward the end of the experiment (Figure 1). Each experiment was performed in two replicates.

**Choosing the time slice to fit.** As illustrated in detail below, our method does not assume that communities have reached equilibrium, but it does assume that species have passed through the transient phase of their dynamics and settled into some sort of stationary distribution. Similarly, species cannot be declining due to progressive nutrient limitation or inhibition. Because the experiment by Pennekamp *et al.* was not designed specifically for our methodological approach, we must identify a satisfactory time period in which communities have reached a stationary distribution (though not necessarily equilibrium) but have not yet begun to decline. To illustrate how one could identify a satisfactory time point in real-world experimental systems, we calculated the total abundance of each community at each time point, reflecting a readily accessible quantity that can be easily measured in many experimental systems (e.g., using optical density or respiration as a proxy of total biomass). We then searched for the time point around which the total abundance changed as little as possible, suggesting a window in which the species had reached some sort of meta-stationary distribution.

In particular, for each species in each replicate, assemblage, and temperature, we computed

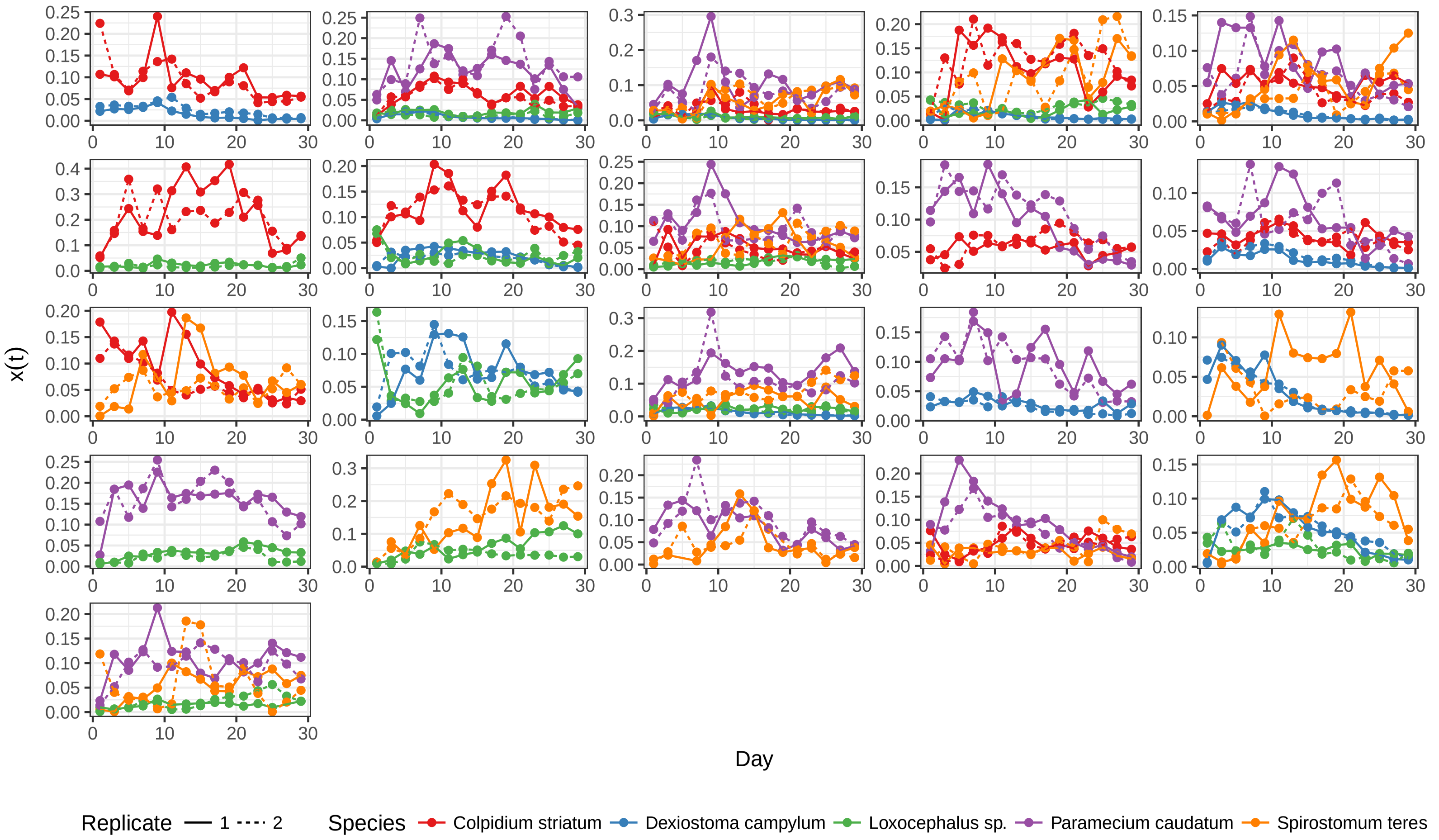

Supplementary Figure 1: Time series for all the available combinations of four species from Pennekamp *et al.*[8]. For each combination, we plot how the density of each species ($y$-axis) changes in time ($x$-axis). The time series are for incubation at 17°C.

$SSQ = \sum_{k \in \{-2,-1,1,2\}} (x(t+k)/x(t) - 1)^2$ which can be interpreted as the (normalized) sum of squares between the density at time $t$ and that at the surrounding four time steps. If species $x$ were at equilibrium, then the value would be zero; if species fluctuate wildly or decline/increase steadily, then this value would be large. We calculated the median value across all species for each time point, with the smallest value indicating when the system is closest to being stationary.

As Fig. 2 demonstrates, biomass was most stable between days 13 and 19 for all temperatures. For simplicity, we selected a consistent 3-day time slice across all temperatures, using days 13, 15, and 17 to capture the average stationary distribution. Thus, each assemblage contained 6 pseudo-replicate measurements, given by the 2 replicates per combinations $\times$ 3 time points.

**Stan fitting details.** To properly scale the data, we first estimated the maximum abundance for each species, calculated as the upper $95^{th}$ percentile across all communities. We then divided each species' abundances in all assemblages by their respective maximum value, such that their self-regulation term $B_{ii}$ would be approximately -1. Under the null hypothesis that species are neutral with respect to each other, the MCMC chains for all temperatures were initialized with $B$ equal to the negative identity matrix. i.e.:

$$\mathbf{B} = \begin{array}{c} \\ co \\ de \\ lo \\ pa \\ sp \end{array} \begin{array}{c} \begin{array}{ccccc} co & de & lo & pa & sp \end{array} \\ \begin{pmatrix} -1.00 & 0 & 0 & 0 & 0 \\ 0 & -1.00 & 0 & 0 & 0 \\ 0 & 0 & -1.00 & 0 & 0 \\ 0 & 0 & 0 & -1.00 & 0 \\ 0 & 0 & 0 & 0 & -1.00 \end{pmatrix} \end{array}$$

We used non-informative priors for the coefficients $B_{ij}$, with each entry sampled from a

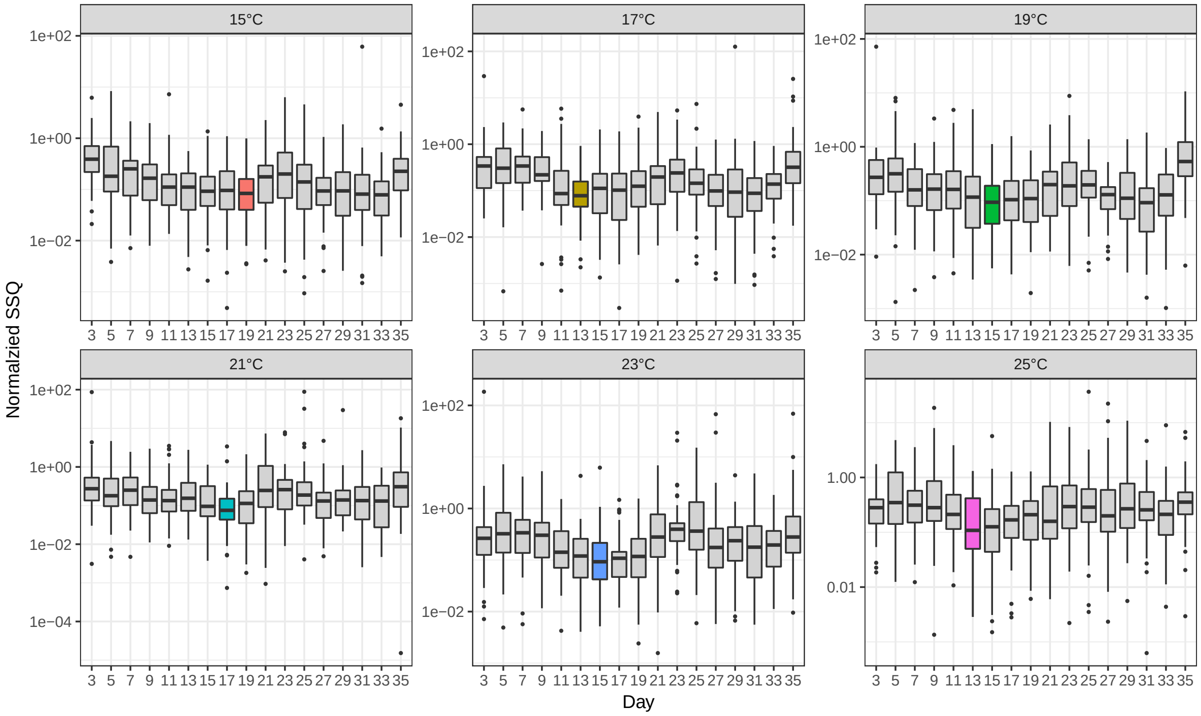

Supplementary Figure 2: The normalized sum of squares of the deviations in total biomass for a 5-day time slice centered at the focal day ($x$-axis). All temperatures exhibit minimal deviations in biomass at days 13-19. The focal day with minimum sum of squares is colored for each temperature. For consistency, we fit all temperatures using days 13, 15, and 17, resulting in six pseudo-replicates for each assemblage.

Normal $\mathcal{N}(0, 10)$ distribution. We additionally constrained diagonal entries to be negative, to reflect the fact that this is a competitive system where each species exhibits positive growth in monoculture. The species-specific standard errors were sampled from a normal $\mathcal{N}(0, 0.25)$ distribution.

**Results.** This method was implemented on the protist system at all six temperatures. The resulting median posterior $B$ values for 15°C is :

$$\mathbf{B} = \begin{array}{c} \\ co \\ de \\ lo \\ pa \\ sp \end{array} \begin{array}{c} \begin{array}{ccccc} co & de & lo & pa & sp \end{array} \\ \begin{pmatrix} -8.38 & -7.17 & 21.21 & -5.35 & -4.17 \\ -5.27 & -22.34 & 14.70 & -4.46 & -6.17 \\ -4.20 & -2.30 & -14.30 & -3.54 & -4.73 \\ -3.73 & -8.99 & 26.05 & -8.23 & -3.69 \\ -1.42 & -6.17 & 4.51 & -1.73 & -14.82 \end{pmatrix} \end{array}$$

And for 17°C:

$$\mathbf{B} = \begin{array}{c} \\ co \\ de \\ lo \\ pa \\ sp \end{array} \begin{array}{c} \begin{array}{ccccc} co & de & lo & pa & sp \end{array} \\ \begin{pmatrix} -8.49 & -6.67 & 15.41 & -5.02 & -4.59 \\ -5.24 & -26.41 & 11.42 & -4.60 & -5.23 \\ -4.85 & -6.45 & -10.11 & -4.23 & -4.42 \\ -2.09 & -6.53 & 17.57 & -8.43 & -4.07 \\ 0.62 & -10.11 & 15.37 & -1.82 & -14.68 \end{pmatrix} \end{array}$$

And for 19°C:

$$\mathbf{B} = \begin{array}{c} \\ co \\ de \\ lo \\ pa \\ sp \end{array} \begin{array}{c} \begin{array}{ccccc} co & de & lo & pa & sp \end{array} \\ \begin{pmatrix} -9.79 & -4.28 & 18.44 & -5.76 & -7.23 \\ -6.93 & -26.43 & 19.31 & -5.27 & -8.39 \\ -4.35 & -9.16 & -8.17 & -5.15 & -5.88 \\ -6.11 & 1.63 & 25.41 & -8.45 & -6.57 \\ -0.90 & 5.25 & 16.84 & -2.12 & -17.07 \end{pmatrix} \end{array}$$

And for 21°C:

$$\mathbf{B} = \begin{array}{c} \\ co \\ de \\ lo \\ pa \\ sp \end{array} \begin{array}{c} \begin{array}{ccccc} co & de & lo & pa & sp \end{array} \\ \begin{pmatrix} -16.64 & -3.36 & 13.28 & -8.76 & -12.80 \\ -11.70 & -34.12 & 17.19 & -8.33 & -14.99 \\ -5.48 & -9.99 & -7.13 & -6.81 & -11.11 \\ -9.81 & 8.72 & 19.92 & -13.57 & -9.44 \\ -9.91 & -2.08 & 17.63 & -6.87 & -22.63 \end{pmatrix} \end{array}$$

And for 23°C:

$$\mathbf{B} = \begin{array}{c} \\ co \\ de \\ lo \\ pa \\ sp \end{array} \begin{array}{c} \begin{array}{ccccc} co & de & lo & pa & sp \end{array} \\ \begin{pmatrix} -19.29 & -8.92 & 12.35 & -10.53 & -8.77 \\ -11.12 & -40.79 & 15.27 & -10.47 & -10.65 \\ -9.33 & -14.80 & -7.08 & -9.87 & -8.54 \\ -2.47 & 7.36 & 30.90 & -17.95 & -6.54 \\ -1.32 & -9.49 & 24.06 & -7.45 & -22.55 \end{pmatrix} \end{array}$$

And for 25°C:

$$
\mathbf{B} = \begin{array}{c} \\ co \\ de \\ lo \\ pa \\ sp \end{array} \begin{array}{c} \begin{array}{ccccc} co & de & lo & pa & sp \end{array} \\ \begin{pmatrix} -29.31 & -8.71 & 4.82 & -13.55 & -10.00 \\ -15.45 & -41.82 & 7.47 & -13.90 & -10.37 \\ -4.88 & -22.74 & -7.97 & -12.28 & -8.86 \\ -13.18 & 10.41 & 8.64 & -18.53 & -7.50 \\ -21.38 & 11.77 & 17.92 & -14.08 & -15.93 \end{pmatrix} \end{array}
$$

The full sets of predictions for these six temperature are depicted in Figs. 9-12. As noted in the main text, this approach is able to predict with high accuracy the median endpoint abundances ($R^2 = 0.92$ for observed vs. predicted), and exhibits strong agreement between the predicted and observed variation in abundances for each species. All temperatures exhibited high probability of coexistence for all subsets (Figs. 9-12b), consistent with few species going extinct across the time-series replicates (e.g., Fig. 1), although the four-species assemblage comprising *C. striatum, D. campylum, P. caudatum* and *S. teres* has a nontrivial risk of extinction for 19, 21, and 23$^o$C.

As shown in the main text, we can compare the structure of $B$ across the temperature gradient (Fig. 4). Doing so reveals consistent changes in the effect that species have on each other, suggesting that the entries of the $B$ matrices are not arbitrary, but in fact capture some meaningful biology of the system. These patterns could help inform future ecological hypotheses and design experiments to directly test the mechanisms underlying these relationships (e.g., *Loxocephalus* sp. has a large beneficial effect on the other species, suggesting some form of "apparent predation" not otherwise evident from the data). Such results should be interpreted cautiously, however, with any mechanistic or dynamical inference requiring additional experimentation. From a purely methodological perspective, these findings demonstrate that this method yields relationships that are ecologically consistent across assemblages and temperatures.

If we are willing to assume Lotka-Volterra dynamics for these assemblages, then we can make additional conclusions about this system. The protists exhibited complex stability results across all temperatures (Figs. 9-12c): every community across all temperatures exhibited a high likelihood of local stability, but varied likelihood of global stability. In general, most of the pairwise interactions exhibited high probability of global stability, which became less likely at higher richness levels; though there are some consistent patterns, such as the four-species community comprising *C. striatum, D. campylum, P. caudatum*, and *Sp. teres* showing high probability of global stability at all temperatures. Similarly, likelihood of invasibility is near 100% for all communities across all temperatures (Figs. 9-12d). Though these results are speculative due to the fact that these dynamics likely deviate from GLV dynamics to some extent, such information can help design and select future assemblages for analysis.

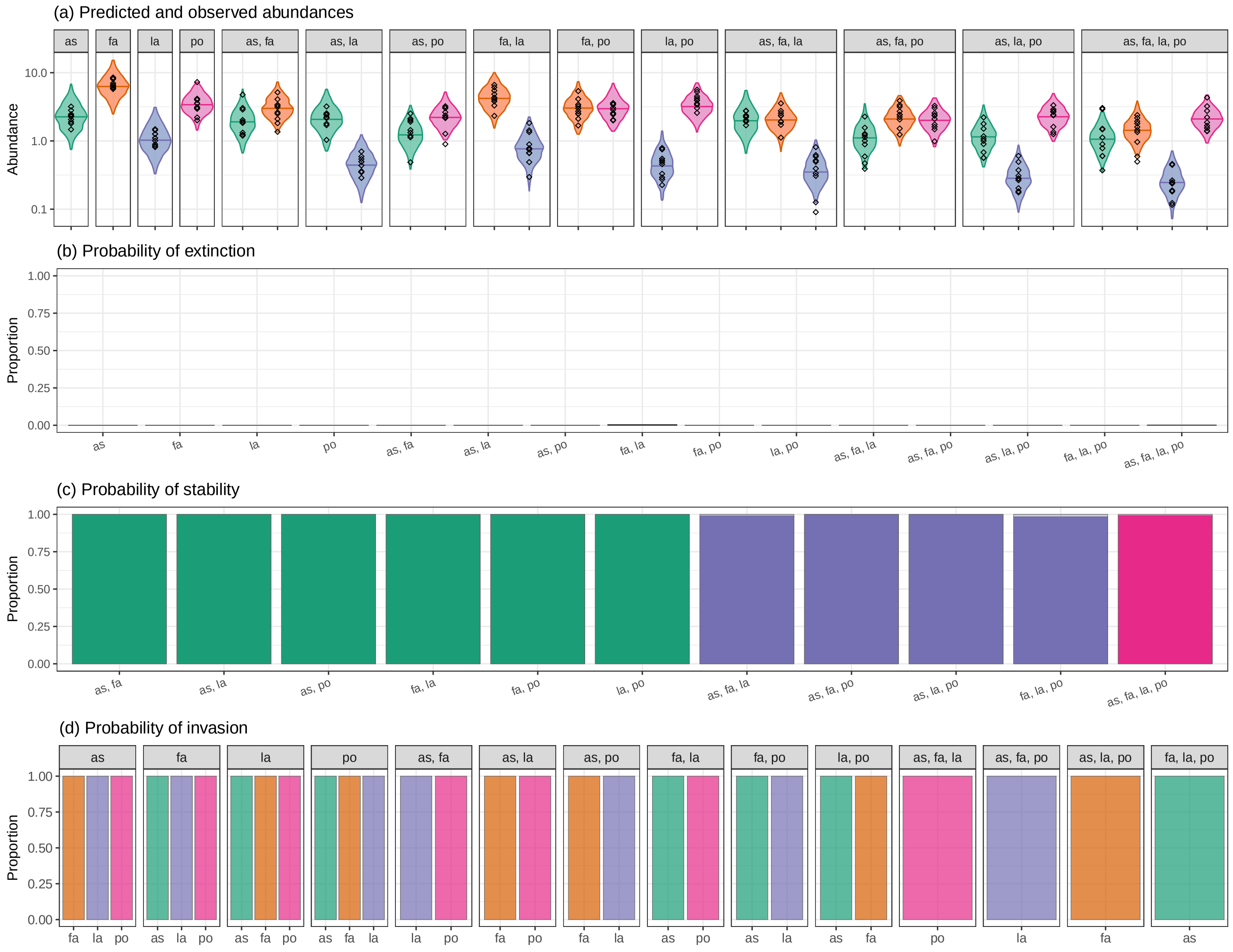

Supplementary Figure 3: **The results for the native plant system.** **(a)** The violin plot gives the predicted posterior distribution of each species' abundance in the community, with the black diamonds showing the observed endpoint abundances. **(b)** The probability of extinction, calculated as the proportion of posterior $B$'s that resulted in a non-feasible endpoint, indicating that these species cannot coexist together (observed endpoints in green, unobserved in orange). **(c)** The probability of stability, calculated as the proportion of posterior $B$'s that exhibited approximate local stability, under the assumption of equal growth rates (light shaded), or global stability, with no assumption on the growth rates (dark shaded). Colors denote the size of the community. **(d)** The probability of invasion, calculated as the proportion of posterior $B$'s for which the focal species (x-axis) was able to invade into the community. Note that (b) and (c) were calculated by removing the endpoint of interest from the analysis, thus testing out-of-fit predictions; and that (c) and (d) assume GLV dynamics around the feasible fixed point, and thus are speculative.

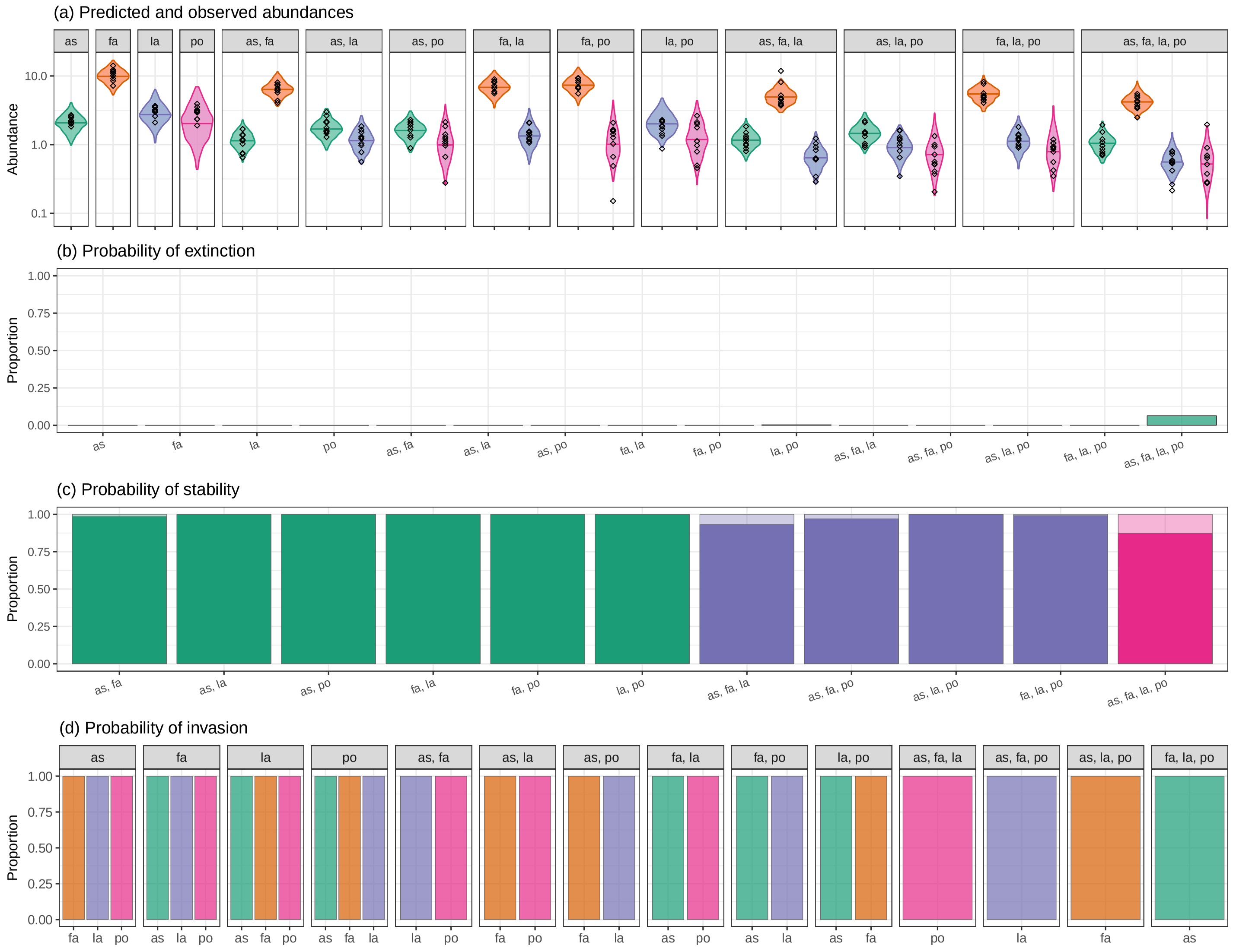

Supplementary Figure 4: **The results for the nonnative plant system.** **(a)** The violin plot gives the predicted posterior distribution of each species' abundance in the community, with the black diamonds showing the observed endpoint abundances. **(b)** The probability of extinction, calculated as the proportion of posterior $B$'s that resulted in a non-feasible endpoint, indicating that these species cannot coexist together (observed endpoints in green, unobserved in orange). **(c)** The probability of stability, calculated as the proportion of posterior $B$'s that exhibited approximate local stability, under the assumption of equal growth rates (light shaded), or global stability, with no assumption on the growth rates (dark shaded). Colors denote the size of the community. **(d)** The probability of invasion, calculated as the proportion of posterior $B$'s for which the focal species (x-axis) was able to invade into the community. Note that (b) and (c) were calculated by removing the endpoint of interest from the analysis, thus testing out-of-fit predictions; and that (c) and (d) assume GLV dynamics around the feasible fixed point, and thus are speculative.

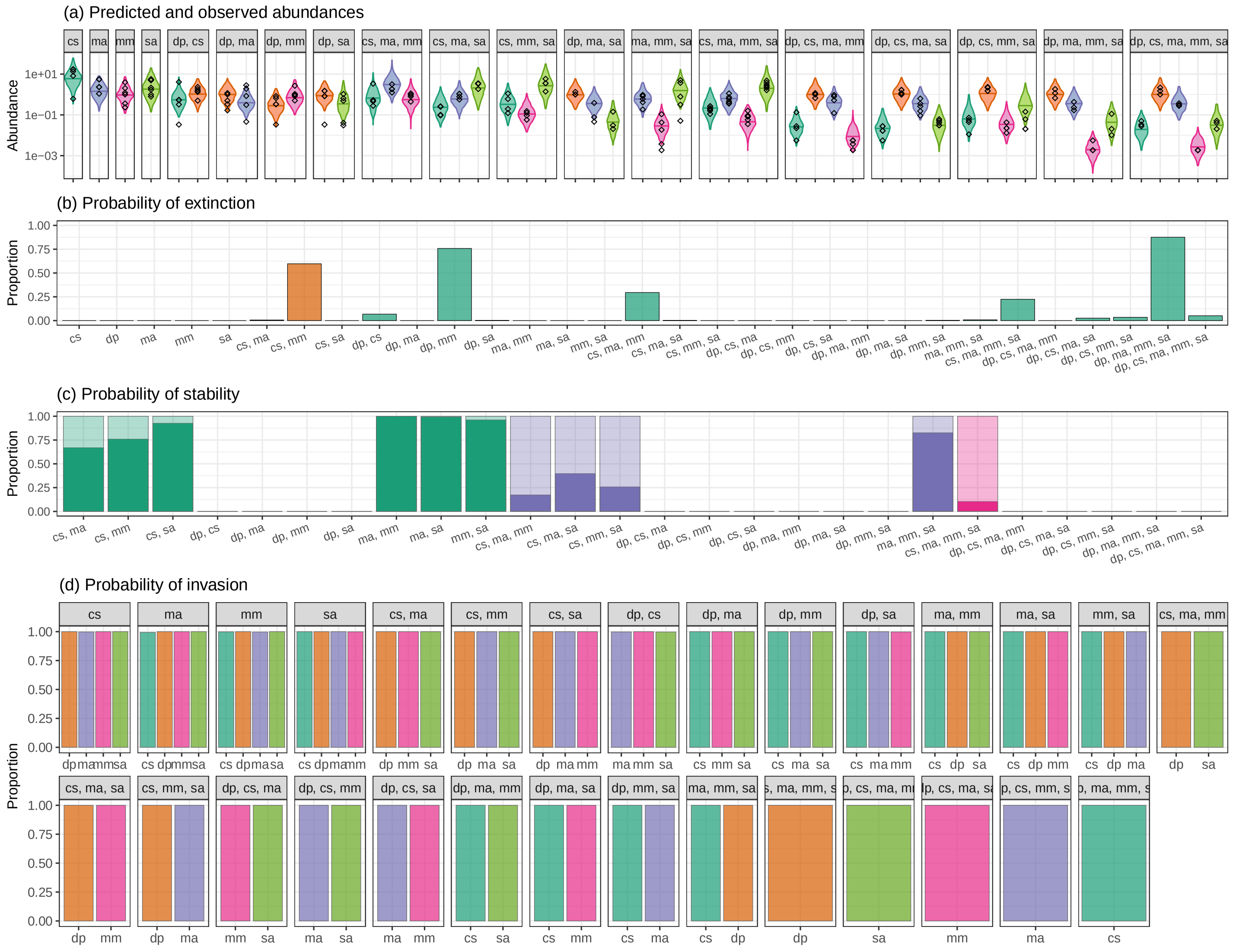

Supplementary Figure 5: **The results for the herbivore-algae system containing *D. pulex*.** **(a)** The violin plot gives the predicted posterior distribution of each species' abundance in the community, with the black diamonds showing the observed endpoint abundances. **(b)** The probability of extinction, calculated as the proportion of posterior $B$'s that resulted in a non-feasible endpoint, indicating that these species cannot coexist together (observed endpoints in green, unobserved in orange). **(c)** The probability of stability, calculated as the proportion of posterior $B$'s that exhibited approximate local stability, under the assumption of equal growth rates (light shaded), or global stability, with no assumption on the growth rates (dark shaded). Colors denote the size of the community. **(d)** The probability of invasion, calculated as the proportion of posterior $B$'s for which the focal species (x-axis) was able to invade into the community. Note that (b) and (c) were calculated by removing the endpoint of interest from the analysis, thus testing out-of-fit predictions; and that (c) and (d) assume GLV dynamics around the feasible fixed point, and thus are speculative.

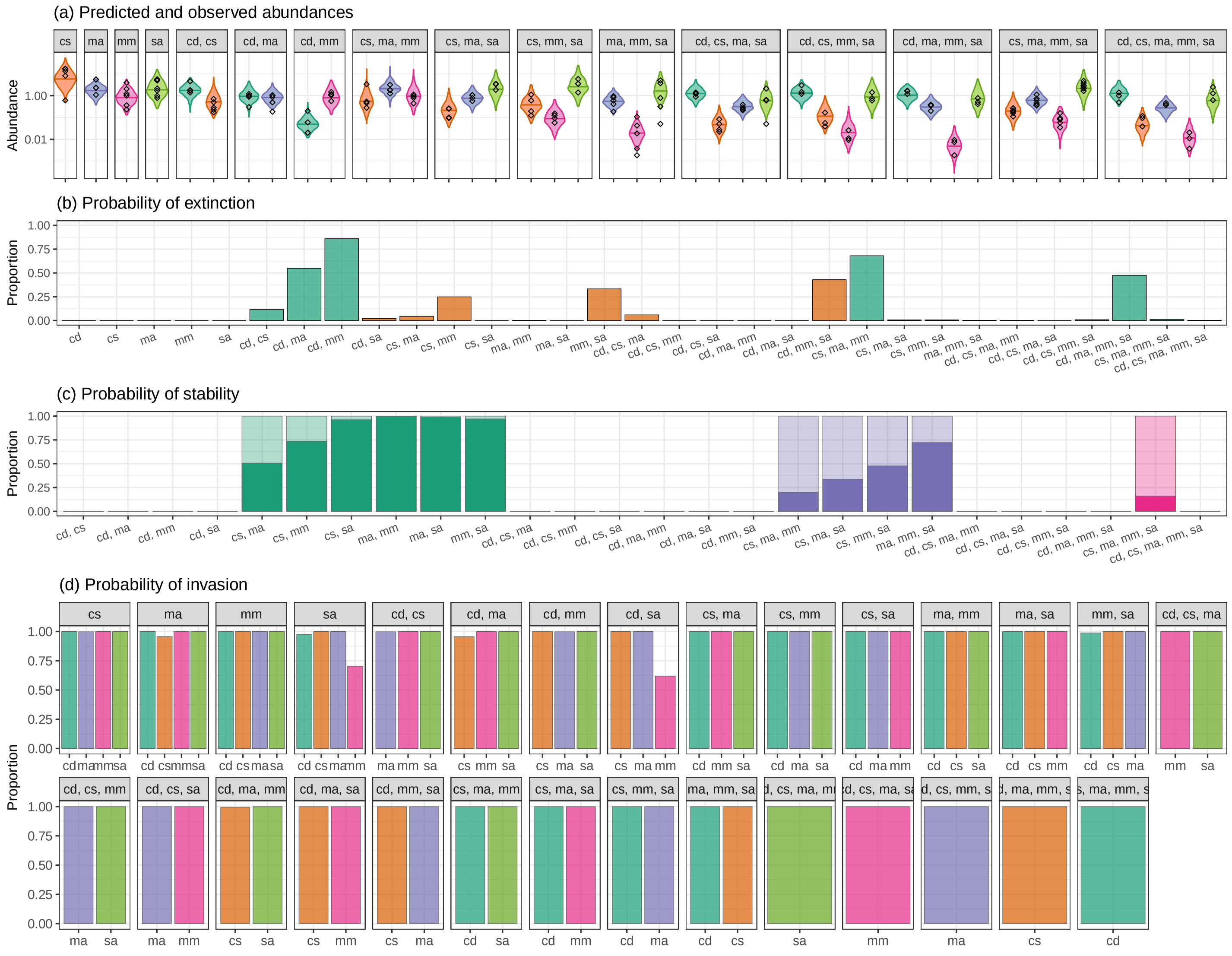

Supplementary Figure 6: **The results for the herbivore-algae system containing *C. dubia*.** **(a)** The violin plot gives the predicted posterior distribution of each species' abundance in the community, with the black diamonds showing the observed endpoint abundances. **(b)** The probability of extinction, calculated as the proportion of posterior $B$'s that resulted in a non-feasible endpoint, indicating that these species cannot coexist together (observed endpoints in green, unobserved in orange). **(c)** The probability of stability, calculated as the proportion of posterior $B$'s that exhibited approximate local stability, under the assumption of equal growth rates (light shaded), or global stability, with no assumption on the growth rates (dark shaded). Colors denote the size of the community. **(d)** The probability of invasion, calculated as the proportion of posterior $B$'s for which the focal species (x-axis) was able to invade into the community. Note that (b) and (c) were calculated by removing the endpoint of interest from the analysis, thus testing out-of-fit predictions; and that (c) and (d) assume GLV dynamics around the feasible fixed point, and thus are speculative.

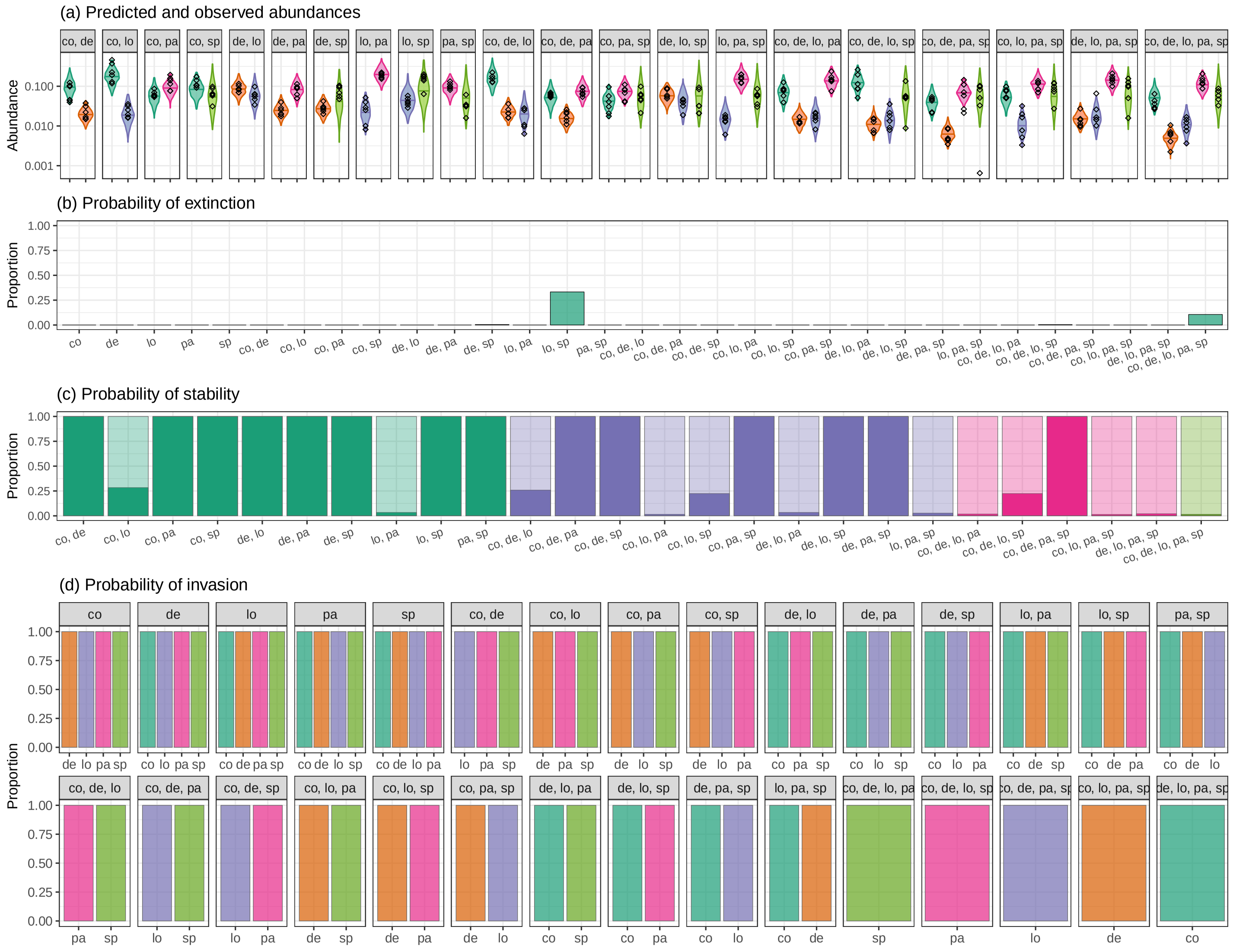

Supplementary Figure 7: **The results for the protist system at 15$^o$C.** **(a)** The violin plot gives the predicted posterior distribution of each species' abundance in the community, with the black diamonds showing the observed endpoint abundances. **(b)** The probability of extinction, calculated as the proportion of posterior $B$'s that resulted in a non-feasible endpoint, indicating that these species cannot coexist together (observed endpoints in green, unobserved in orange). **(c)** The probability of stability, calculated as the proportion of posterior $B$'s that exhibited approximate local stability, under the assumption of equal growth rates (light shaded), or global stability, with no assumption on the growth rates (dark shaded). Colors denote the size of the community. **(d)** The probability of invasion, calculated as the proportion of posterior $B$'s for which the focal species (x-axis) was able to invade into the community. Note that (b) and (c) were calculated by removing the endpoint of interest from the analysis, thus testing out-of-fit predictions; and that (c) and (d) assume GLV dynamics around the feasible fixed point, and thus are speculative.

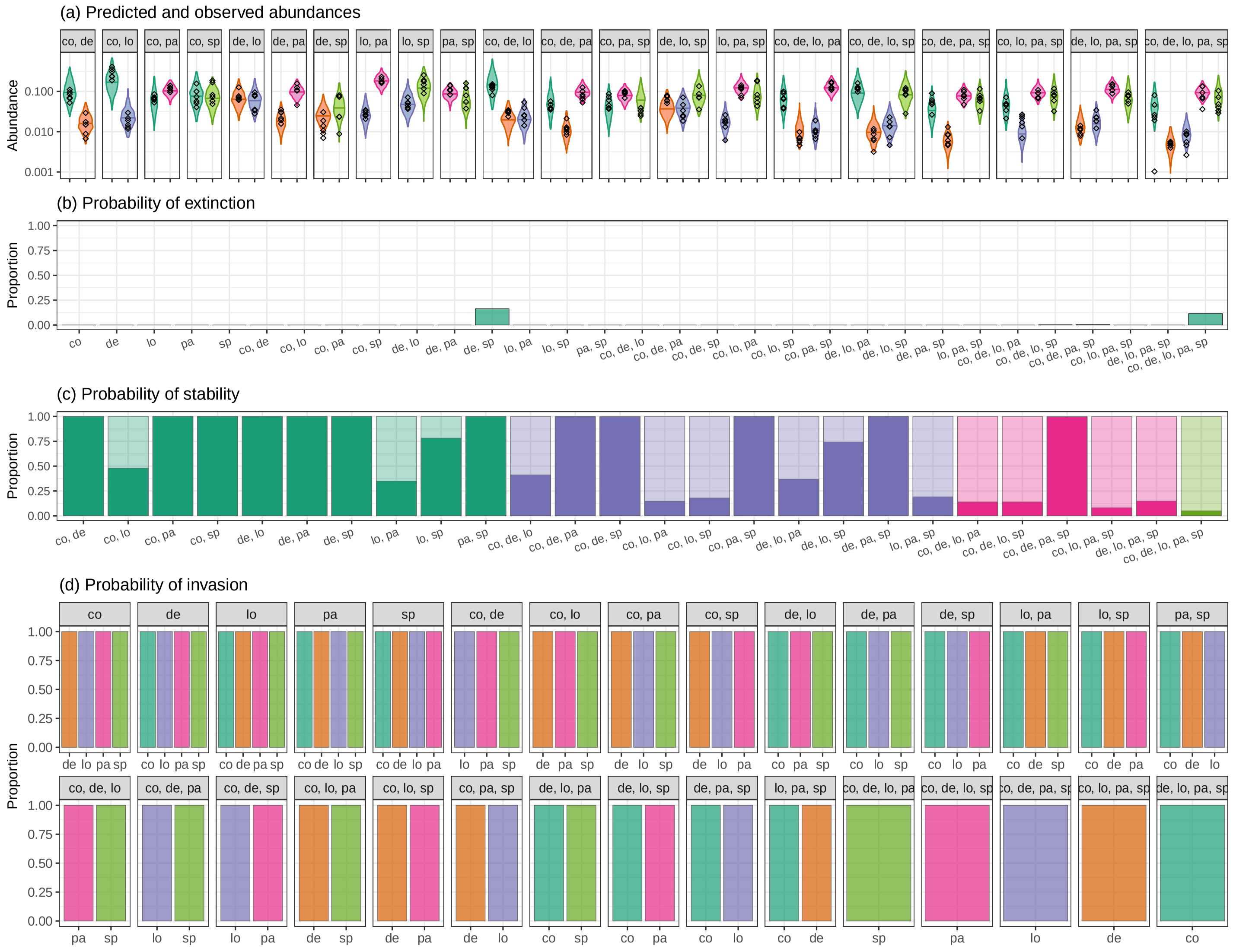

Supplementary Figure 8: **The results for the protist system at 17$^o$C.** **(a)** The violin plot gives the predicted posterior distribution of each species' abundance in the community, with the black diamonds showing the observed endpoint abundances. **(b)** The probability of extinction, calculated as the proportion of posterior $B$'s that resulted in a non-feasible endpoint, indicating that these species cannot coexist together (observed endpoints in green, unobserved in orange). **(c)** The probability of stability, calculated as the proportion of posterior $B$'s that exhibited approximate local stability, under the assumption of equal growth rates (light shaded), or global stability, with no assumption on the growth rates (dark shaded). Colors denote the size of the community. **(d)** The probability of invasion, calculated as the proportion of posterior $B$'s for which the focal species (x-axis) was able to invade into the community. Note that (b) and (c) were calculated by removing the endpoint of interest from the analysis, thus testing out-of-fit predictions; and that (c) and (d) assume GLV dynamics around the feasible fixed point, and thus are speculative.

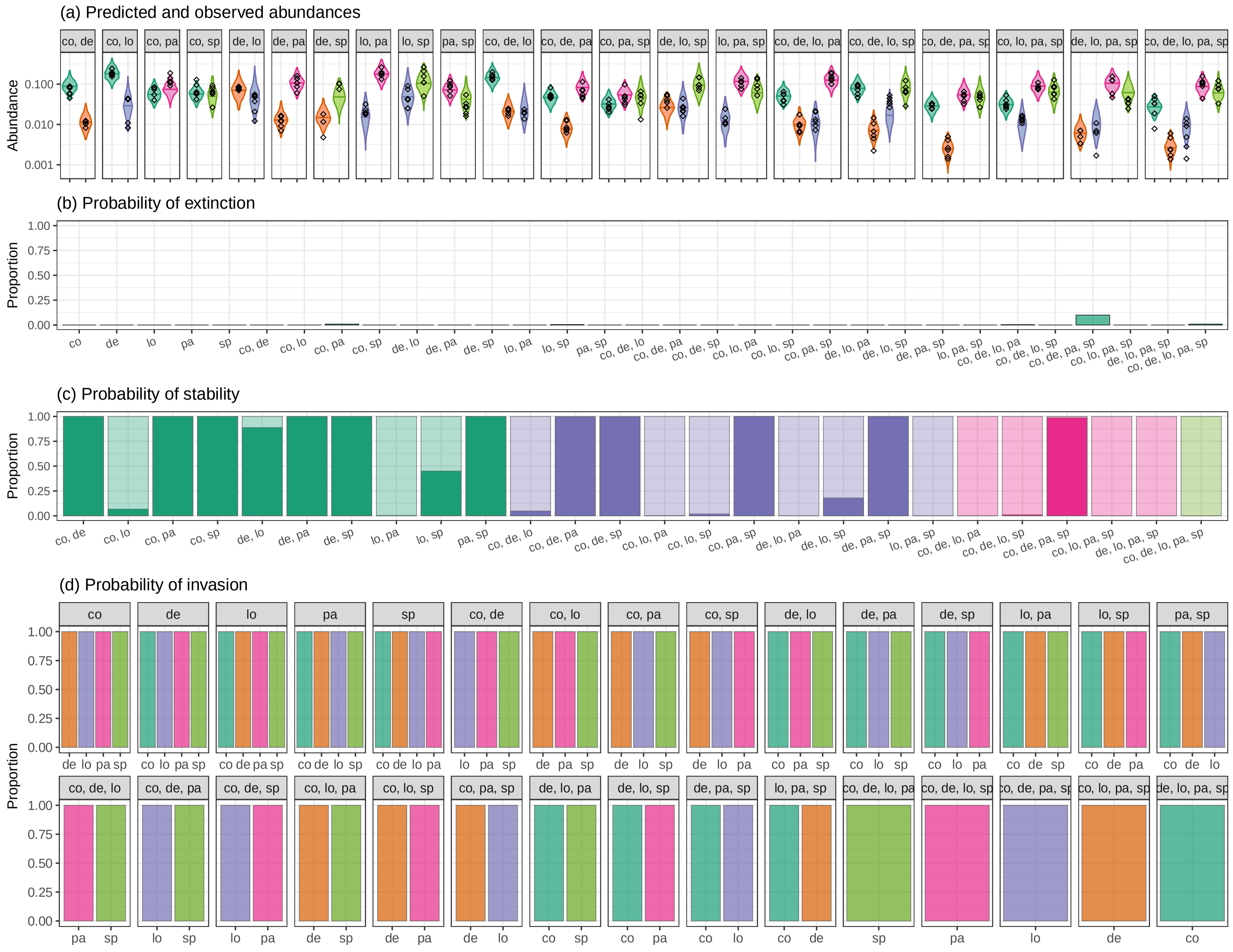

Supplementary Figure 9: **The results for the protist system at 19$^o$C.** **(a)** The violin plot gives the predicted posterior distribution of each species' abundance in the community, with the black diamonds showing the observed endpoint abundances. **(b)** The probability of extinction, calculated as the proportion of posterior $B$'s that resulted in a non-feasible endpoint, indicating that these species cannot coexist together (observed endpoints in green, unobserved in orange). **(c)** The probability of stability, calculated as the proportion of posterior $B$'s that exhibited approximate local stability, under the assumption of equal growth rates (light shaded), or global stability, with no assumption on the growth rates (dark shaded). Colors denote the size of the community. **(d)** The probability of invasion, calculated as the proportion of posterior $B$'s for which the focal species (x-axis) was able to invade into the community. Note that (b) and (c) were calculated by removing the endpoint of interest from the analysis, thus testing out-of-fit predictions; and that (c) and (d) assume GLV dynamics around the feasible fixed point, and thus are speculative.

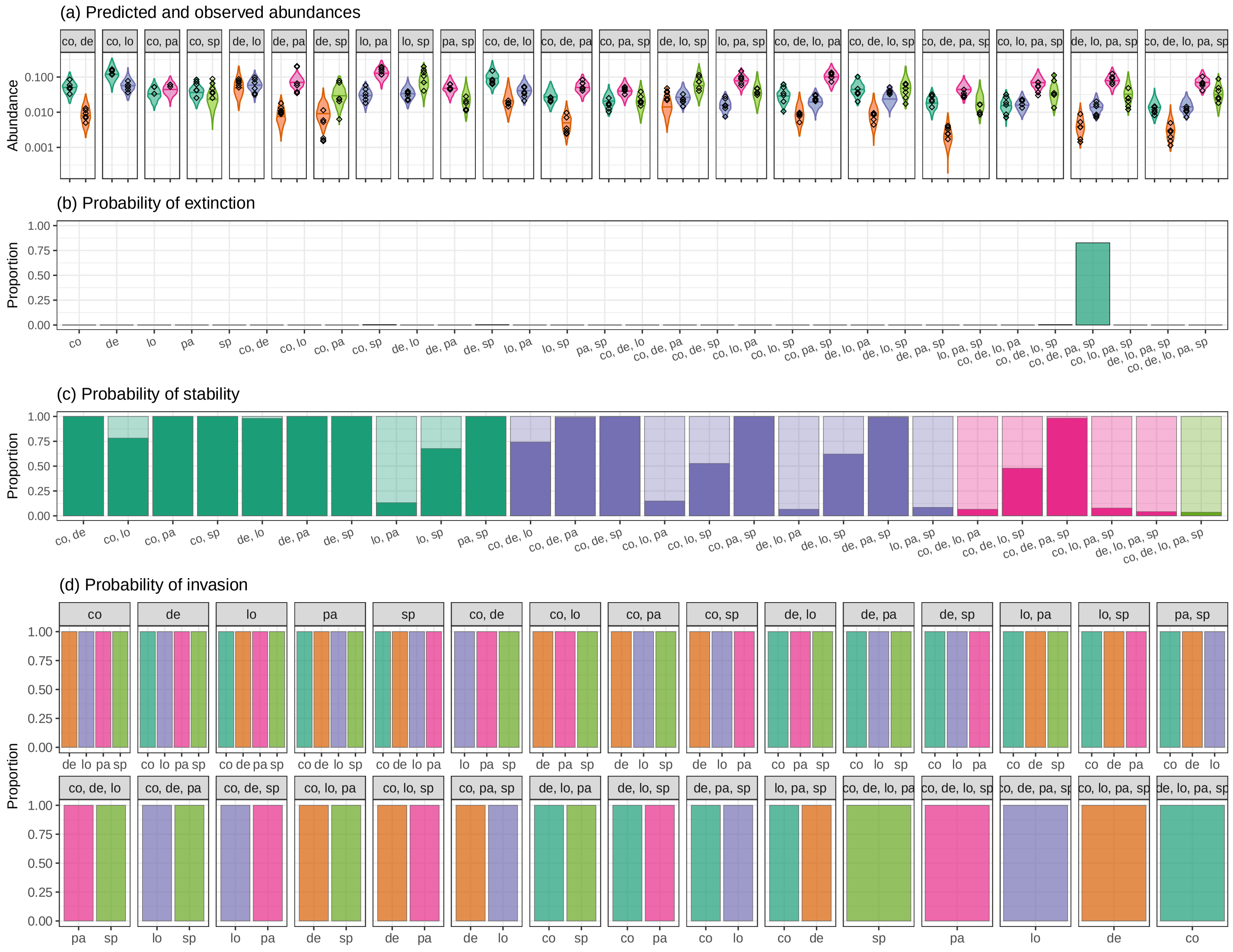

Supplementary Figure 10: **The results for the protist system at 21$^o$C.** **(a)** The violin plot gives the predicted posterior distribution of each species' abundance in the community, with the black diamonds showing the observed endpoint abundances. **(b)** The probability of extinction, calculated as the proportion of posterior $B$'s that resulted in a non-feasible endpoint, indicating that these species cannot coexist together (observed endpoints in green, unobserved in orange). **(c)** The probability of stability, calculated as the proportion of posterior $B$'s that exhibited approximate local stability, under the assumption of equal growth rates (light shaded), or global stability, with no assumption on the growth rates (dark shaded). Colors denote the size of the community. **(d)** The probability of invasion, calculated as the proportion of posterior $B$'s for which the focal species (x-axis) was able to invade into the community. Note that (b) and (c) were calculated by removing the endpoint of interest from the analysis, thus testing out-of-fit predictions; and that (c) and (d) assume GLV dynamics around the feasible fixed point, and thus are speculative.

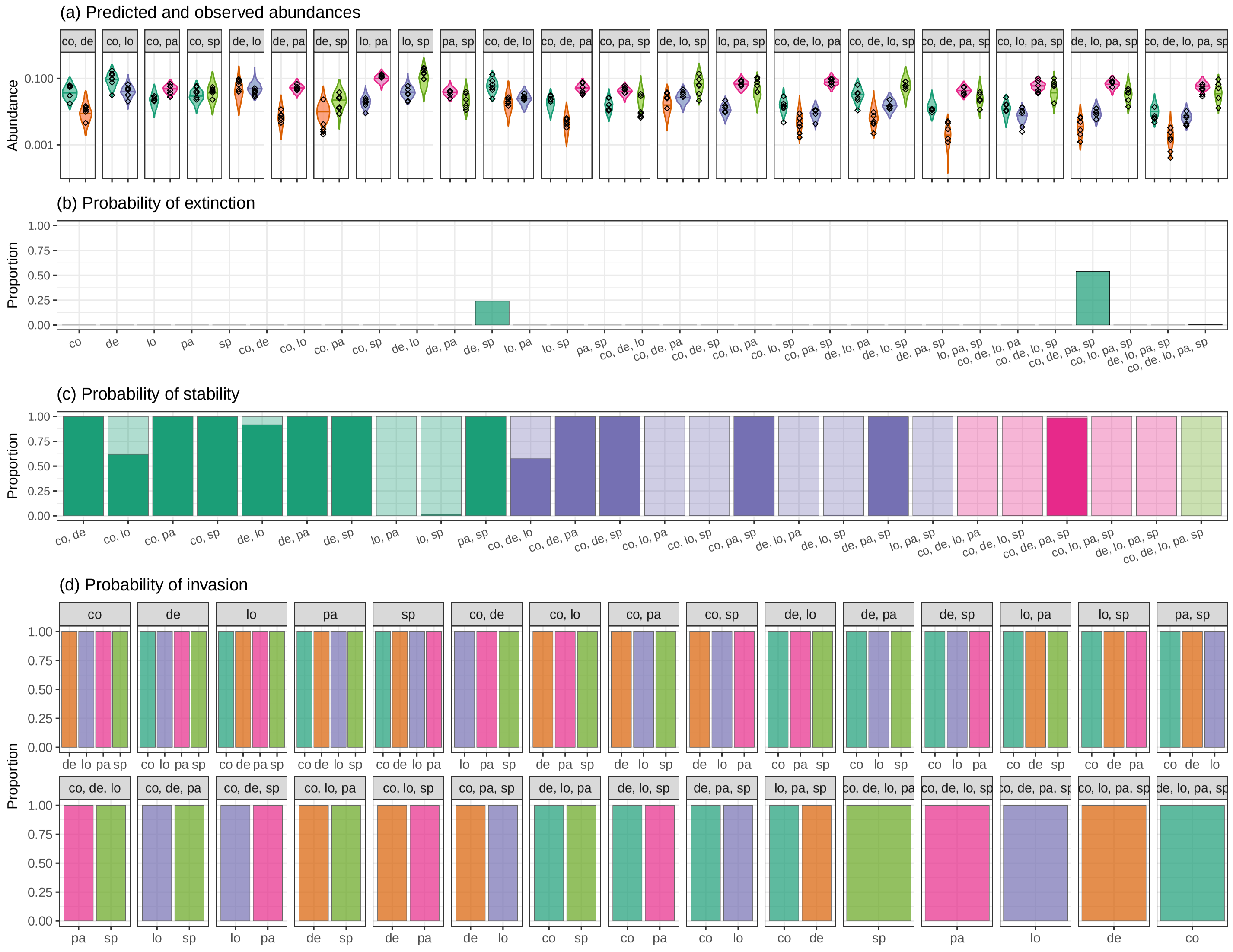

Supplementary Figure 11: **The results for the protist system at 23$^o$C.** **(a)** The violin plot gives the predicted posterior distribution of each species' abundance in the community, with the black diamonds showing the observed endpoint abundances. **(b)** The probability of extinction, calculated as the proportion of posterior $B$'s that resulted in a non-feasible endpoint, indicating that these species cannot coexist together (observed endpoints in green, unobserved in orange). **(c)** The probability of stability, calculated as the proportion of posterior $B$'s that exhibited approximate local stability, under the assumption of equal growth rates (light shaded), or global stability, with no assumption on the growth rates (dark shaded). Colors denote the size of the community. **(d)** The probability of invasion, calculated as the proportion of posterior $B$'s for which the focal species (x-axis) was able to invade into the community. Note that (b) and (c) were calculated by removing the endpoint of interest from the analysis, thus testing out-of-fit predictions; and that (c) and (d) assume GLV dynamics around the feasible fixed point, and thus are speculative.

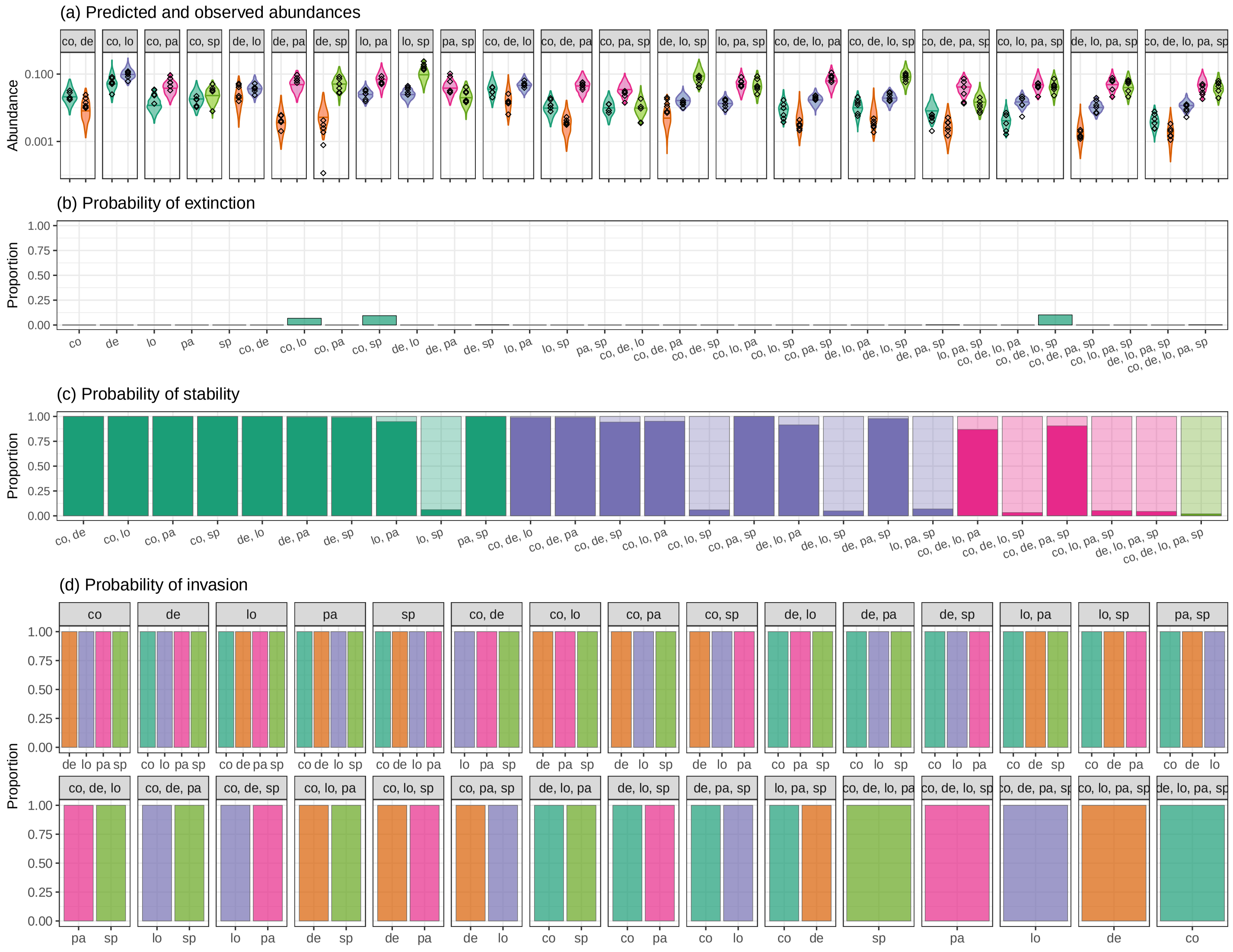

Supplementary Figure 12: **The results for the protist system at 25°C.** **(a)** The violin plot gives the predicted posterior distribution of each species' abundance in the community, with the black diamonds showing the observed endpoint abundances. **(b)** The probability of extinction, calculated as the proportion of posterior $B$'s that resulted in a non-feasible endpoint, indicating that these species cannot coexist together (observed endpoints in green, unobserved in orange). **(c)** The probability of stability, calculated as the proportion of posterior $B$'s that exhibited approximate local stability, under the assumption of equal growth rates (light shaded), or global stability, with no assumption on the growth rates (dark shaded). Colors denote the size of the community. **(d)** The probability of invasion, calculated as the proportion of posterior $B$'s for which the focal species (x-axis) was able to invade into the community. Note that (b) and (c) were calculated by removing the endpoint of interest from the analysis, thus testing out-of-fit predictions; and that (c) and (d) assume GLV dynamics around the feasible fixed point, and thus are speculative.

### C Time-series analysis

Because the data of Pennekamp *et al.* contain the full time series for each assemblage, we can compare our method with the traditional approach of parameterizing a dynamical model (in this case, the standard Lotka-Volterra model, parameterized using trajectory matching) in order to predict the evolution of the system under new conditions (in this case, with different species composition). Since several of the species begin to decline in abundance at higher temperatures and beyond day 17 (possibly due to thermal stress and/or nutrient limitation), we fit the time series for 15° C, using days 1-17 so as to match the analysis in the main text.

#### C1 Model

The dynamical model we use is the standard Lotka-Volterra model of competition. The rate of change for each species $i = 1, \dots, 5$ is a function of its intrinsic growth rate, $r_i$, scaling the set of interactions $\frac{A_{ij}}{r_i} = \tilde{A}_{ij}$, for $j = 1, \dots, 5$, where $A_{ij}$ give the per-capita effect of species $j$ on species $i$. If we let $x_i(t) = x_i$ be the abundance of species at time $t$, then the dynamics of this system is given by:

$$\frac{\mathrm{d}x_i}{\mathrm{d}t} = x_i r_i \left(1 + \sum_{i=1}^{5} \tilde{A}_{ij} x_j\right) \quad i = 1, \dots, 5 \ , \tag{10}$$

which can be written in matrix notation as:

$$\frac{\mathrm{d}x}{\mathrm{d}t} = D(x \circ r)(1 + \tilde{A}x) \ , \tag{11}$$

where $x = (x_1, \dots, x_5)^t$, $r = (r_1, \dots, r_5)^t$, $\tilde{A}$ is the matrix of dimension $5 \times 5$ comprised of the $\tilde{A}_{ij}$, and where $D(x \circ r)$ is a diagonal matrix with the vector $x \circ r$ on the diagonal. Here we use $\circ$ to denote the Hadamard (entrywise) product. Note that the standard Lotka-Volterra interaction matrix can be calculated by $A = D(r)\tilde{A}$.

To parameterize this model we thus need to estimate the entries of $r$ and $\tilde{A}$. Once we have these values, then for any subset of species, $s$, comprising $k \leq 5$ species, we can integrate the temporal dynamics:

$$\frac{\mathrm{d}x^{(s)}}{\mathrm{d}t} = D(x^{(s)} \circ r^{(s)})(1 + \tilde{A}^{(s)} x^{s}) \ , \tag{12}$$

where $x^{(s)}$ and $r^{(s)}$ are the vectors of length $k$ corresponding to the species in $s$, and $\tilde{A}^{(s)}$ is the $k \times k$ submatrix comprising the rows and columns corresponding to the species in $s$. The equilibrium abundances for this subset $x^{\star(s)}$ can be calculated as:

$$x^{\star(s)} = -\tilde{A}^{(s)-1} 1 \ , \tag{13}$$

which we can compare to the observed abundances, as in the main text.

#### C2 Fitting procedure

To estimate $r$ and $\tilde{A}$ we implemented standard trajectory matching by minimizing the log sum-of-squares (SSQ) between the predicted and observed time series. The basic approach was to: (1) initialize $r$ and $\tilde{A}$ with some starting values, (2) use these parameters to project the 17-day time series by integrating Eqn. 12 for each observed subset of species, with initial abundances set to the observed abundances on day 1, (3) calculate the log-SSQ deviation between the observed and predicted observations by summing the log deviations over all time points (days $1, 3, 5, \ldots, 17$) for each species in each community, and (4) repeat, using a standard search algorithm to find the values of $r$ and $\tilde{A}$ that minimize the log-SSQ.

The detailed fitting procedure is as follows:

1. We initialized $\tilde{A}$ to be a diagonal matrix, where $\tilde{A}_{ii}$ is taken to the negative inverse of the maximum abundance of species $i$, that is $\tilde{A}_{ii} = -1/\max(x_i)$, where $\max(x_i)$ was

calculated as the upper $95^{th}$ percentile abundance across all observations. As with the Bayesian approach, this initial $\tilde{A}$ thus assumes species are neutral with respect to each other, and that variation in abundances across days and communities is purely a result of each species fluctuating about its carrying capacity. We initialized $r = (1, 1, 1, 1, 1)^t$ as a non-informative initial guess for the growth rates.

2. For each of the 21 unique species combinations (Fig. 1) we separately integrated the temporal dynamics of each replicate using the current estimate of $\tilde{A}$ and $r$. For each species in each replicate we set its initial abundance equal to the first observed non-zero abundance in the time series, as some species had zero abundance at day 1. For each species $i$ in assemblage $k$ we thus obtained an estimate of its abundance for each day of each time series, $\widehat{x}_i^{(k)}(t)$ for $t = 1, 3, \ldots, 17$, which we can then compare to the corresponding observed abundances, $x_i^{(k)}(t)$ for $t = 1, 3, \ldots, 17$.

3. We calculated the log sum of squares by summing the squared deviation across each assemblage $k = 1, \ldots, 21$, across each time points $t = 1, 3, \ldots, 17$, across each replicate, and across each species $i$ that was present in assemblage $k$ (note that the replicates are dropped from this summation to simplify the notation).

$$SSQ = \sum_{\substack{t=1 \\ t\ odd}}^{17} \sum_{k=1}^{21} \sum_{i \in E_k} (\log x_i^{(k)}(t) - \log \widehat{x}_i^{(k)}(t))^2 \tag{14}$$

If species $i$ was present in endpoint $k$ but its predicted abundance at time $t$ was zero (it went extinct) or if its abundance had grown to infinity, we set $\widehat{x}_i^{(k)}(t) = 10^{100}$ thus ensuring a large penalization for this outcome. Using the log sum-of-squares rather than the raw sum-of-squares accounts for the fact that the errors are proportional to the mean abundances.

4. To find $\tilde{A}$ and $r$ that minimize the sum of squares, we used the `optim` function in $R$, using the default "Nelder-Mead" search algorithm. We constrained the growth rates to be positive and the diagonals of $\tilde{A}$ to be negative, consistent with a species-specific carrying capacity, but otherwise we places no constraints on the entries. Tolerance for convergence was set to machine epsilon.

This approach results in a matrix $\tilde{A}$ and vector $r$ that yield dynamics which most closely track the observed time-series data across all species, replicates, and assemblages Fig. 13.

#### C3 Results

The estimates of $r$ and $\tilde{A}$ that minimize the log-sum-of-squares are:

$$\mathbf{r} = \begin{matrix} co \\ de \\ lo \\ pa \\ sp \end{matrix} \begin{pmatrix} 3.7 \\ 1.2 \\ 72.4 \\ 29.7 \\ 1.1 \end{pmatrix} \qquad \mathbf{B} = \begin{matrix} \\ co \\ de \\ lo \\ pa \\ sp \end{matrix} \begin{matrix} co & de & lo & pa & sp \\ \end{matrix} \begin{pmatrix} -8.1 & -5.0 & 35.7 & -5.1 & -2.7 \\ -4.8 & -16.1 & 19.9 & -4.0 & -8.1 \\ -2.5 & -1.8 & -24.7 & -3.4 & -4.2 \\ -2.8 & -2.3 & 41.1 & -8.2 & -2.6 \\ -0.8 & -3.4 & 1.1 & -1.6 & -20.6 \end{pmatrix}$$

resulting in the predicted time series shown in Fig. 13.

When we compare this $\tilde{A}$ matrix to the corresponding $B$ matrix obtained via the method developed here, we see that the two matrices are highly correlated (Fig. 14). This demonstrates two things. First, it suggests that the proposed method is essentially equivalent to finding an effective Lotka-Volterra model with equilibria that map onto the observed endpoints. Second, it shows that the endpoints alone carry essentially the same information as the $\tilde{A}$ matrix one would obtain by considering the full dynamics.

The trajectory matching procedure essentially fits a smooth curve through the observed time series, ignoring much of the fine-scale dynamics of the system (Fig. 13). Some of the variation in observed dynamics may simply be due to sample error and stochastic noise; yet the dynamics of this system almost certainly deviate from Lotka-Volterra in nature, such that the best-fitting parametrization is only able to approximate the coarse dynamics of this system. Such results highlight a key challenge in fitting time-series data: the quality of the the fit is limited by the appropriateness of the chosen model.

The time-series fit does a satisfactory job predicting the endpoints of the system (Fig. 15), yet the quality of fit is substantially lower than the prediction made using the $B$ matrix as estimated via the proposed method. The dynamical model weights each time point equally, sacrificing predictive accuracy of the endpoint for predictive accuracy across the time series. In contrast, the proposed method focuses solely on recovering the endpoints of the system, at the cost of a complete inability to predict the dynamics. Because the method is agnostic about the dynamics of the system, it is less prone to model misspecification. In doing so, it is able to predict the outcomes with higher accuracy, requiring many fewer sampling points than traditional time-series analysis.

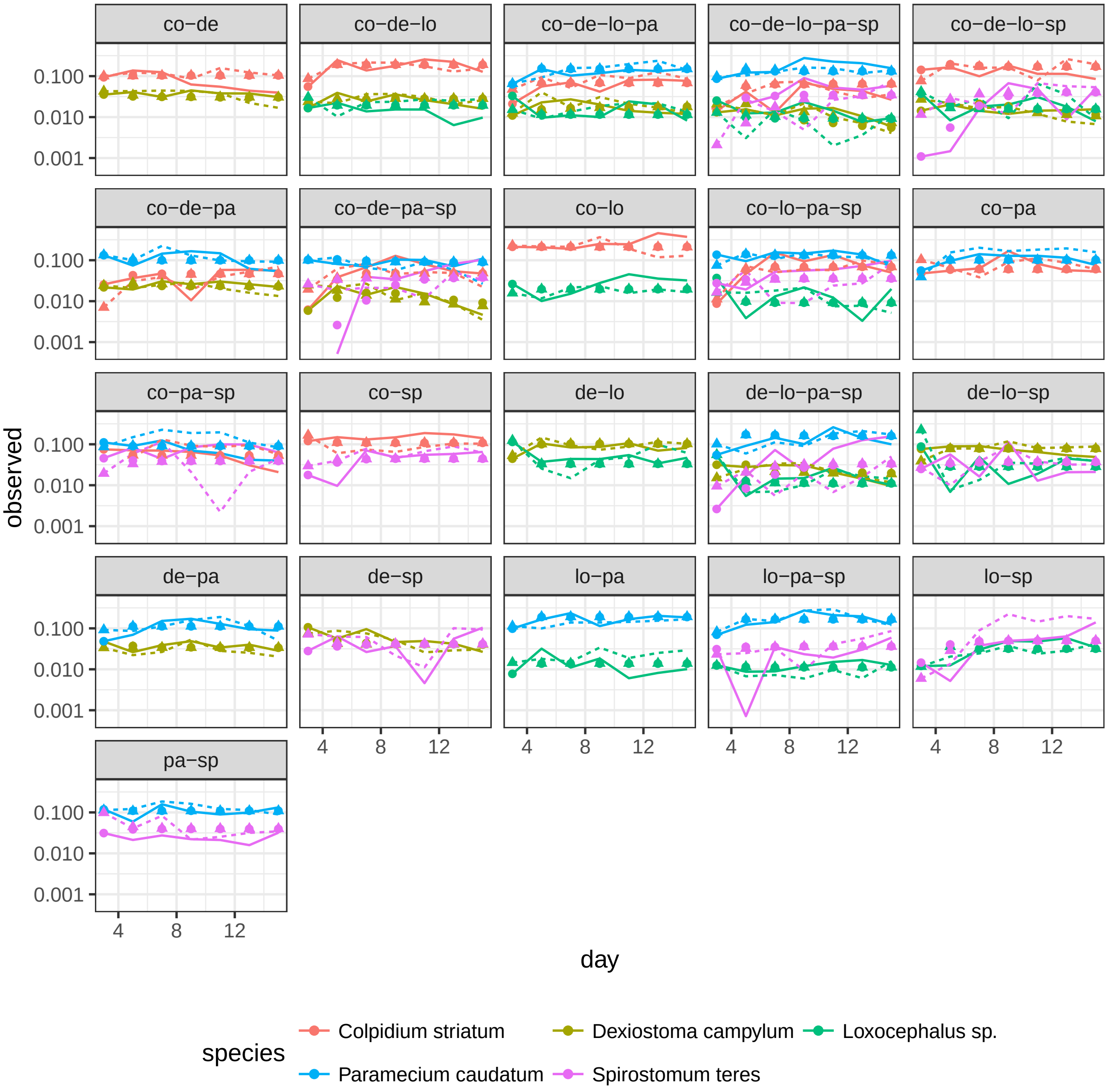

Supplementary Figure 13: The observed and predicted dynamics for the five-species protist system at 15$^o$C across days 1-17. The lines give the observed dynamics, with solid vs. dashed denoting each of two replicates. The symbols give the modeled values, with circles vs. triangles denoting the replicates. The best-fitting time series model essentially fits a smooth curve through the data, unable to capture the fine-scale temporal variation in species abundances.

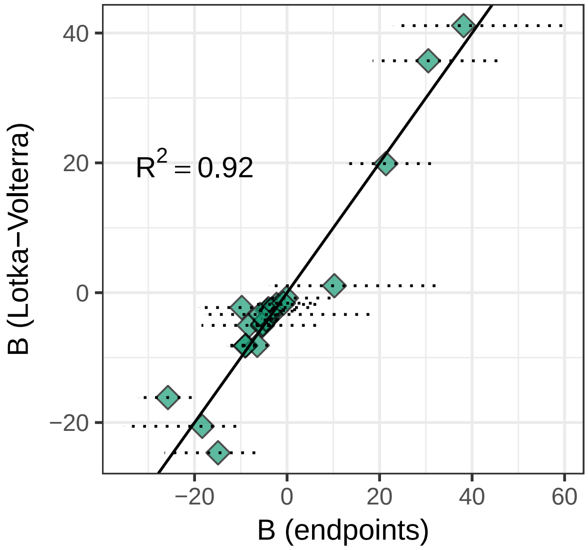

Supplementary Figure 14: Comparison of the $\tilde{A}$ matrix as obtained using only the endpoints (x-axis) versus trajectory matching of the time-series (y-axis). Dashed lines give the 95% prediction interval for the posterior. Both methods recover a similar structure.

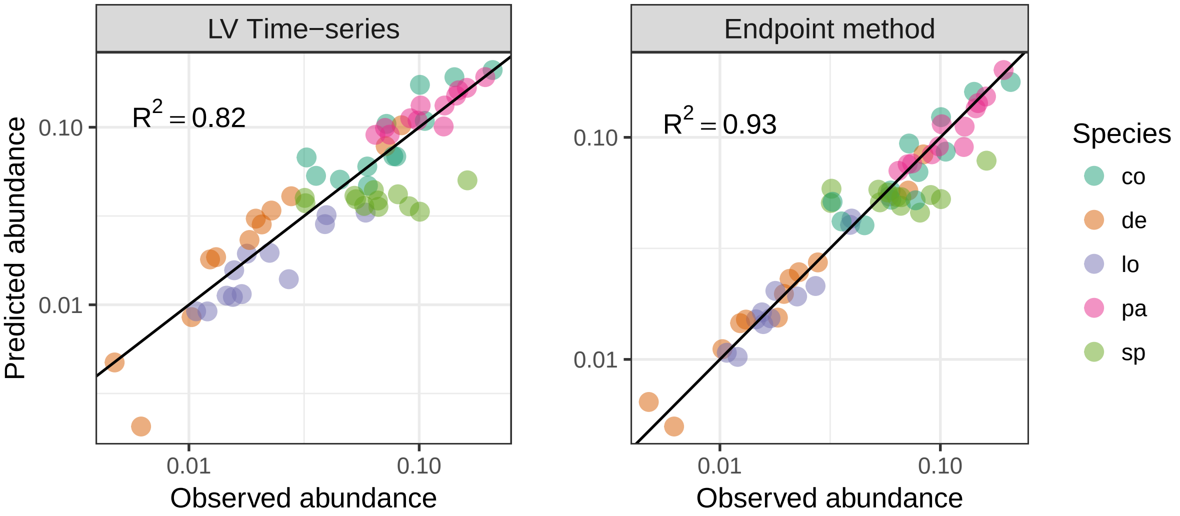

Supplementary Figure 15: The observed versus predicted abundances using the time-series (a) and the endpoints (b). The time-series method does a sufficient job predicting coexistence patterns, albeit with lower accuracy than the method proposed here. This lack of predictive power arises from the fact that the trajectory matching approach is trying to minimize the deviations between predicted and observed across the entire time series, not simply the final time point. The 1:1 line is shown in black.

#### C4 Effect of sampling during the transients

An important experimental consideration is when to sample the endpoints of the system. Because our approach assumes the observed endpoints are close to—but not exactly at—the centroid of the attractor, the goal of sampling should be to ensure that the dynamics have played out long enough to push the endpoints near to their true values. Thus, in practice, one should ensure that the dynamics have passed through the transient phase of rapid growth and turnover. In systems where the dynamics are oscillatory or chaotic, a single sampled endpoint might be quite far from the centroid of the attractor; however, provided the dynamics have played out long enough to ensure each species is caught in the basin of its attractor, then a sufficient number of replicates can be used to estimate the centroid. Note that this “centroid” may not be equivalent to the average of the endpoints, as clearly demonstrated in the non-equilibrium Lotka-Volterra simulation below, but rather the point about which the dynamics revolve.

On the one hand, these considerations suggest that experiments should be run for as long as possible to ensure the dynamics settle in to the attractor. Yet in many systems, progressive nutrient limitation or build-up of toxins may start to inhibit growth, leading to decline of the system. For example, as demonstrated in the fitting details for the Protist system (Fig. 2), the total community biomass stabilized between days 13 and 17 for all temperatures, suggesting that the community had reached a stationary distribution, but then declined.

To more rigorously demonstrate these effects, we can compare the quality of fit of our approach by sampling the the endpoints at different days (Fig. 16). Doing so reveals an initial period of transient dynamics between days 0 and 10 where the model fits relatively poorly, followed by a relatively stable period where the model exhibits high quality of fit ($R^2 \sim 0.95$), followed by a gradual reduction in fit as the dynamics deteriorate, perhaps due to resource depletion or toxin accumulation. Thus, provided one runs the experiment long enough to pass through the transients, our method can be relatively robust to the choice of when to sample (or, equivalently, the initial conditions). Nevertheless, in practice, effort should be taken to identify an optimal (or biologically relevant) sampling point on a system-by-system basis.

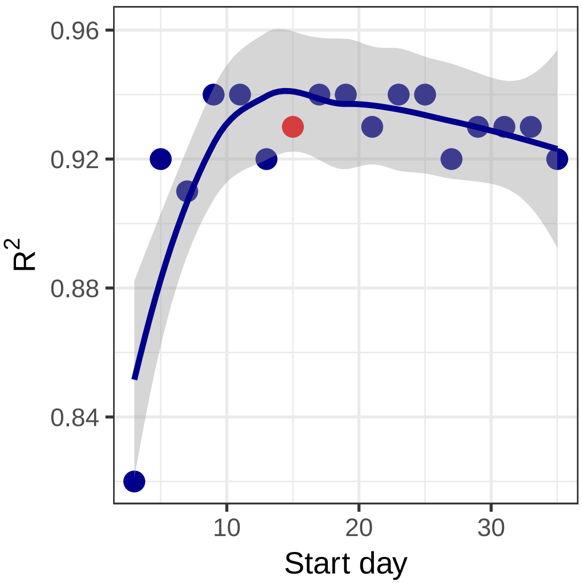

Supplementary Figure 16: The quality-of-fit for the protist system at 15$^o$C as a function of the sampling day. Rather than sample the community at days 15-17, as in the main text, we "ended" the experiment at the indicated day, ±2 days, and fit the model with the corresponding endpoints. These results demonstrate that there is a clear initial period where the model fits poorly due to transient dynamics; followed by a stable period between days 10 and 20 where the approach performs well; followed by a period where the quality of fit starts to deteriorate as the species decline in abundances. The point in red denotes the point used in the main analysis, independently identified by quantifying when total biomass of the community stabilized (Fig. 16).

### D Simulated data

In the previous sections, we have shown how our approach can be successfully applied to experimental data. In this section, we test the method's performance using data produced by numerical simulations, for which the model generating the data is known. We test the method using a variety of dynamical models, including cases of non-linear, out-of-equilibrium dynamics, and models containing "higher-order interactions" (i.e., where the relationships between species abundances are not pairwise).

For all models, we choose parameters at random, initialize the model with all combinations of species presence/absence, and then integrate dynamics for an arbitrarily long time. For each combination of species, we perform 96 simulations (as in a 96-well microplate) starting from different initial densities. For our method to be applicable, we need to observe a sufficiently large and diverse set of endpoints; consequently, we re-sample random parameters until we find a parameterization yielding a usable set of endpoints.

For each model and parameter set, we "record" the endpoints and attempt to recover the matrix $B$ by minimizing the sum of squares between the observed and predicted endpoints.predicted endpoints (i.e., using the sum-of-squares, rather than the Bayesian, fitting approach). The method's performance in recovering observed endpoints is therefore to be considered "in-fit". However, because our method is completely blind to initial conditions, cases in which species do not coexist (and therefore the system collapses to a different endpoint containing fewer species), are by definition "out-of-fit". Contrary to what was done when analyzing empirical data, we can therefore use simulations to assess whether our method can correctly predict cases in which the coexistence between a given set of species is precluded.

#### D1 Generalized Lotka-Volterra competition

To illustrate how simulations are performed, and introduce the plots used to assess the quality of the prediction, we start by considering the simplest case of Lotka-Volterra dynamics (Eq. 6), for which we should recover a perfect fit. We consider the parameters:

$$r = \begin{pmatrix} 0.58 \\ 0.93 \\ 0.72 \\ 0.7 \end{pmatrix},$$

$$A = \begin{pmatrix} -1.02 & -0.24 & -0.7 & -0.36 \\ -0.86 & -1.04 & -0.37 & -0.87 \\ -0.98 & -0.62 & -1.08 & -0.09 \\ -0.99 & -0.74 & -0.13 & -0.72 \end{pmatrix},$$

and integrate the dynamics for 1,500 time units when starting from a variety of initial conditions. Specifically, for each of the $2^4 - 1 = 15$ possible combinations of species presence/absence, we choose 96 random starting points, setting each species that is present at a density sampled independently from the uniform distribution $\mathcal{U}[0, 1]$.

For these parameters, we find that all species grow in isolation, and coexist when co-cultured in pairs—in all cases converging to a globally-attractive equilibrium (Fig. 17). Assemblages initialized with more than two species result in the extinction of one or more species, and are therefore never observed.

We use the endpoints for monocultures and pairs to parameterize the matrix $B$, and then use $B$ to predict the outcome of all other experiments (Fig. 17). The figure shows that, as expected, the method fits the observed data (final densities in monoculture or pairs) perfectly. Moreover, it correctly predicts the lack of coexistence in the experiments involving three species. The method also predicts a feasible equilibrium with all the species. However, the equilibrium is predicted to be unstable (under the assumption of equal growth rates), and is therefore not accessible "experimentally".

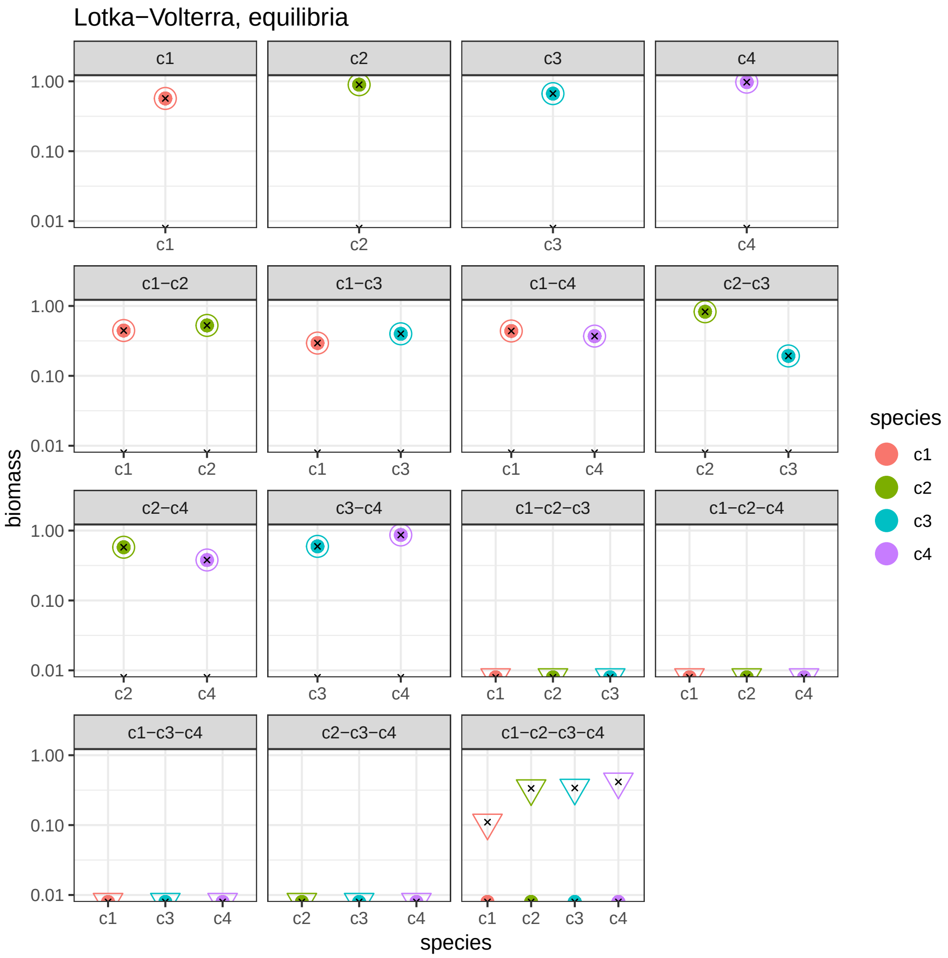

Supplementary Figure 17: Simulation results for a GLV competitive system. For each combination of the four competitors (*c1* to *c4*, colors), we ran 96 simulations starting from different initial conditions. Each panel shows the location of the simulations' endpoints (solid circles), as well as the true location of the equilibria for the system (crosses, computed analytically). Experiments resulting in a lack of coexistence are represented as half-points at the bottom of the graph. The predictions obtained using our method are reported using open symbols. There are two cases: we use circles for predictions of (locally) stable endpoints, and triangles for unstable ones; the stability is calculated under the assumption of Lotka-Volterra dynamics and equal growth rates. For this system, as expected, we recover a perfect fit for the positive densities. We can also correctly predict the lack of coexistence among triplets. We predict perfectly the position of the four-species equilibrium, and correctly classify it as unstable, despite having used growth rates that differ substantially from each other.

#### D2 Generalized Lotka-Volterra with limit cycles

With enough species, the Generalized Lotka-Volterra system can produce wide range of dynamics. To test how the method performs for the case of out-of-equilibrium dynamics, we kept sampling parameters at random until we found a parameterization that leads to widespread coexistence, but exhibits a limit cycle (for experiments in which all the species are present). Specifically, we use the parameters:

$$r = \begin{pmatrix} 1.56 \\ 1.69 \\ 2.11 \\ 0.61 \end{pmatrix},$$

$$A = \begin{pmatrix} -0.65 & -0.75 & -0.97 & -0.1 \\ -0.44 & -1.22 & -0.1 & -0.9 \\ -0.89 & -0.36 & -0.99 & -0.83 \\ -0.3 & -0.01 & -0.27 & -0.27 \end{pmatrix},$$

and integrate the dynamics for a variety of initial conditions, as done for the previous case.

The results (Fig. 18) are more interesting than in the case of equilibrium dynamics examined above, and include a case of dependence on initial conditions (*c1-c3-c4*, signaling that the equilibrium is locally, but not globally, stable), and a limit cycle. The proposed method fares well, correctly predicting the lack of coexistence for four combinations, and finding the right location of the equilibrium surrounded by the limit cycle. The four-species equilibrium is however misclassified as stable (because our estimates assumes equal growth rates, while they are substantially different for the parameterization above).

#### D3 Competition with Allee effect

Having shown that the method can correctly predict instances in which species can or cannot coexist when dynamics are defined by the GLV model, we turn to more complex models. To

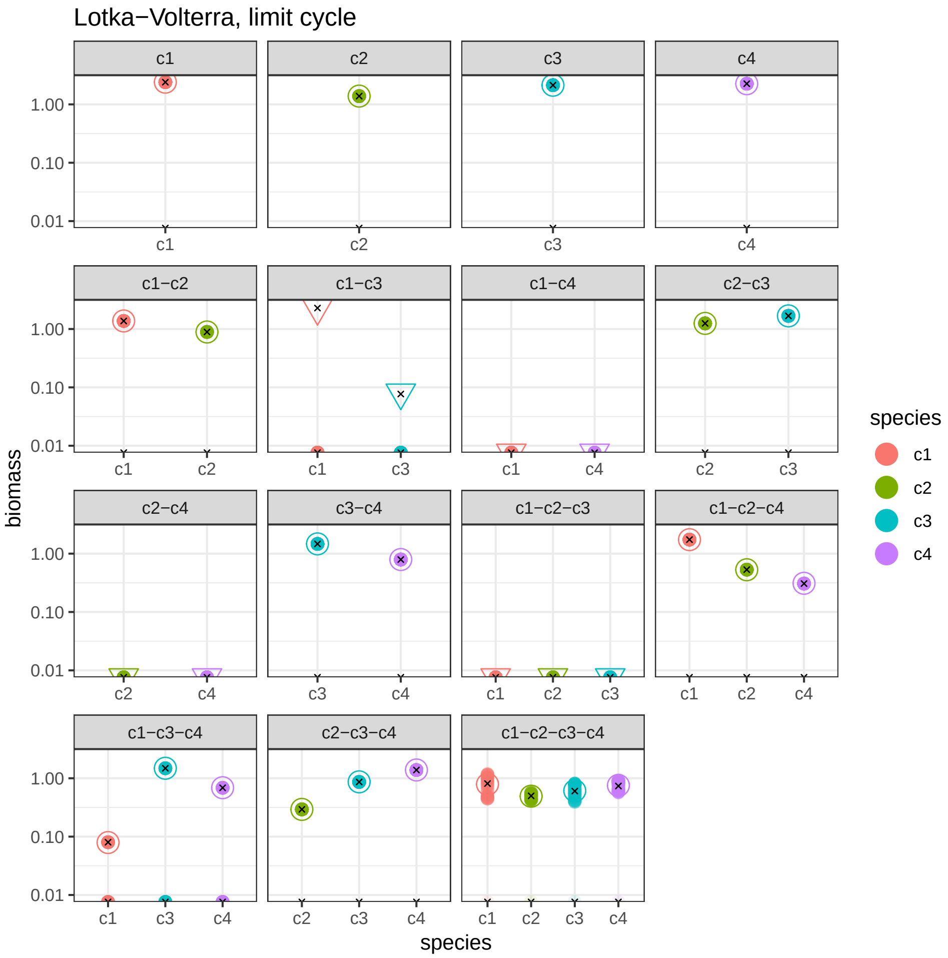

Supplementary Figure 18: Simulation results for a GLV competitive system exhibiting a limit cycle when all competitors are included. Colors and symbols are as in Fig. 17. Notable features of this system are: competitors might (*c1*-*c2*, *c2*-*c3*) or might not (*c1*-*c3*, *c1*-*c4*) coexist in pairs. However, in one case a feasible but unstable equilibrium exists (*c1*-*c3*, correctly predicted by our method), while in the other there is no feasible equilibrium (*c1*-*c4*, also correctly predicted). The system including *c1*-*c3*-*c4* shows dependence on initial conditions (some trajectories collapse to another system, while others converge to equilibrium), signaling a locally (but not globally) stable equilibrium. The method correctly identifies the position of the 4-species equilibrium surrounded by the limit cycle. However, it suggests stability for the equilibrium, while it must be unstable to give rise to the stable limit cycle. The misclassification stems from the fact that in the calculation of stability, we consider growth rates to be equal (because we cannot infer growth rates from endpoints), while this is not the case here.

start, we consider a model in which competitors experience an Allee effect[10]:

$$\frac{\mathrm{d}x_i}{\mathrm{d}t} = r_i x_i \left(\frac{x_i}{s_i} - 1\right)\left(1 - \frac{x_i}{k_i}\right) + r_i x_i \sum_j A_{ij} x_j \ ,$$

in which species that are at low abundance ($x_i(t) < s_i$) cannot grow and will go extinct. This model gives rise to several (three for a single species) equilibria, and displays strong multistability. We choose the parameters:

$$r = \begin{pmatrix} 0.1 \\ 0.38 \\ 0.06 \\ 0.96 \end{pmatrix},$$

$$A = \begin{pmatrix} 0 & -0.92 & -0.1 & -0.69 \\ -0.49 & 0 & -0.62 & -0.32 \\ -0.7 & -0.38 & 0 & -0.97 \\ -0.4 & -0.39 & -0.07 & 0 \end{pmatrix},$$

$$k = \begin{pmatrix} 0.02 \\ 0.1 \\ 0.46 \\ 0.05 \end{pmatrix},$$

$$s = \begin{pmatrix} 0.01 \\ 0.04 \\ 0.16 \\ 0.02 \end{pmatrix},$$

and integrate the dynamics initializing the species at random (uniformly chosen initial densities between 0 and $2\max(k_i)$). Despite the fact that this model contains cubic terms, the fit to the observed data is very good (Fig. 19). Predictions of the location of the positive endpoints is perfect, while we predict the stable coexistence for all other configurations—which however always result in extinctions. This shows that when the model is misspecified, the results can be less reliable.

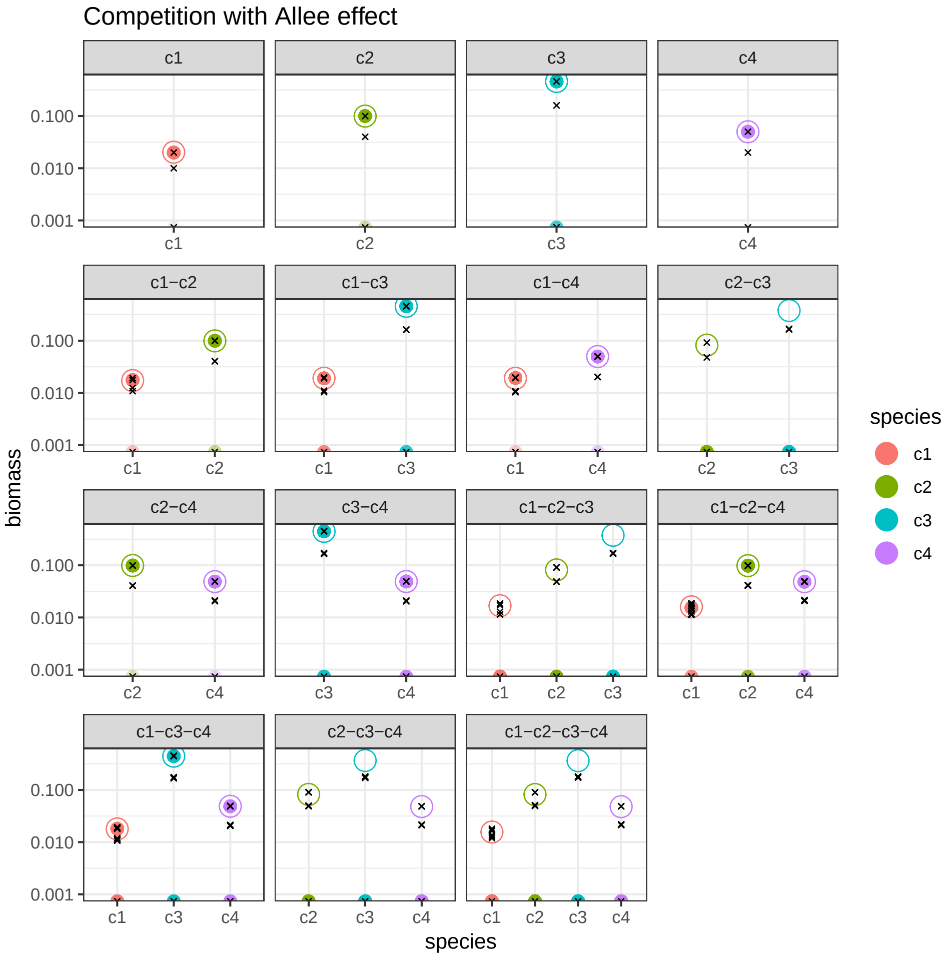

Supplementary Figure 19: Simulation results for a competitive system in which competitors cannot grow when rare. This results in multistability in all cases (half-points at the bottom of each graph signal trajectories that resulted in extinctions). Despite the fact that the model contains cubic terms (while our method can deal only with quadratic terms), the in-fit is excellent, in all cases fitting the location of the endpoints perfectly. Experiments in which the species do not coexist are however misclassified—while there exist equilibria close to the prediction, they are unstable, rather than stable as predicted by our method.

#### D4 Facultative mutualism with saturation.

To test the method under stronger departures from GLV, we adapt the model for facultative mutualism by Holland *et al.*[11], and include competition between members of the same group:

$$\frac{\mathrm{d}x_i}{\mathrm{d}t} = x_i \left( r_i - \frac{x_i}{k_i} + \sum_j \frac{A_{ij}x_j}{1 + \sum_k h_i x_k A_{ij}} + \sum_j B_{ij}x_j \right) \ ,$$

in which two set of species (plants, animals) can grow in isolation, but benefit from the interaction with each other. We choose the parameters:

$$r = \begin{pmatrix} 1.71 \\ 1.25 \\ 1.39 \\ 1.09 \end{pmatrix},$$

$$h = \begin{pmatrix} 0.96 \\ 0.01 \\ 0.57 \\ 0.76 \end{pmatrix},$$

$$A = \begin{pmatrix} 0 & 0 & 0.54 & 0.21 \\ 0 & 0 & 0.88 & 0.82 \\ 0.99 & 0.88 & 0 & 0 \\ 1.32 & 0.54 & 0 & 0 \end{pmatrix},$$

$$B = \begin{pmatrix} -0.64 & -0.03 & 0 & 0 \\ -0.62 & -1.01 & 0 & 0 \\ 0 & 0 & -0.88 & -0.42 \\ 0 & 0 & -0.63 & -0.96 \end{pmatrix},$$

and again integrate dynamics for an arbitrary long time starting from a variety of initial conditions. Despite the fact that the model contains a saturating functional response, the method recovers almost perfect predictions for all cases, and correctly predicts the lack of coexistence between the two plants (Fig. 20).

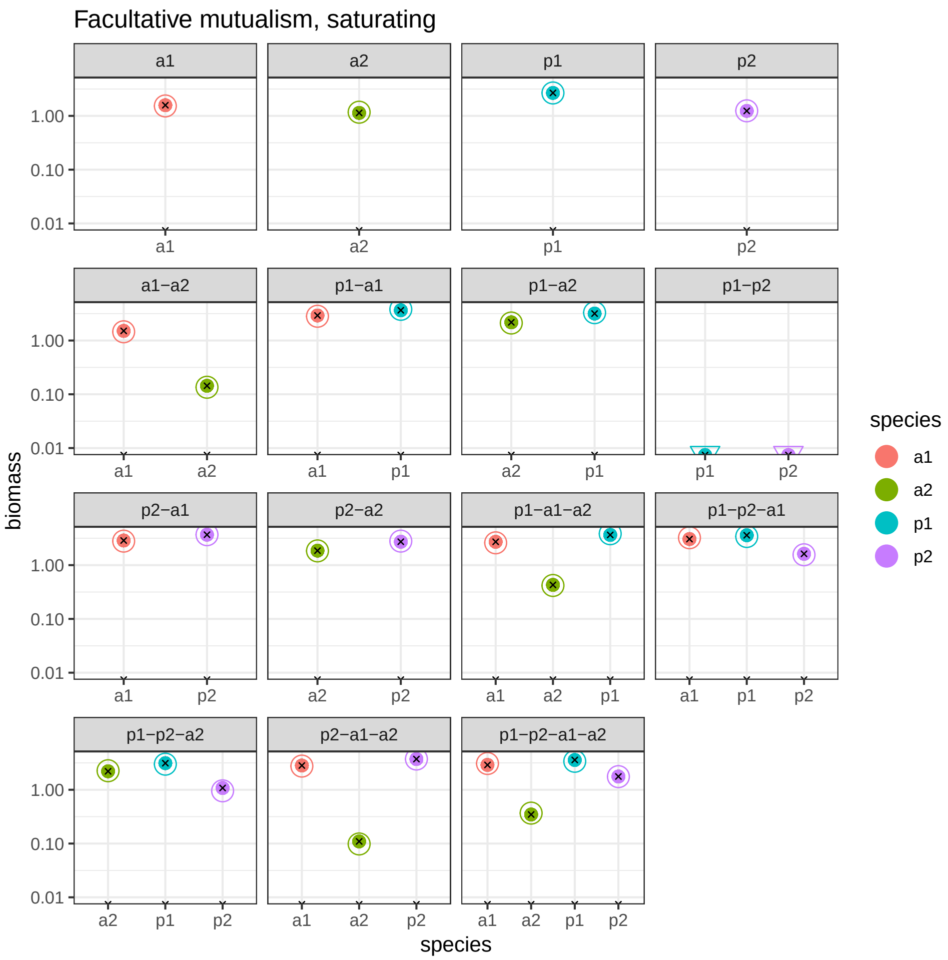

Supplementary Figure 20: Simulation results for a system of facultative mutualism between two classes of competitors (plants, $p$, and animals, $a$). Despite the non-linear functional response, the method predicts the location of the endpoints (all characterized by equilibrium dynamics) almost perfectly. The method also correctly predicts the lack of coexistence between the two plants (panel *p1-p2*).

#### D5 Consumption with saturation

We then adapted a consumer-resource model parameterized as in Fussman & Heber[12]:

$$\frac{\mathrm{d}x_i}{\mathrm{d}t} = r_i x_i - s_i x_i^2 + x_i \left( \sum_j \frac{A_{ij} x_j}{1 + \sum_k B_{kj} x_k} - \frac{\sum_j A_{ji} x_j}{1 + \sum_k B_{ki} x_k} \right) ,$$

with parameters:

$$r = \begin{pmatrix} 0.82 \\ 0.93 \\ -0.46 \\ -0.41 \end{pmatrix},$$

$$s = \begin{pmatrix} 0.1 \\ 0.1 \\ 0.1 \\ 0.1 \end{pmatrix},$$

$$A = \begin{pmatrix} 0 & 0 & 0 & 0 \\ 0 & 0 & 0 & 0 \\ 0.8 & 0.87 & 0 & 0 \\ 0.31 & 0.82 & 0 & 0 \end{pmatrix},$$

$$B = \begin{pmatrix} 0 & 0 & 0 & 0 \\ 0 & 0 & 0 & 0 \\ 0.48 & 0.62 & 0 & 0 \\ 0.19 & 0.36 & 0 & 0 \end{pmatrix},$$

defining a system with two resources (*r1*, *r2*) and two consumers (*c1*, *c2*). The dynamics for these simulations are rich, encompassing four limit cycles, and a five cases of equilibrium dynamics (Fig. 21). Remarkably, our method predicts the location of the equilibria perfectly, and correctly predicts all cases of coexistence and lack of coexistence. The only incorrect inference is that it predicts the equilibria giving rise to the limit cycles to be stable.

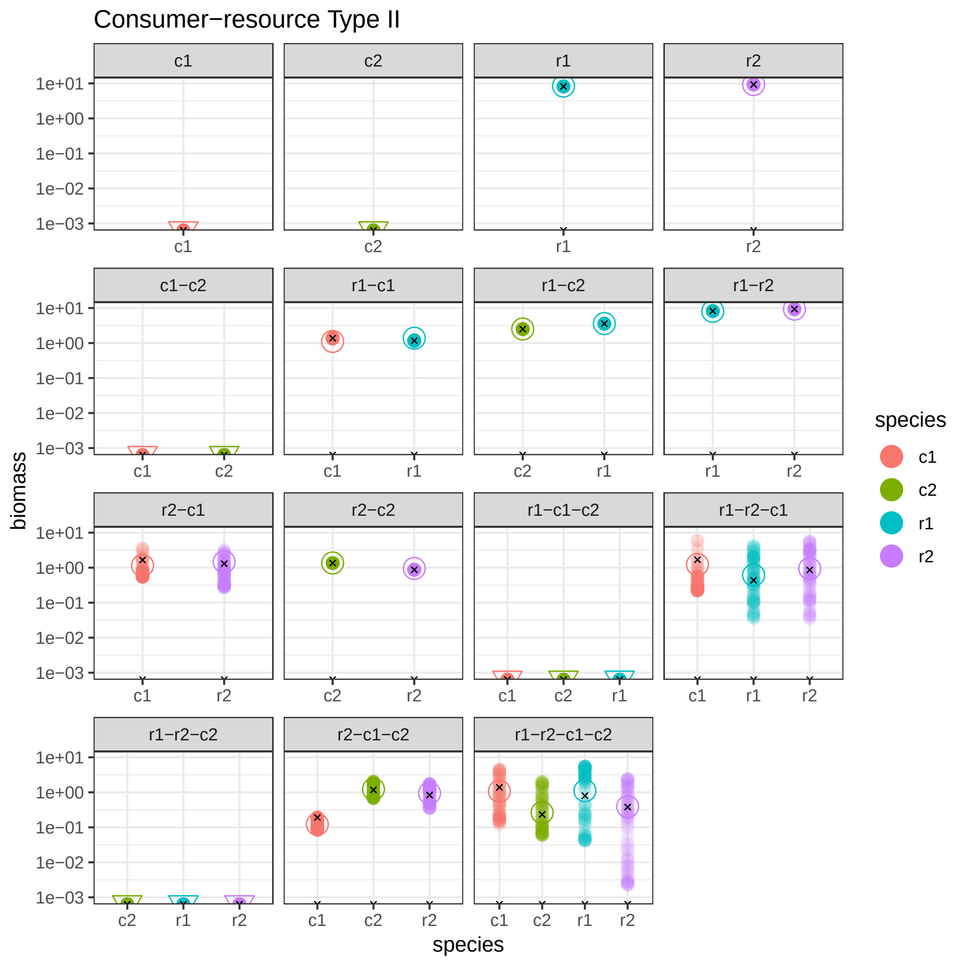

Supplementary Figure 21: Simulation results for a consumer-resource system. Two resources and two consumers are simulated in all possible combinations, giving rise to cases of coexistence at equilibrium, stable limit cycles, or extinctions. In all cases, the proposed method predicts the location of the equilibria of the nonlinear system quite perfectly, making the correct inference for all cases in which species cannot coexist.

#### D6 Higher-order interactions

Finally, we consider a case in which interactions are not fundamentally pairwise. We extend the GLV model to higher-order interactions by considering the system of equations:

$$\frac{\mathrm{d}x_i}{\mathrm{d}t} = x_i \left( r_i + \sum_j A_{ij} x_j + \sum_{j,k} B_{ijk} x_j x_k \right) ,$$

where $B$ is a three-dimensional tensor. We want to focus on higher-order interactions that can be interpreted as a third species modifying the interaction between other two species. To make this more explicit, we rewrite the equations as:

$$\frac{\mathrm{d}x_i}{\mathrm{d}t} = x_i \left( r_i + \sum_j \left( A_{ij} + \sum_k B_{ijk} x_k \right) x_j \right) .$$

We set $B_{ijk} \neq 0$ only if $i \neq j \neq k$ (the other terms would give rise to quadratic or cubic effects of $i$ and $j$ on the growth of $i$). Note also that only the symmetric part $B_{ijk} + B_{ikj}$ matters for the dynamics, and therefore we can consider $B_{ijk} = B_{ikj}$ without loss of generality. With these simplifications in place, we choose the parameters:

$$r = \begin{pmatrix} 0.31 \\ 0.41 \\ 0.43 \\ 0.19 \end{pmatrix},$$

$$A = \begin{pmatrix} -0.5 & -0.11 & -0.28 & -0.78 \\ -0.65 & -0.83 & -0.78 & -0.27 \\ -0.98 & -0.42 & -1 & -0.07 \\ -0.03 & -0.21 & 0 & -0.93 \end{pmatrix},$$

$$B_1 = \begin{pmatrix} 0 & 0 & 0 & 0 \\ 0 & 0 & -0.16 & 0.19 \\ 0 & -0.16 & 0 & 0.58 \\ 0 & 0.19 & 0.58 & 0 \end{pmatrix},$$

$$B_2 = \begin{pmatrix} 0 & 0 & 1.52 & 2.51 \\ 0 & 0 & 0 & 0 \\ 1.52 & 0 & 0 & -0.32 \\ 2.51 & 0 & -0.32 & 0 \end{pmatrix},$$

$$B_3 = \begin{pmatrix} 0 & -2.03 & 0 & -1.53 \\ -2.03 & 0 & 0 & 1.47 \\ 0 & 0 & 0 & 0 \\ -1.53 & 1.47 & 0 & 0 \end{pmatrix},$$

$$B_4 = \begin{pmatrix} 0 & 1.92 & 1.37 & 0 \\ 1.92 & 0 & 0.01 & 0 \\ 1.37 & 0.01 & 0 & 0 \\ 0 & 0 & 0 & 0 \end{pmatrix},$$

where we have written each slice of the tensor separately.

Despite the strong higher-order interactions, we can find a solution that approximates all coexistence endpoints quite closely (Fig. 22). Our method also predicts correctly two cases in which the species cannot coexist. For the remaining two triplets, however, the method predicts stable coexistence, while coexistence was not observed in the simulations. This shows that the presence of sufficiently strong higher-order interactions would result in a very poor quality of predictions, requiring the method to be extended as detailed in the main text.

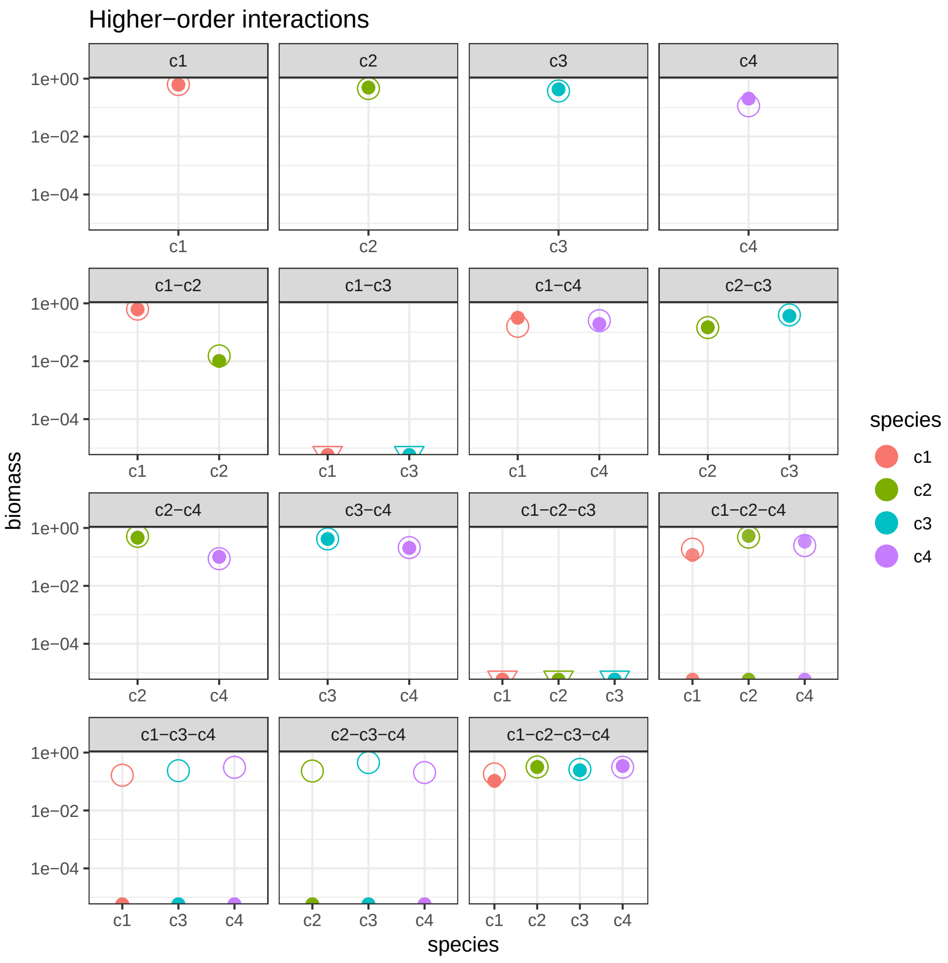

Supplementary Figure 22: Simulation results for a system characterized by higher-order interactions. Despite the strong effect of HOIs, the recovered solution is close to all endpoints. Moreover, the method correctly predicts that coexistence between *c1* and *c2*, or *c1*, *c2* and *c3* is precluded. The method however predicts coexistence between two triplets, despite simulations showing that either no feasible equilibrium exists, or it is unstable.

### E Experimental design

#### E1 Number of experiments

As highlighted in the Methods section, endpoints in which multiple species coexist appear in multiple matrices $E_i$. In particular, if an endpoint contains $l$ species, it will be reported in $l$ distinct matrices $E_i$. Each $E_i$ is used to fit one row of $B$, and each row of $E_i$ places one additional constraint on the coefficients of $B_i$. Clearly, then, speciose endpoints provide many constraints at once, effectively yielding more information about $B$ compared to experiments in which few species coexist. This means that by observing species-rich endpoints, one might fit the coefficients of $B$ very efficiently, using a small number of experiments. Here we show that this observation suggests efficient strategies for experimental design that are fundamentally different from traditional approaches based on growing species in monocultures and pairs.

As an illustration, take the idealized case of a pool of $n$ species in which every possible assemblage of species yields coexistence at a unique endpoint, and for which we can measure endpoint abundances without error. To infer $B$, we need to fit $n^2$ coefficients, and therefore we need to write $n^2$ linearly independent equations, subject to the conditions outlined in Methods. As one necessary condition, each species must occur in $n$ distinct endpoints (equations). Using the endpoint in which all $n$ species coexist, we can thus write $n$ equations, and using each of the $n$ endpoints in which all but one species are present, we can write $n - 1$ equations ($n^2 - n$ in total). As such, by using only $n + 1$ endpoints (complete community + leave-on-out communities), we can parameterize the matrix $B$.

This stands in contrast to more typical designs, in which all species are grown in isolation and in pairs. With this experimental design, even assuming all experiments lead to coexistence, we would need to conduct $n$ single-species experiments and $\binom{n}{2}$ experiments with pairs of species, for a total of $n(n+1)/2$ experiments. This number grows proportionally to $n^2$, while the number of experiments required for the "top-down" approach introduced above grows linearly in $n$. For $n = 10$, estimating $B$ using the top-down approach would require $11$ experiments, while the "bottom-up" approach uses $55$. For $n = 20$, combining single species and pairs would require 10 times as many experiments as the using the full pool and all leave-one-out communities (210 experiments instead of 21).

Of course, real ecological systems do not satisfy these idealized conditions: coexistence is usually not feasible for all species assemblages, and measurement errors can be minimized but not avoided. The former deviation is more challenging to account for. It is difficult to know *a priori* whether coexistence in experimentally manipulated assemblages will be common or rare, and this will certainly depend on the nature of interactions in any particular system. Additionally, if the likelihood of coexistence varies systematically with community size, as expected under most ecological theories, this will have significant bearing on the choice of experimental design. However, we note that top-down designs should be generally more robust to lack of coexistence than bottom-up designs. By this, we mean that in the bottom-up design, whenever two species do not coexist in a pairwise community, the system will necessarily collapse to a monoculture endpoint, or to no species at all. In either case, this results in the loss of a distinct endpoint, adding no new information for inference beyond replication of single-species endpoints. On the other hand, using a top-down design, if all species do not coexist in the full or leave-one-out assemblages, the system might collapse to any one of a large number of sub-communities, potentially still containing a significant fraction of the species. Although such an outcome generates fewer constraints than desired, it is still likely to contribute new information. For this reason, large initial assemblages are unlikely to result in wasted experiments.

#### E2 Simulations results

We can investigate the effect of measurement error more directly through simulations. To probe which experimental designs are most efficient in the presence of noise, we constructed a 6-species GLV competitive system in which all communities form feasible, stable equilibria, and computed all of the $2^6 - 1 = 63$ possible endpoints. We then created five noisy "replicate" measurements for each endpoint by perturbing the true abundance of each species in the endpoint by up to $15\%$: $x_i^{(k)} \sim z_i^{(k)} \mathcal{U}[0.85, 1.15]$ (similar results are obtained for any moderate level of error). Because all species coexist for any assemblage, we have many possible choices for experimental designs in which we estimate the matrix $B$ using a fraction of the data, and predict the rest of the experiments out of fit. We therefore explored several realistic experimental designs: (i) *leave-one-outs* + *all* (i.e., we infer $B$ using only the endpoints containing five or six species—7 assemblages in total, and predict the abundance of the species in all the other 56 assemblages); (ii) *mono* + *leave-one-outs* (i.e., using only endpoints containing either one or six species—12 assemblages in total); (iii) *mono* + *leave-one-outs* + *all* (13 assemblages in total); (iv) *mono* + *pairs* (21 assemblages); (v) all *quadruplets* (15 assemblages); (vi) all *triplets* (20 assemblages); (vii) *mono* + *pairs* + *leave-one-outs* + *all* (28 assemblages); (viii) *mono* + *pairs* + *triplets* (41 assemblages).

For each of these designs, whenever feasible (e.g., there is only one way to parameterize $B$ using 7 assemblages—the design (i) above) we also generated 100 randomized designs, using the same number of assemblages chosen at random. Of course, while the fixed designs were chosen to guarantee our ability to fit $B$ (provided that all subsets coexist), the random designs were not. For example, a random selection of endpoints might not have every species represented $n$ times. This problem is severe when trying to devise a random experimental design based on a small numbers of endpoints. We therefore discarded designs that did not allow for the fit of $B$ and kept sampling until we had 100 random configurations that could be used to infer the whole matrix. In some cases, these 100 random designs included repetitions. However, for all cases but the extremes, repetitions were rare.

For each of the 8 designs introduced above, and for the random designs, we inferred the corresponding $B$ from the noisy endpoints, and then predicted the abundance of the species in all the remaining endpoints that were not used to fit the data. We scored the quality of the prediction by computing the mean squared log-deviation: $\mathbb{E}\left[\left(\log\left(x_i^{(k)}\right) - \log\left(\hat{z}_i^{(k)}\right)\right)^2\right]$, where $\hat{z}_i^{(k)}$ is the predicted, and $x_i^{(k)}$ is the observed abundance of species $i$ in endpoint $k$. Note that this metric penalizes equally errors made when estimating low- and high-abundance species, and that the use of the logarithm is justified by the way the noise was introduced (i.e., proportionally to abundances). Because we are predicting out-of-fit data, one can recover predictions with negative abundances. In such cases, the estimate is not only quantitatively erroneous (as is inevitable when the system is noisy), but rather qualitatively wrong: the method would predict that certain assemblages do not coexist, when in fact they do.

The results are presented in Figure 23. When densities are measured with error, the minimal design using only the *leave-one-outs + all* (top-down) performs poorly. Adding the endpoints for the monocultures, however, improves the performance considerably—notably, this design performs much better than a design taking 12 assemblages at random (many of which return qualitatively wrong predictions). What is perhaps most striking, however, is that this design outperforms almost *all* other designs, some of which require more than three times the amount of data. Also notable is the fact that the traditional design of *mono + pairs* fares quite badly (despite requiring many experiments). In fact, it is among the worst possible designs requiring 21 assemblages, and performs worse than many random designs using half as many points to fit the data. Similarly, the design utilizing all *triplets* is among the worst that can be found using 20 assemblages, and the bottom-up design *mono + pairs + triplets* is among the worst designs requiring 41 assemblages to predict the remaining 22 out-of-fit.

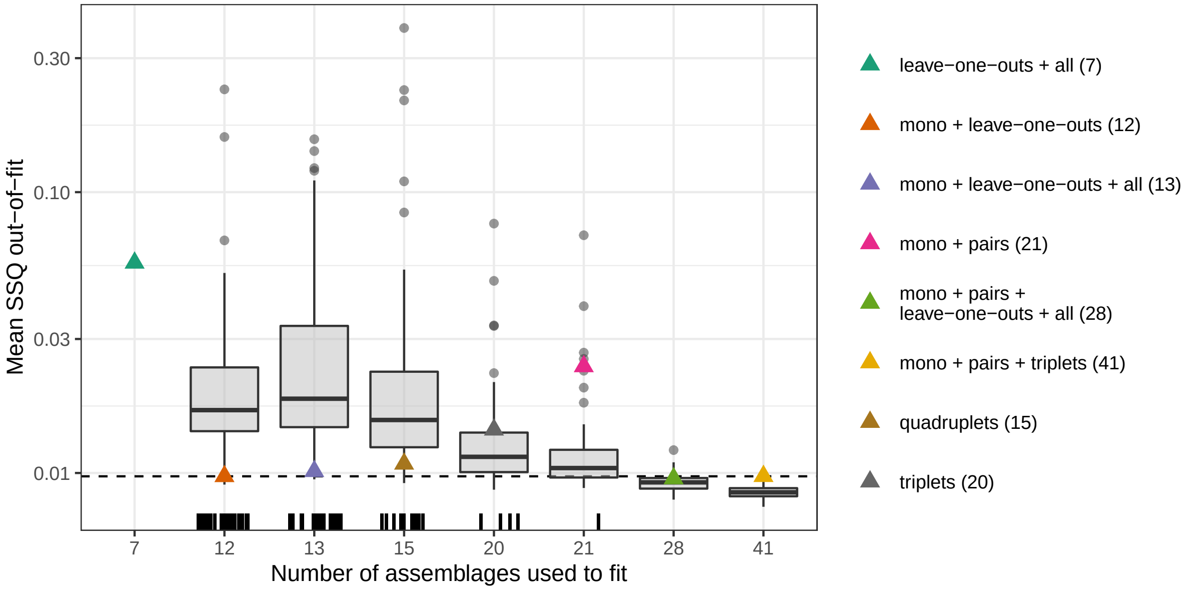

Supplementary Figure 23: Quality of fit for different experimental designs. We simulated a 6-species GLV model, in which all 63 possible assemblages lead to coexistence. We measured abundances at these endpoints and added noise, producing five "replicates". For each design, we use the specified number of assemblages to fit the model, and predict out-of-fit the abundance of all species at all other endpoints. Designs that produce qualitatively wrong predictions (i.e., predicting a lack of coexistence for assemblages that do in fact coexist) are represented by vertical bars at the bottom of each boxplot. The horizontal dashed line marks the performance of the *mono + leave-one-outs* design, which fares among the best designs overall, despite using only 12 assemblages to predict the remaining 51.

These results suggest that using a combination of experiments with few species and experiments with many species (e.g., *mono* + *leave-one-outs*) gives the best trade-off between quality of fit and number of required experiments. This makes sense when the inference of $B$ is viewed as fitting $n$ hyperplanes to the endpoint data. Exactly as in linear regression, the best strategy is then to choose endpoints that are somewhat distant from each other, "anchoring" the hyperplane more securely. Viewed another way, using a diversity of endpoint sizes minimizes the extent to which any prediction requires extrapolation beyond the range of the data used to fit the model. In a competitive system, a species $i$ is found at highest abundances when growing in isolation, and lowest when grown with many other species; therefore, it is not surprising that a combination of bottom-up and top-down works best.

#### E3 Testing experimental designs on real data

In the previous sections, we have highlighted that a) it is possible to fit the matrix $B$ (and therefore predict coexistence for any assemblage) using few (on the order of $n$) experimental endpoints, and b) experimental designs containing a mix of highly speciose and less diverse endpoints should perform better than traditional approaches in which all the monocultures and pairs of species are considered. In this section, we put this notion to the test using the data by Kuebbing *et al.*[6].

For both the native species and the invasive species experimental systems, we have access to measurements for 14 out of the 15 possible assemblages that can arise from a pool of four species, with each assemblage replicated ten times, and all species coexisting for each assemblage. Here we consider as an "experimental design" a subset of the 14 experimental assemblages: as for the simulations in Section E2, we fit matrix $B$ using a given number of distinct assemblages, and then we attempt to predict out of fit all other endpoints.

For these systems, one can show that there are 3,282 possible experimental designs that contain a sufficient diversity of endpoints to estimate the matrix $B$ (i.e., satisfying the conditions outlined in the Methods); of these, 15 designs require measuring the endpoints for only 6 assemblages, 215 designs are based on 7 assemblages, and so on, with a single, complete experimental design making use of all 14 assemblages. We consider here all designs making use of 13 or fewer assemblages, and predict out-of-fit the abundances of all the species in the remaining endpoints, as done in the previous section for simulated data. Of the specific designs considered before, only *mono* + *leave-one-outs* + *all* (using 8 assemblages to predict the remaining 6), *mono* + *pairs* (10 assemblages) and *mono* + *pairs* + *triplets* (using all assemblages but the 4-species community) are feasible (one of the leave-one-outs was not used in the experiments, precluding the possibility of fitting the model using only 5 assemblages).

The results are presented in Fig. 24 for the native community and in Fig. 25 for the non-native community. Both system yield very similar results: First, the errors are much larger than in the simulated data (i.e., the measurements are more noisy, or endpoints are not exactly described by a linear model)—this results in many designs failing to predict coexistence correctly (e.g., in the native system, 41 out of 215 designs using 7 assemblages fail to predict coexistence out-of-fit). Second, among the designs that return qualitatively wrong predictions we find the traditional design with monocultures and pairs. Third, the design including only monocultures, leave-one-outs and all species (i.e., predicting all pairs out of fit) performs almost as well as (non-native plants) or better than (natives) the design including all data but the full community (which in fact is among the worst designs using 13 assemblages)–again making the point that mixing high- and low-diversity assemblages provides a good starting point to predict the outcomes of experiments.

While in the main text Fig. 2 and in Section B we have presented results in which only one assemblage was excluded from the fit (each in turn), and predicted abundances with high accuracy, here we show that for well-replicated and well-behaved systems, as in the Kuebbing

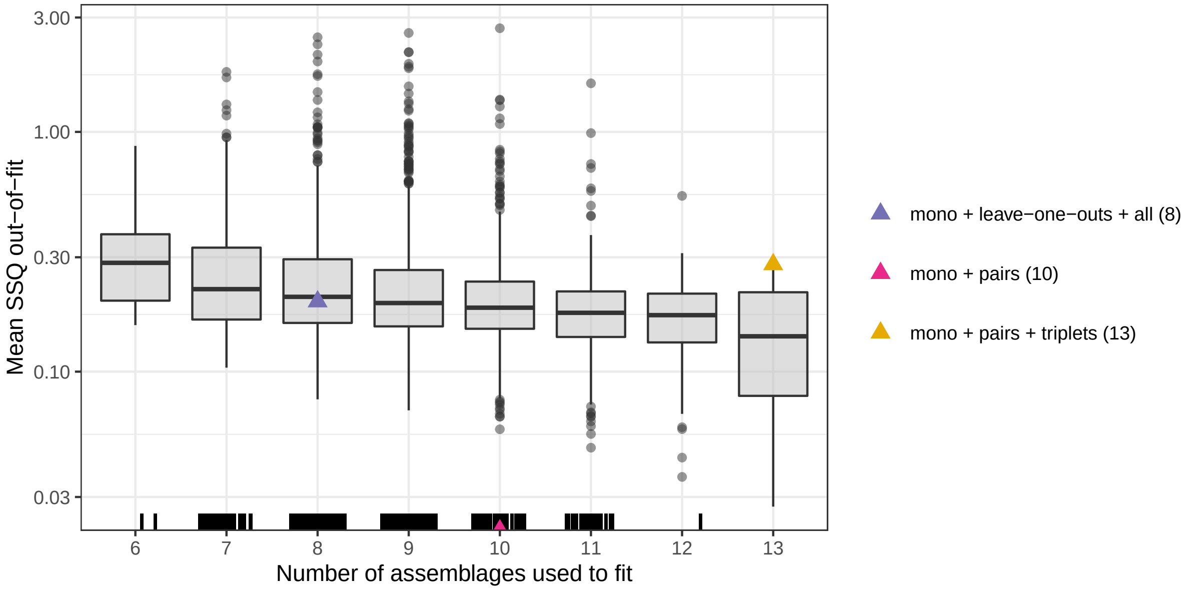

Supplementary Figure 24: Quality of fit for different experimental designs using the data by Kuebbing *et al.*[6] (native plants). For each of the 3,281 experimental designs (each requiring a certain number of assemblages, $x-$axis, and excluding the "complete" design, which leaves no endpoints out-of-fit), we can evaluate the goodness of out-of-fit predictions as in Fig. 23. Notably, the best-performing designs that are based on 7 or 8 endpoints (i.e., considering about half of the data) perform almost as well as designs using almost all endpoints. The traditional design of monocultures + pairs fails to predict coexistence for some of the out-of-fit assemblages, while a design mixing monocultures and leave-one-out assemblages produces good results.

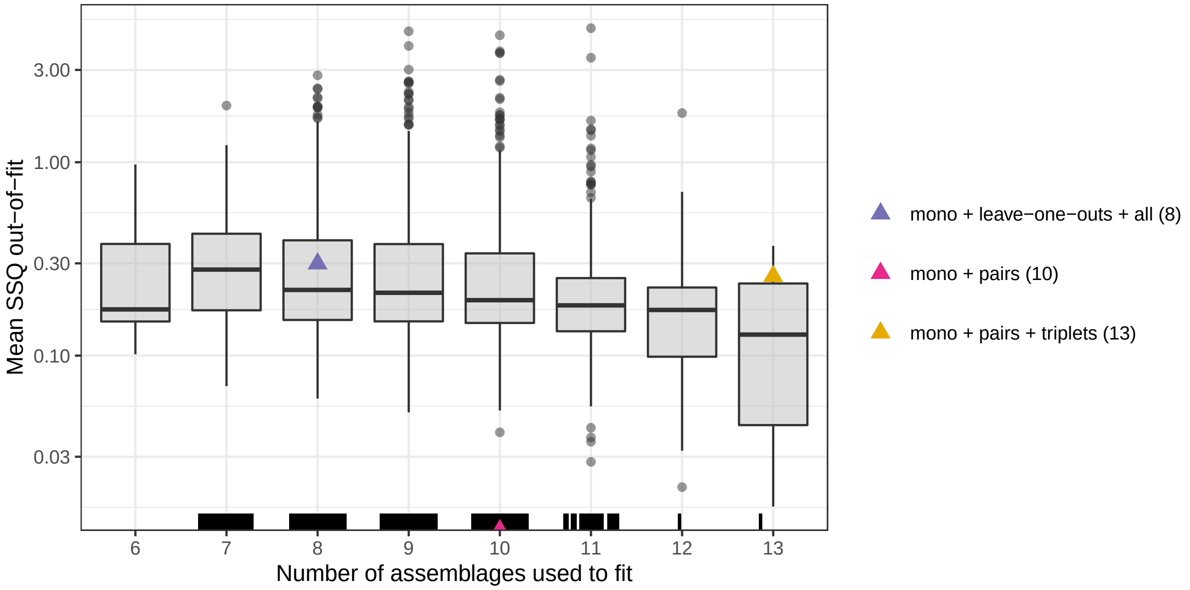

Supplementary Figure 25: As Fig. 24, but using the data for the non-native plants community.

*et al.*[6] study, we can make highly accurate predictions using a fraction of the available data. For example, in Fig. 5 of the main text we show the quality of fit of the best and worst 6-assemblage designs for the native plants. Fig. 26 shows the same analysis for the non-native plants system.

#### E4 Iterative designs

To conclude, we note the potential for iterative experimental design procedures, which might be useful to navigate large species pools when coexistence is not ubiquitous. For example, one could conduct an initial set of experiments using a fixed design (e.g., *singles* + *leave-one-outs* + *all*, totaling $2n+1$ experiments). Some experimental assemblages might experience extinctions, collapsing to smaller sub-systems, and precluding a full fit of $B$. If so, one could examine which pairs of species co-occur in this dataset, and which pairs have not yet been observed. Then, a second round of experiments can be designed to maximize the chance of observing new endpoints with previously unobserved co-occurence. For example, one might initialize experiments with combinations of species that have not been observed to co-occur, or choose

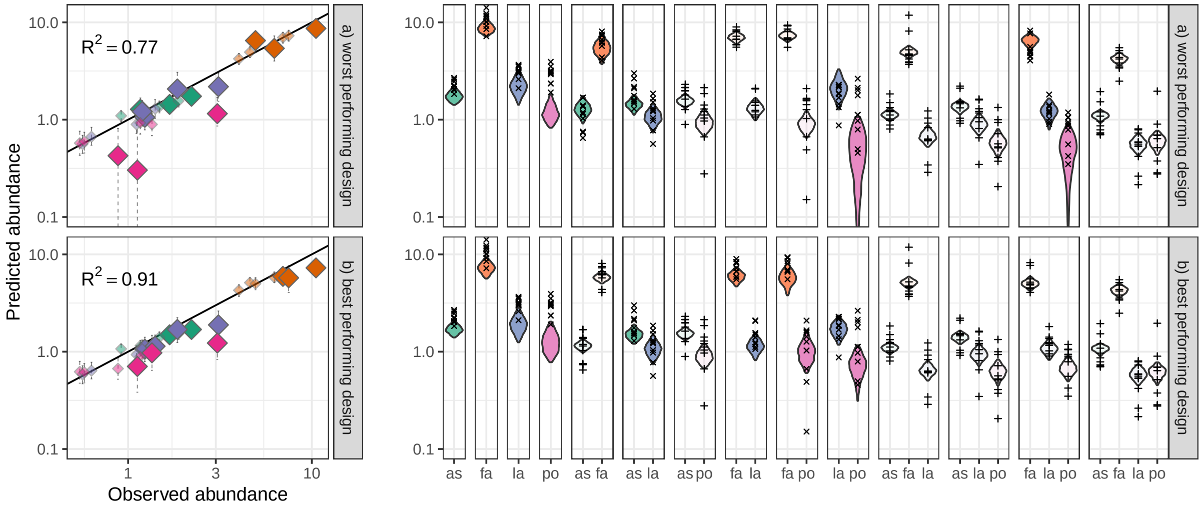

Supplementary Figure 26: As Fig. 5 of the main text, but using the data for the non-native plants community.

initial assemblages that do not contain previously observed endpoints as a subset. Additionally, the first set of experiments might yield sufficient constraints to estimate some subset of the coefficients of $B$, and this partial model could be used to target unobserved endpoints that have a high probability of feasibility or coexistence. Alternatively, one could estimate a "draft" of the full $B$ matrix using the incomplete data (for instance, with some regularization) for the same purpose.

These approaches could be iterated several times, updating the design after each experiment to maximize the probability that the next experiment is informative. Devising iterative schemes that are optimal, in theory or in practice, remains an intriguing open problem. We simply note that this kind of principled, "on-line" updating of experimental designs holds significant promise for navigating the enormous space of combinations faced when experimenting with a large species pool.

Finally, in cases where the coexistence of many species is rare, an iterative search beginning with small assemblages may be a fruitful way to assemble much larger ones. Rather than trying many combinations of large numbers of species in the hope of hitting on the rare case of coexistence, one could begin to infer the coefficients of $B$ from small communities, and gradually target larger coexisting communities, updating $B$ along the way. This approach can be used to find large sets of species that coexist stably and repeatably, for use in other experiments and applications.
